# Subcellular localization of drug distribution by super-resolution ion beam imaging

**DOI:** 10.1101/557603

**Authors:** Xavier Rovira-Clave, Sizun Jiang, Yunhao Bai, Graham Barlow, Salil Bhate, Ahmet F. Coskun, Guojun Han, Bokai Zhu, Chin-Min Kimmy Ho, Chuck Hitzman, Shih-Yu Chen, Felice-Alessio Bava, Garry P. Nolan

## Abstract

Technologies that visualize multiple biomolecules at the nanometer scale in cells will enable deeper understanding of biological processes that proceed at the molecular scale. Current fluorescence-based methods for microscopy are constrained by a combination of spatial resolution limitations, limited parameters per experiment, and detector systems for the wide variety of biomolecules found in cells. We present here super-resolution ion beam imaging (srIBI), a secondary ion mass spectrometry approach capable of high-parameter imaging in 3D of targeted biological entities and exogenously added small molecules. Uniquely, the atomic constituents of the biomolecules themselves can often be used in our system as the “tag”. We visualized the subcellular localization of the chemotherapy drug cisplatin simultaneously with localization of five other nuclear structures, with further carbon elemental mapping and secondary electron visualization, down to ∼30 nm lateral resolution. Cisplatin was preferentially enriched in nuclear speckles and excluded from closed-chromatin regions, indicative of a role for cisplatin in active regions of chromatin. These data highlight how multiplexed super-resolution techniques, such as srIBI, will enable studies of biomolecule distributions in biologically relevant subcellular microenvironments.

**One Sentence Summary:** Three-dimensional multiplexed mass spectrometry-based imaging revealed the subcellular localization of proteins and small molecules at super-resolution.

## Main Text

Form drives function in cells and tissues. This is especially true when researchers attempt deciphering of the complex interplay of molecular components that drive the biology of cells. Currently, biochemical approaches decipher meaning and mechanism from inferential experiments that allow researchers to abstract a picture of how the molecular machines of a cell function. This begs the question—would it not be more useful to image these machines in their natural contexts? Fluorescence microscopy is currently the de-facto choice for biomolecular imaging, although multiplexing is often challenging due to the spectral overlap of fluorophores. Recent advancements partially overcome this limitation by using cyclic-hybridization protocols to enable multiparameter visualization of proteins, RNA, and DNA (*1-4*). These techniques allow a range of studies from single-cells at super-resolution (*5*) to whole tissues (*6*) and have revealed a rich interplay between nucleic acids (*5, 7, 8*). There remain challenges however, such as acquisition time for imaging and sample positional shifts between cycles, which become more pronounced at super-resolution scales (*9*). In addition, imaging of small molecules, such as drugs, metabolites, or lipids, is also complicated since tags, such as fluorophores conjugated to biomolecules, can affect biological activities (*10*). The ability to directly image atomic components as labels would, with higher resolution, allow for a three-dimensional (3D) reconstruction of the distribution of multiple biomolecules in whole cells. This will complement the tissue mapping efforts now ongoing world-wide by bringing subcellular maps of the 3D nucleome and proteome to more public use and application.

An alternative modality of biomolecular imaging is mass spectrometry imaging (MSI). Images are obtained via the rastering of a biological sample with a laser (*11*) or an ion beam (*12*) to generate free molecules or ions that are analyzed by a mass spectrometer. We have previously demonstrated the ability of secondary ion beam imaging (SIMS), a form of MSI, to image multiple targeted-proteins in tissue samples using a technique named multiplexed ion beam imaging (MIBI) (*13*). In MIBI, proteins of interest are stained with isotope-tagged antibodies, and the sample is rasterized with an oxygen-based primary ion beam to generate a two-dimensional composite image of up to 50 proteins down to 260 nm resolution (*14*). In a similar approach referred to as imaging mass cytometry (IMC), protein and RNA can be visualized in tissues using laser ablation at 1000 nm resolution (*15, 16*). Although MIBI can define subcellular distribution of dozens of biomolecules, resolution limits (defined by the ion gun, vibration effects, and the working distance from the gun to the sample) are currently the major handicap for 3D cellular imaging at the nanoscale. Instrument development in the SIMS field has yielded devices that enable 3D visualization of small molecules with subcellular resolution suggesting the potential for multiplexed analysis of dozens of biomolecules and small molecules at the nanoscale (*17-23*). However, there is currently no method that allows specific targeting of multiple biomolecules via SIMS at super-resolution.

Here, we report the development and application of super-resolution MIBI using a positively charged cesium primary ion beam that allows visualization of subcellular structures with lateral (XY) and axial (Z) resolutions down to approximately 30 nm and 5 nm, respectively. We highlight the development of a toolkit for specific protein imaging with this approach. This method, which we call super-resolution ion beam imaging (srIBI), is capable of determining the precise subcellular locations of multiple small molecules, proteins, and/or nucleic acids at single-molecule resolution. We demonstrate the capabilities of srIBI by simultaneously imaging five distinct subnuclear structures and the chemotherapeutic drug cisplatin. Cisplatin is enriched in nuclear speckles, suggesting that this drug influences pre-mRNA processing. Utilizing a framework incorporating dimensional reduction and clustering methods to analyze srIBI data, we identify microenvironments within the nucleus, termed nuclear neighborhoods. Studying the interaction of these nuclear neighborhoods, our results suggest a directionality of cisplatin action within nuclear speckles. srIBI joins the growing suite of techniques that image molecular component in situ. Use of these approaches will allow study of the roles of small molecules in biological processes and the functions of distinct biomolecules in multi-component molecular pathways.

## Results

### Overview of srIBI

srIBI uses a positively charged cesium primary ion beam with a small spot size to obtain 3D super-resolution multiparametric visualization of cellular features (**Fig. 1A**), including the distribution of small molecules. Cells are cultured on a conductive substrate and treated with a small molecule that carries a stable isotope that can be efficiently ionized by the cesium beam. Cells are then simultaneously stained with multiple isotope-tagged antibodies using a protocol optimized for intracellular staining, dried under vacuum, and loaded into the instrument. A cell of interest is identified with an optical camera and then iteratively rasterized with the cesium primary ion beam. This process releases a cloud of negative secondary ions at the point of contact on the cell surface. The liberated secondary ions are collected and recorded, pixel-by-pixel, using a magnetic sector mass spectrometer. This results in a two-dimensional (2D) image for each analyzed isotope. Serial acquisition of hundreds of planes yields a stack of 2D images for each isotope that can be merged to obtain a multiparameter visualization of the cellular features of interest and volumetrically reconstructed to provide a 3D composite image of the analyzed cell at super-resolution.

**Figure 1.**
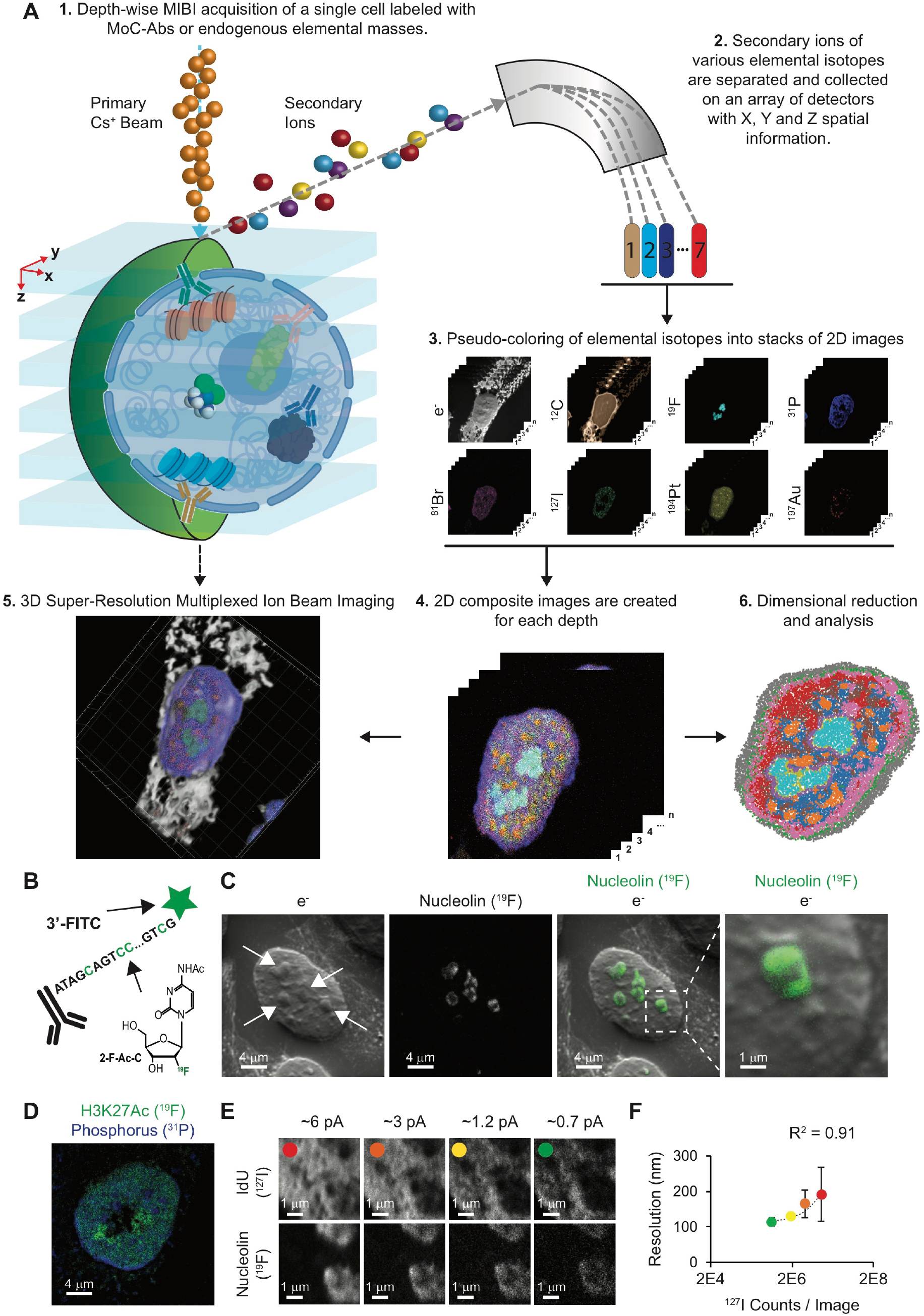
Super-resolution visualization of nuclear structures using srIBI. (**A**) Workflow of super-resolution ion beam imaging (srIBI). (1) Cells are treated with the drug of study, fixed, and stained with MoC-Abs. Endogenous elemental masses can also be detected. (2) Cells are rasterized by a cesium primary ion beam, and the secondary ions are collected by a magnetic sector mass spectrometer. (3) Spatial information is serially recorded in up to 8 channels simultaneously. (4) A composite image of the cell is reconstructed by combining and pseudocoloring the total number of channels at each depth. (5) srIBI yields a 3D rendering based on the depth profile that reveals the specific sub-cellular localization of endogenous and labeled targets.(6) Ion count extraction on a pixel-by-pixel basis and application of dimensional reduction methods identifies nuclear neighborhoods for feature extraction. (**B**) Schematic of ^19^F-based MoC-Ab. The oligonucleotide includes a FITC fluorophore at the 3’ end and the modified base 2-F-Ac-C (green) in place of deoxycytidine. (**C**) Representative ^−^e and ^19^F images of a HeLa cell stained with anti-nucleolin-^19^F/FITC. From left to right: (1) ^−^e image of a HeLa cell; nucleoli are indicated with white arrows. (2) Image based on ^19^F signals originating from anti-nucleolin-^19^F/FITC. (3) Overlay of images showing the colocalization of ^19^F signal with the subnuclear structures predicted to be the nucleoli from the ^−^e image. (4) An enlarged image of a nucleolus. (**D**) Representative srIBI image of a HeLa cell stained with anti-H3K27Ac-^19^F/FITC. Overlay of ion images for ^19^F (green) and ^31^P (blue). The image is the sum of 10 consecutive planes. (**E**) HeLa cells were labeled for 24 hours with IdU (^127^I) and stained with anti-nucleolin-^19^F/FITC. Using srIBI, 40 depths of a single cell were acquired with an increase in the current every 10 scans. Images of newly synthesized DNA (top) and nucleolin (middle) and their overlay (bottom) show details of a region within the nucleus, at different currents. Details of DNA and nucleolus in these settings are increasing by lowering the current, at the cost of ion yield. Colored dots indicate the image used for the quantification shown in panel F. (**F**) Quantification of the relationship between srIBI resolution and secondary ion yield. The lateral resolution of ^127^I (present in IdU) at different currents from panel E was calculated using the 84% to 16% criterion. The R^2^ for the relationship between the increase in resolution and decrease in ion yield was 0.91. See Fig. S5 for details.

### Design and validation of novel antibody tagging protocol for srIBI

In our original MIBI approach, an oxygen duoplasmatron source was used to generate 2D images of tissue sections stained with lanthanide-tagged antibodies down to 260 nm resolution (*13*). Attaining resolutions beyond this limit requires a more finely focused beam. The cesium primary ion beam can achieve resolutions down to 30 nm in biological samples (*17, 24*). However, lanthanide-tagged antibodies cannot be used since lanthanides are not efficiently ionized with a cesium ion beam (*25*). To circumvent this issue, we developed a new labeling strategy using isotopes that are efficiently ionized with a cesium beam. DNA is a versatile polymer that has been extensively functionalized for imaging purposes (*3, 26-28*). We reasoned that antibodies conjugated with single-stranded DNA (ssDNA) oligonucleotides carrying stable isotopes that are present in low amounts in cells would enable super-resolution protein visualization by MIBI. To this end, we developed a tool we call a mass-oligonucleotide-conjugated antibody (MoC-Ab). Isotope-modified ssDNA oligonucleotides were conjugated to antibodies through thiol-maleimide click chemistry (**Fig. S1**) (*29*). MoC-Abs can be used for intracellular protein staining in an optimized protocol that involves fixation with 1.6% paraformaldehyde, permeabilization, and blocking with a high salt concentration in the presence of sheared salmon sperm DNA to avoid non-specific binding (**Fig. S2**).

In a proof-of-concept experiment, the single stable fluorine isotope (^19^F) was incorporated by chemical synthesis into a FITC-labeled ssDNA by substituting all cytidines with the commercially available N4-acetyl-2’-fluoro-2’-deoxycytidine (2-F-Ac-C) (**Fig. 1B;** ^19^F/FITC MoC-Ab). The behavior of MoC-Abs is similar to that of unmodified antibodies as shown by confocal microscopy analyses of target specificity, staining intensity, and background levels of nucleolin staining in HeLa cells (**Fig. S3A-C**). We next assessed the performance of MoC-Abs in srIBI by visualizing HeLa cells stained with anti-nucleolin-^19^F/FITC. Rastering a HeLa cell with the cesium ion beam revealed an enriched ^19^F signal inside the nucleus akin to the fluorescence signal observed using confocal microscopy (**Fig. S3D**, right images). Moreover, control experiments without anti-nucleolin-^19^F/FITC resulted in absence of ^19^F signal (**Fig. S3D**, left images), suggesting that the ^19^F signal observed in the stained cell resulted from the MoC-Ab. The high ^31^P content of the DNA phosphate backbone enabled visualization of the nucleus in the srIBI experiment in a manner analogous to use of DAPI in conventional fluorescent microscopy (**Fig. S4**).

In srIBI, secondary electron (^-^e) and elemental ion information are simultaneously recorded for each plane. The ^−^e image, akin to that from a scanning electron microscope, enables the identification of certain subnuclear structures such as the nucleolus (**Fig. 1C**, left image). To confirm the specificity of anti-nucleolin-^19^F/FITC, we merged the ^19^F and ^-^e signals and observed a specific enrichment of the ^19^F signal in nucleolar structures identified through the ^−^e image (**Fig. 1C**, overlay and right image). To demonstrate the versatility of srIBI, we labeled transcriptionally active chromatin using a ^19^F/FITC MoC-Ab that binds to H3K27Ac (**Fig. 1D**). The ^19^F signal overlapped with nuclear regions low in ^31^P confirming the specificity of anti-H3K27Ac-^19^F/FITC (**Fig. S5**). These results demonstrate that detection of MoC-Abs labeled with ^19^F by ion beam imaging can reveal subnuclear structures of known provenance.

In SIMS, the beam size dictates the lateral resolution and ion yield of the acquired image (*30*). The size of the beam is proportional to its current. We quantified the effect of beam current on the ion count per pixel in our samples by acquiring sequential images of the nucleolus of a HeLa cell. Raster size, pixel dwell time, and number of pixels acquired were maintained at constant levels, and the beam current was varied (**Fig. S6A and B**). We observed that the increase in current scaled with ion counts per pixel (**Fig. S6C and D**) at the expense of resolution (**Fig. S6B**). To quantify the lateral resolution, we acquired sequential images of a HeLa cell incubated for 24 hours with 5-iodo-2’-deoxyuridine (IdU) to specifically label newly replicated DNA. In the condition tested, reduction of the beam current improved the lateral resolution from 193 nm to 113 nm and lowered the total ion count 16 fold (R^2^ = 0.91; **Fig. 1E and F** and **Fig. S6E-I**). These results show that srIBI resolves targeted subcellular structures with a resolution, to our knowledge, not attainable with current MSI methods that target specific proteins (*13, 15*).

### Multiparametric imaging and unsupervised identification of subnuclear structures

We leveraged the capabilities of mass spectrometry for simultaneous data acquisition by expanding the number of elements used to label MoC-Abs. Bromine (^79/81^Br) and iodine (^127^I) are two elements used to produce synthetic nucleotides classically harnessed to study cell proliferation, DNA replication, and RNA synthesis. We synthesized and validated MoC-Abs containing 5-bromo-2’-deoxycytidine (5-Br-dC) and 5-iodo-2’-deoxycytidine (5-I-dC) for the labeling of centromeres (anti-CENP-A-^81^Br/Cy3) (**Fig. 2A** and **Fig. S7**) and transcriptionally active chromatin (anti-H3K27Ac-^127^I/Cy5) (**Fig. 2B** and **Fig. S8**), respectively.

**Figure 2.**
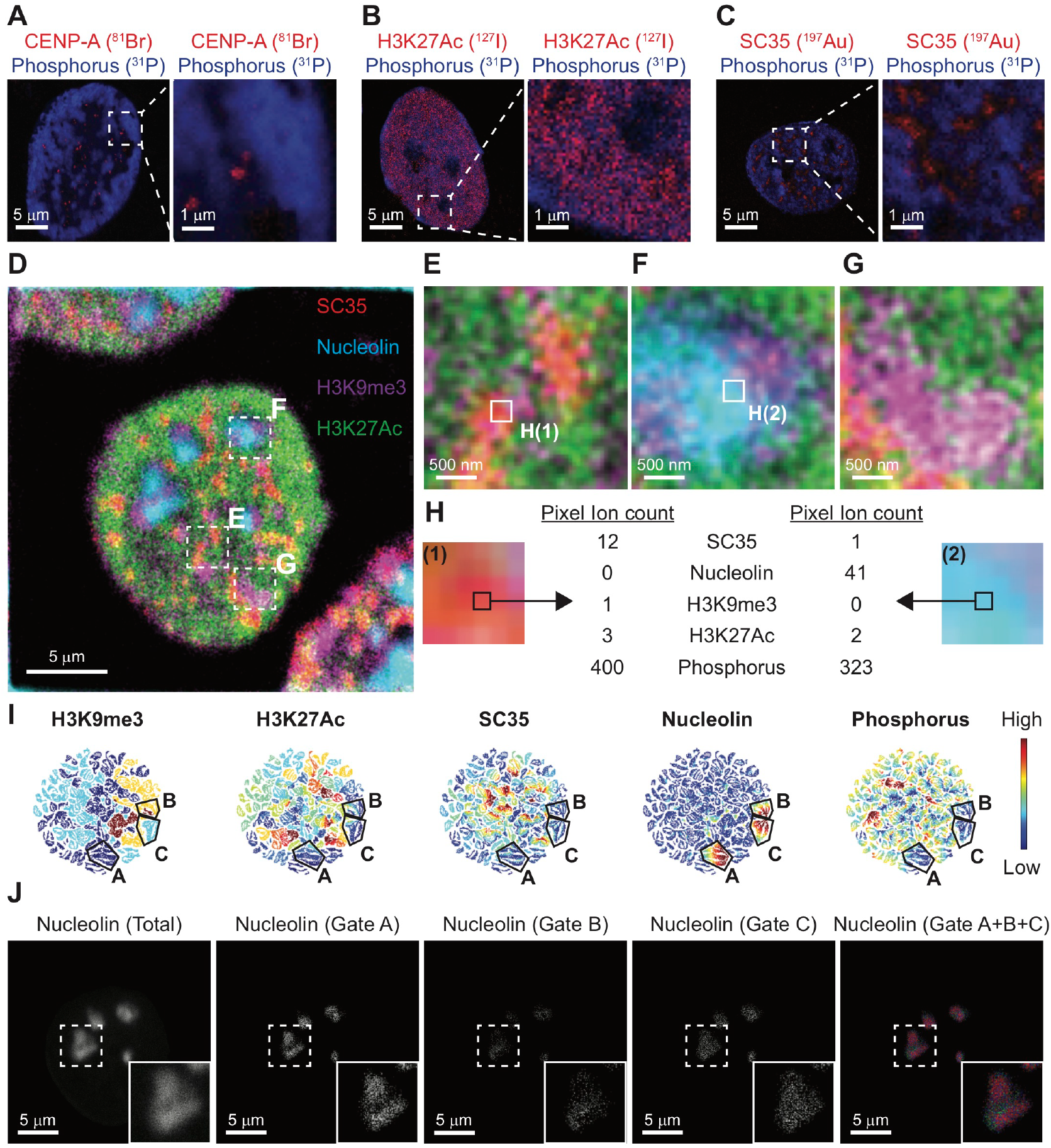
Multiparametric srIBI using MoC-Abs identifies unique subcellular features. (**A-C**) (Left) Representative srIBI images of a HeLa cell stained with **A**) anti-CENP-A-^81^Br/Cy3, **B**) anti-H3K27Ac-^127^Ir/Cy5, and **C**) anti-SC35-biotin recognized by streptavidin-^197^Au/FITC. Overlay of ion images for MoC-Ab (red) and phosphorus (^31^P; blue). (Right) A digital zoom of boxed area in the original image to show specificity of MoC-Ab signal. All images consist of the sums of 10 consecutive planes. (**D**) Composite srIBI image of nucleolin (cyan), H3K9me3 (magenta), H3K27Ac (green), and SC35 (red) in a HeLa cell. Cells were stained with anti-nucleolin-^19^F/FITC, anti-H3K9me3- ^81^Br/Cy3, anti-H3K27Ac-^127^Ir/Cy5, and anti-SC35-biotin (recognized by streptavidin- ^197^Au/FITC). The image consists of the sum of 10 consecutive planes. (**E**-**G**) Multiple enlarged images from boxed regions in panel D showing **E**) a nuclear speckle, **F**) a nucleolus, and **G**) heterochromatin. (**H**) The largest nucleus in panel D was masked using the phosphorus signal. The ion counts for each acquired marker (nucleolin, ^19^F; DNA, ^31^P; H3K9me3, ^81^Br; H3K27Ac, ^127^I; and SC35, ^197^Au) were extracted. (**I**) viSNE maps of the nucleus in the center of the image shown in panel D were created using the ion counts for each of the 84836 pixels from the mask. The ion count for each marker in each pixel was min-max normalized to scale the range to [0, 1]. Each point represents a pixel. Pixels grouped in distinct regions based on the expression of each marker. The color in each map represents the intensity of the indicated marker in each pixel. The areas marked A-C indicate three distinct groups of nucleolin-positive pixels identified manually in the nucleolin map. (**J**) Grouped pixels from the unsupervised viSNE map are differentially distributed in space as shown in srIBI images of (1) total nucleolin (^19^F) from the HeLa cell shown in panel D, (2) ^19^F signals from pixels within gate A, (3) gate B, (4) gate C, and (5) overlay of ^19^F signals from gates A (red), B (green), and C (blue).

In addition to nucleotide-based labeling methods, other reagents can be harnessed for srIBI (*31, 32*). We stained HeLa cells with phalloidin conjugated to ATTO514, a fluorophore that contains six ^19^F atoms (**Fig. S9A**). The distribution of actin filaments observed in HeLa cells labeled with this reagent was comparable in confocal microscopy and srIBI images (**Fig. S9B and C**). Nuclear speckles are nuclear domains containing inter-chromatin material enriched in pre-mRNA splicing components (*33*). We validated detection of SC35, a nuclear speckle protein, using anti-SC35-biotin with streptavidin-labeled 1.4 nm gold nanoparticles (streptavidin-^197^Au/FITC) by srIBI (**Fig. 2C** and **Fig. S10**). To further demonstrate the specificity of antibody-based tools for srIBI, we labeled transcriptionally silent chromatin (anti-H3K9me3-^81^Br/Cy3) and nucleoli (antinucleolin-^127^I/Cy5 and anti-NPM1-biotin recognized by streptavidin-^197^Au/FITC) in HeLa cells, obtaining the expected patterns for all (**Fig. S11**).

We evaluated the crosstalk between channels by staining HeLa cells with individual MoC-Abs and assessing the signal in unstained channels. Minimal crosstalk between channels was observed (**Fig. S12**), therefore multiplexed imaging, in a manner akin to fluorescence microscopy, can be performed using srIBI (**Fig. S13**). To demonstrate the multi-parameter capabilities of srIBI, we simultaneously visualized nucleolin, H3K9me3, H3K27Ac, and SC35 in HeLa cells (**Fig. 2D**). Magnification of the composite image revealed a complexity not observed in single-channel images (**Fig. 2E-G** and **Fig. S14**). We extracted the ion count per channel on each pixel of the multiplexed image (**Fig. 2H**) and used viSNE to identify subnuclear structures (**Fig. 2I**) as previously described (*34*). Pixels with high expression of nucleolin were grouped in three distinct regions that had variable levels of H3K9me3 and low levels of SC35, H3K27Ac, and phosphorus (**Fig. 2I**; gates A to C).

To gain insights into nucleolar organization, we categorized the pixels within nucleolin-rich regions in the three distinct subsets and generated a composite image of spatial distribution (**Fig. 2J**). There was a different localization pattern of pixels with high H3K9me3 signal within the nucleolin-rich regions (**Fig. 2J**, gate B) than in the regions with low amounts of H3K9me3 (**Fig. 2J**, gate A); these patterns were consistent in all nucleoli of this cell (**Fig. 2J**, gate A+B+C). Thus, srIBI can be used to visualize multiple isotopes simultaneously, and the application of dimensionality reduction techniques to the data enables identification of distinct subnuclear structures.

### 3D reconstruction at the nanoscale with srIBI

In srIBI, ion beam strength and residence time at the spot can yield secondary ion information for each plane that results from the ablation of a just a few nanometers of the sample surface, and consecutive planes have similar ion counts (**Fig. S15**). We reasoned that whole 3D nuclear reconstruction could be achieved with srIBI if data were acquired on many consecutive planes. Previous axial resolution quantifications in biological samples on the nanoSIMS was determined to be around 5 nm (*35*). In agreement with this measurement, reconstructions of centromeres from srIBI (**Fig. S16**) resembled those identified by recent super-resolution microscopy experiments (*36*).

To demonstrate the feasibility of whole organelle 3D reconstruction, we next acquired 785 planes of a single nucleolus (**Fig. 3A** and **Fig. S17**) and performed a volumetric reconstruction (**Fig. 3B** and **Movies S1 and S2**). srIBI is capable of resolving juxtaposed subnuclear structures as a 50-plane acquisition was sufficient to distinguish between the centromere, nucleolus, and nuclear speckles in a HeLa cell (**Fig. 3C**). As further validation, we performed 3D rendering of 400 planes from a single HeLa cell labeled with anti-nucleolin-^19^F, anti-H3K9me3-^81^Br, anti-H3K27Ac-^127^I, and anti-SC35 (recognized by streptavidin-^197^Au), creating a composite 3D visualization of the nucleus (**Fig. 3D** and **Movie S3**). These results indicate that srIBI enables whole-cell reconstruction of the genome and epigenome. Combined with an analytical framework, as shown below, srIBI will allow intricate dissection of subcellular microenvironments in single cells.

**Figure 3.**
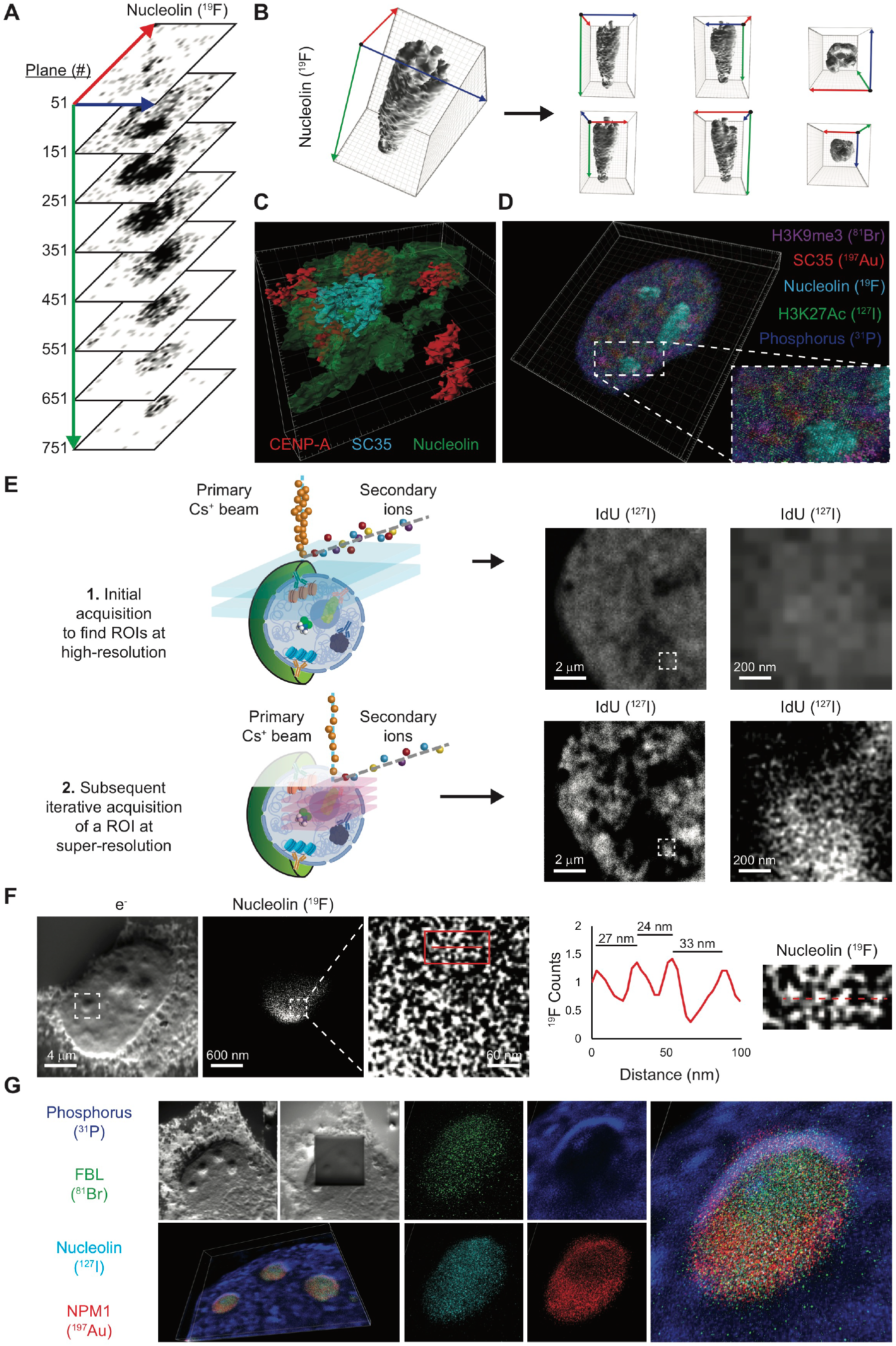
3D nanoscale imaging of the nucleus through iterative srIBI. (**A**) Representative single-plane srIBI images at different depths of a HeLa cell nucleolus. HeLa cells were stained with anti-nucleolin-^19^F/FITC, and 785 individual planes were acquired to obtain srIBI images of a nucleolus from its appearance to its disappearance. Single planes every 100 depths show a distinctive molecular distribution of nucleolin in the 3D space. See Fig. S15 for images of each individual plane. Blue, red, and green arrows indicate x-axis, y-axis, and z-axis, respectively. (**B**) (Left) 3D surface reconstruction of images of nucleolin staining of a nucleolus shown in panel A. (Right) Overviews of the same nucleolus along x-axis (blue arrow), y-axis (red arrow), and z-axis (green arrow) with the origin represented as a black dot. (**C**) Representative 3D surface reconstruction of nucleolus (green), centromeres (red), and nuclear speckles (cyan). HeLa cells were stained with anti-nucleolin-^19^F/FITC, anti-CENP-A-^81^Br/Cy3, and anti-SC35-biotin (recognized by streptavidin-^197^Au/FITC). The image consists of the 3D reconstruction of a stack of 40 consecutive planes. (**D**) Representative 3D reconstruction of nucleolin (cyan), phosphorus (blue), H3K9me3 (magenta), H3K27Ac (green), and SC35 (red) in a HeLa cell stained with anti-nucleolin-^19^F/FITC, anti-H3K9me3-^81^Br/Cy3, anti-H3K27Ac-^127^I/Cy5, and anti-SC35-biotin (recognized by streptavidin-^197^Au/FITC). The image consists of the 3D reconstruction of a stack of 400 consecutive planes. Enlarged images from the boxed region show details of marker distribution. (**E**) (Left) Workflow of iterative srIBI: (1) Five to ten depths at high current are acquired in a cell of interest at 25 * 25 µm to identify a ROI. (2) Iterative acquisition is performed by focusing the beam into the ROI at lower currents in a smaller area of 5 * 5 µm or 10 * 10 µm. (Right) (Top) Representative region of the IdU signal (^127^I) in a nucleus of a HeLa cell labeled for 24 hours with IdU (^127^I). (Bottom) A 10 * 10 µm ROI was acquired for super-resolution imaging of chromatin folding. Enlarged image of the boxed region shows fine detail of IdU labeled chromatin. (**F**) Quantification of the resolution of iterative srIBI imaging of a nucleolus. From left to right: ^−^e image of a HeLa cell reveals subnuclear structures including the nucleoli chosen for iterative srIBI. (2) A 3 * 3 µm ROI was acquired by iterative srIBI for nucleolin (^19^F). (3) An enlarged view from the boxed region in image 2 showing the nanoscale organization of nucleolin (^19^F). (4) Line scan along the red line in the boxed region in image 3 demonstrates identification of molecules spaced about 30 nm. (5) An enlarged view of the boxed region in image 3. See Fig. S16 for additional examples. (**G**) Iterative srIBI 3D reconstruction. HeLa cells were stained with anti-FBL-^81^Br/Cy3, antinucleolin-^127^Ir/Cy5, and anti-NPM1-biotin (detected with streptavidin-^197^Au/FITC). Iterative srIBI was performed on a site with three nucleolin. The ^-^e image confirmed that the ROI was acquired. Nucleolin (cyan), phosphorus (blue), FBL (green), and NPM1 (red) were used for 3D reconstruction from 40 consecutive planes. These images identify the granular component (GC, NPM1-positive), dense fibrillar component (FBL-and nucleolin-positive), and perinucleolar heterochromatin (PNC, phosphorus-high).

### Iterative srIBI resolves nanoscale structures

A lateral resolution down to 30 nm in biological samples has been previously reported using the cesium primary ion beam by focusing it to an area of interest measuring a few microns (*17, 24*). During each pass of the primary ion beam only the top material contacted is ablated. The majority of the sample is preserved due to the high axial resolution and thus is available for further imaging. We reasoned that we could perform an initial acquisition of a target cell to define a region of interest (ROI) and subsequently focus on that sub-cellular feature for higher resolution profiling (**Fig. 3E**, workflow). Indeed, re-probing a region of interest in an IdU-treated cell enabled iterative imaging at higher resolution (**Fig. 3E**, IdU images).

To achieve our highest benchmarked resolution in MSI of specific proteins, we performed iterative srIBI in a single HeLa cell labeled with anti-nucleolin-^19^F. Using line scans across the image to quantify the distance between ^19^F signals as commonly done for fluorescence-based super-resolution images (*37*), we identified nucleolin molecules spaced at about 30 nm apart (**Fig. 3F** and **Fig. S18**). The secondary electron image of a cell before and after an iterative srIBI acquisition showed a generally even ablation of the field of view, which focused on three nucleoli (**Fig. 3G**, e^-^image). A 3D reconstruction of a single nucleolus from this acquisition (**Fig. 3G**) revealed the expected spatial distribution of nucleolar proteins within a single nucleolus (**Fig. 3G** and **Fig. S19**). Hence, a first scan to define an area of interest and subsequent imaging to enhance the resolution revealed molecules spaced about 30 nm apart.

### Differential drug distribution in nuclear neighborhoods

SIMS enables subcellular visualization of exogenously incorporated small molecules and has been previously used to study the cellular distribution of the metallo-drug cisplatin (*38*), which is used to treat various types of cancer (*39*). The platinum atom of cisplatin (**Fig. S20A**) can be readily ionized using the cesium beam. We incubated TYK-nu cells, which are immortalized ovarian cancer cells, for 24 hours with a range of concentrations of cisplatin to identify a concentration at which drug was internalized but caused minimal cell death (**Fig. S20B**). Naturally occurring platinum is composed of five stable isotopes (**Fig. S20C**); we took advantage of isotopically pure cisplatin (^194^Pt) to maximize Pt counts. Cells treated with 5 µM cisplatin for 24 hours exhibited high ^194^Pt counts (**Fig. S20D**), and signal was observed in cytoplasm and nuclei (**Fig. S20E**), consistent with published results (*38*). The crosstalk between ^194^Pt and the channels used for srIBI were minimal (**Fig. S20F**). We performed MoC-Ab staining of cisplatin-treated cells and acquired images of ^−^e, ^12^C (indicative of overall cellular structure), ^19^F (nucleolin), ^31^P (DNA), ^81^Br (H3K9me3), ^127^I (H3K27Ac), ^194^Pt (cisplatin), and ^197^Au (SC35) to generate an eight-channel image at super-resolution. srIBI revealed a non-uniform distribution of cisplatin in TYK-nu nuclei (**Fig. 4A**) and composite images enabled the visualization of the subnuclear distribution of cisplatin (**Fig. 4B**).

**Figure 4.**
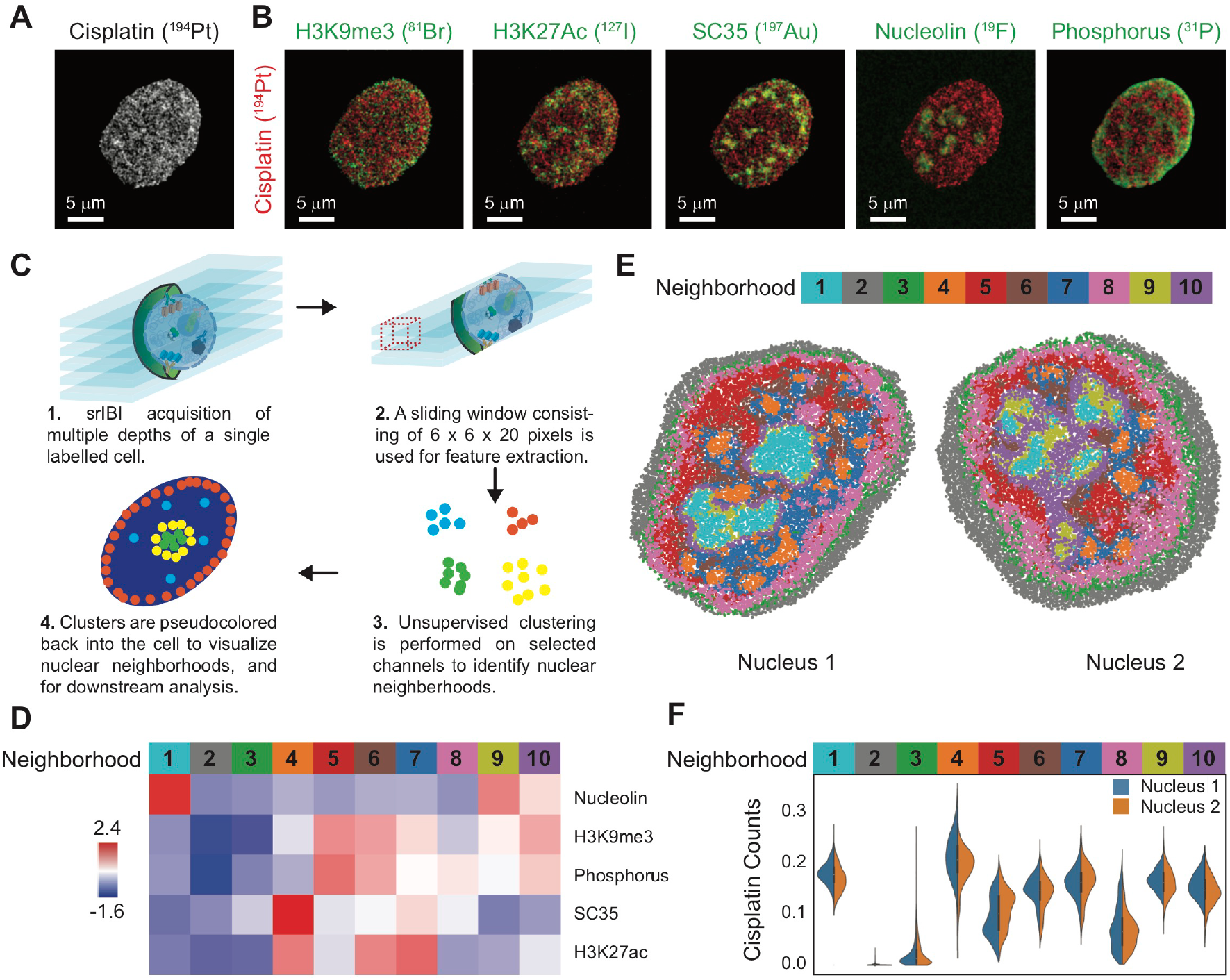
Subcellular localization of cisplatin distribution. (**A**-**B**) TYK-nu ovarian cancer cells were treated with 5 µM cisplatin for 24 hours and stained with anti-nucleolin-^19^F/FITC, anti-H3K9me3-^81^Br/Cy3, anti-H3K27Ac-^127^I/Cy5, and anti-SC35-biotin (detected with streptavidin-^197^Au/FITC). We simultaneously acquired images of nucleolin (^19^F), DNA (^31^P), H3K9me3 (^81^Br), H3K27Ac (^127^I), cisplatin (^194^Pt), and SC35 (^197^Au), in addition to ^−^ e. (**A**) Representative srIBI image of cisplatin (^194^Pt) distribution within the cell. (**B**) Overlay of ion images for cisplatin (^194^Pt; red) and a second marker (green). All images consist of the sums of 10 consecutive planes. (**C**) A schematic for the identification of nuclear neighborhoods. (1) SrIBI acquisition is performed on single TYK-nu cells treated with cisplatin and labeled with MoC-Abs as described in panel A.(2) A sliding window of 6 * 6 * 20 pixels is used for feature extraction for all channels imaged.(3) Unsupervised clustering is performed on extracted features to identify nuclear neighborhoods.(4) Identified clusters are recolored back onto the cell to visualize nuclear neighborhoods spatially. (D) A heatmap of the 10 distinctive neighborhoods identified based on the five indicated nuclear markers. The intensity of each marker is denoted by the scale bar on the left. The identified clusters of interactions, termed nuclear neighborhoods, are colored and numbered. (**E**) Identified nuclear neighborhoods are used to recreate the cell, resulting in distinctive structures that resemble known features in the nucleus. Two representative TYK-nu nuclei are shown. (**F**) The cisplatin counts are plotted for each nuclear neighborhood identified in panel D for the two nuclei shown in panel E.

Biomolecules in the nucleus are organized spatially to form pockets of interaction (*40, 41*), akin to tissue microenvironments (*3, 14, 42*). In such a nuclear neighborhood, the function of various biomolecules are dependent on their spatial localization and the context of other nearby constituents. The multiplexed data generated from srIBI allowed us to test the hypothesis that biomolecules in the nucleus have context-dependent functions. Whole cell srIBI acquisition was performed on cisplatin-treated cells (**Fig. 4C**, top left, and **Movie S4**). We applied a sliding window, consisting of 6 * 6 * 20 pixels, to extract the average isotope counts per window for five channel (H3K9me3, H3K27Ac, SC35, nucleolin and phosphorus; **Fig. 4C**, top right). Unsupervised clustering was then performed on these extracted features to group windows based on similarity (**Fig. 4C**, bottom right). Individual voxels were then pseudo-colored based on these groups for visualization purposes (**Fig. 4C**, bottom left). In total, 20,000 randomly sampled 3D voxels from the middle 40 Z-planes were extracted from each of the 2 cells, for a sum of 40,000 voxels.

Clustering of these 40,000 voxels based on the five channels revealed 10 distinct clusters, which represented 10 different nuclear neighborhoods (**Fig. 4D**) at this scale of observation. We used the histone markers H3K9me3 and H3K27Ac as well-characterized markers for heterochromatin (higher DNA density) and euchromatin (low DNA density), respectively (*43*). That these markers were located in different clusters is supportive of the technique’s capability (**Fig. 4D**, Neighborhoods 5, 7, and 10). We observed regions that are high in H3K9me3 also have abundant phosphorus, as expected because of the condensed nature of heterochromatin (**Fig. 4D**, Neighborhoods 5). We then colored each nucleus based on these groups and projected the 3D voxels onto a 2D plane to better visualize the spatial distribution of the nuclear neighborhoods (**Fig. 4E**). These images reveal distinctive nuclear structures, such as nucleolin-like structures (Neighborhood 1), nuclear speckles (Neighborhood 4), and chromatin that resembles lamin-associated domains (Neighborhood 8). Features that are difficult to distinguish visually in the srIBI images (**Fig. 4B**) could now be appreciated (**Fig. 4E**), such as peri nucleolar heterochromatin (Neighborhood 9), as well as heterochromatin-like regions (Neighborhood 10), and euchromatin (Neighborhoods 6 and 7). This is also able to distinguish features that appear to be edge effects (Neighborhoods 2 and 3).

Identification of subcellular microenvironments, combined with the spatial characterization of atomic components from small molecules, offers the ability to create subcellular maps for multiple types of biomolecules. To identify the distribution of cisplatin in the nucleus, we set out to quantify cisplatin levels in each nuclear neighborhood. We observed that nuclear speckles contained high levels of cisplatin (**Fig. 4F**, Neighborhood 4). Cisplatin was depleted from heterochromatin regions and lamin-associated domains (**Fig. 4F**, Neighborhoods 5 and 8). To obtain a super-resolution image of cisplatin localization we used iterative srIBI, to reveal at an unprecedented level, the spatial enrichment of cisplatin along the nuclear speckles and their depletion from closed chromatin (**Fig. 5A** and **Fig. S21**). These results demonstrate a differential distribution of cisplatin in distinct nuclear neighborhoods, suggestive of different mechanisms of action.

**Figure 5.**
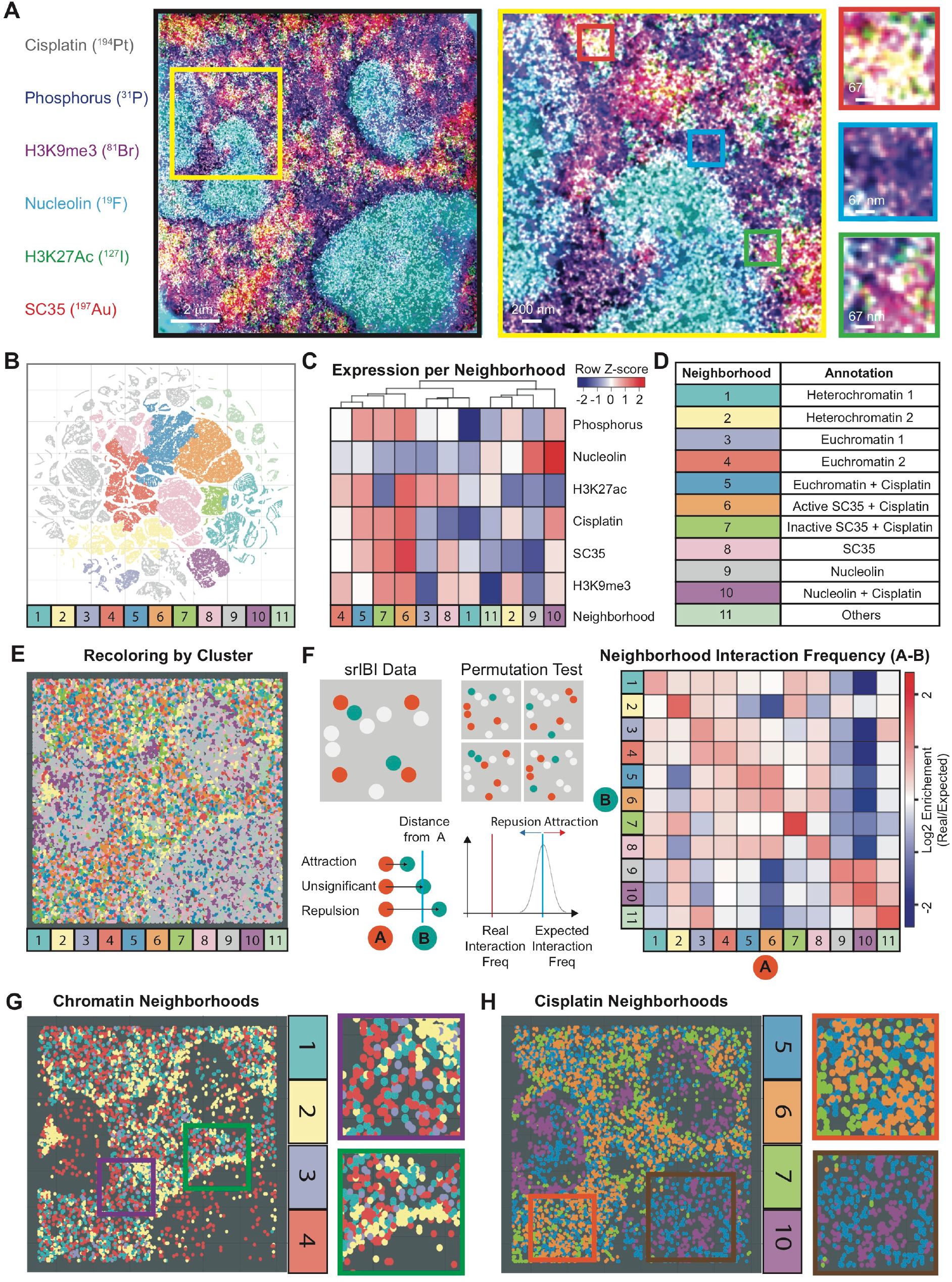
An analytical framework to identify nuclear neighborhood interactions in multiplexed super-resolved data. (**A**) Representative srIBI image of cisplatin-treated TYK-nu cells. TYK-nu ovarian cancer cells were treated with 5 µM cisplatin for 24 hours and stained with anti-nucleolin-^19^F/FITC, anti-H3K9me3-^81^Br/Cy3, anti-H3K27Ac-^127^I/Cy5, and anti-SC35-biotin (detected with streptavidin- ^197^Au/FITC). Images of nucleolin (^19^F), DNA (^31^P), H3K9me3 (^81^Br), H3K27Ac (^127^I), cisplatin (^194^Pt), and SC35 (^197^Au) were simultaneously acquired. A 100-µm^2^ROI in the nucleus was acquired by iterative srIBI. Displayed are nucleolin (cyan), phosphorus (blue), H3K9me3 (magenta), H3K27Ac (green), cisplatin (grey), and SC35 (red). Denoising was performed using a k nearest neighbor approach. An unfiltered image is shown in Fig. S21C. Enlarged images from boxed regions exemplify the nuclear organization diversity. Images consist of the sums of 20 consecutive planes. (**B**) t-SNE map colored by the 11 identified nuclear neighborhoods. The t-SNE map is derived from 100000 voxels of dimension (x, y, z) = (10, 10, 5) pixels (20000 voxels were randomly sampled across 5 different cells). Each point represents a voxel. Voxels grouped in distinct regions based on the expression of each marker. The 11 hierarchical clusters were identified by unsupervised hierarchically clustering, followed by manual annotation. (**C**) An expression profile of the mean marker expression in each of the 11 distinctive nuclear neighborhoods. The scale intensity of each marker is denoted by the color bar on the top right (Z-score, normalized to each row). (**D**) Annotation of each neighborhood, based on their mean marker expression profile from panel C. (**E**) Identified nuclear neighborhoods described in panel D are used to recreate the cell, resulting in distinctive structures that resemble known features in the nucleus shown in panel A. (**F**) Pairwise interaction frequency calculations. (Left) A graphical schematic of the permutation test implemented to quantify the frequency of pairwise neighborhood interactions is shown. If two points (e.g., A and B) were within 5 pixels of each other, they were defined as being “attracted”. Point labels were shuffled 1000 times, and the mean interaction frequency noted. The final distribution of shuffled interaction frequencies was plotted, and the mean was taken as the expected interaction frequency. Real interaction frequencies below the expected were indicative of repulsion between the points, whereas those above were attraction. (Right) The neighborhood interaction frequency map was calculated using the log2 enrichment of the real over expected number of pairwise interaction frequencies. (**G**-**H**) Nuclear neighborhoods are colored as shown in the legend of panel D to show **G**) chromatin-specific neighborhoods, and **H**) cisplatin-enriched neighborhoods. Enlarged images from boxed regions exemplify different neighborhood interactions: accessible euchromatin (purple box), heterochromatin near the nucleolus (green box), an inactive/active boundary of cisplatin containing nuclear speckles (orange box), and cisplatin-euchromatin adducts within the nucleolus (brown box).

### An analytical framework to identify nuclear neighborhood interactions in multiplexed super-resolved data

Diversity of nuclear organization is readily appreciated upon inspection of iterative srIBI images (**Fig. 5A**, right panels). To extract biological meaning from these images, we combined the visual nature of dimensionality reduction methods with the spatial information encoded within our data. We randomly selected 20000 voxels across five different cells for combined analysis. We then performed dimensional reduction using t-Distributed Stochastic Neighbor Embedding (t-SNE) on the log transformed, mean isotope counts per voxel (**Fig. S22A**). As expected, signals were grouped together based on levels of marker expression (**Fig. S22A**). We next applied unsupervised hierarchically clustering followed by manual annotation of the clusters to identify 11 unique nuclear neighborhoods (**Fig. 5B-D** and **Fig. S22B-D**). Each cell was relatively well represented in each neighborhood (**Fig. S22E**). We were able to reconstruct the nuclear organization diversity observed in iterative srIBI images of cisplatin treated cells (**Fig. 5A**) using the neighborhood and spatial information alone (**Fig. 5E** and **Fig. S23**), demonstrating that the inferred neighborhoods are an appropriate representation of nuclear organization.

If the nucleus is organized with intent, certain neighborhoods would interact with each other at higher frequency than random. To infer this, we devised and implemented a pairwise interaction permutation test (**Fig. 5F**, see Materials and Methods for details). In addition to testing for the frequency of interactions, we also tested for statistical significance for each pairwise interaction (**Fig. 5F** and **Fig. S24**). Certain nuclear neighborhoods tended to attract or repel each other. For example, neighborhoods rich in both SC35 and cisplatin repelled neighborhoods rich in nucleolin (**Fig. 5F**, Neighborhood 6 in A to Neighborhood 9 in B).

Focusing on neighborhoods representing chromatin (Neighborhoods 1 to 4), we observed that heterochromatin self-aggregated, was tightly packed, and surrounded the nucleolus (**Fig. 5G**, green box), whereas euchromatin was dispersed, as expected due to higher accessibility of the DNA (**Fig. 5G**, violet box). In line with previous reports, we corroborated that cisplatin accumulated in the nucleolus (**Fig. 5H**) (*38*) and also observed that cisplatin within the nucleolus spatially segregated into two distinct neighborhoods (**Fig. 5H**, brown box, Neighborhoods 5 and 10). In addition, there was a difference between two nuclear speckle neighborhoods enriched in cisplatin (**Fig. 5H**, red box, Neighborhoods 6 and 7). We observed that inactive SC35 neighborhoods (Neighborhood 7) tended to envelope the active SC35 neighborhoods (Neighborhood 6), suggesting directionality of cisplatin action (**Fig. 5H**, orange box and **Fig. S25**).

These results indicate that biologically relevant information can be obtained through the analysis of iterative srIBI images of subcellular entities.

## Discussion

srIBI is an MSI method enabled by a new set of reagents, MoC-Abs, that allow for the specific detection of proteins using MIBI coupled to a cesium primary ion beam. This method enables targeted super-resolution, 3D, multiplexed imaging of different biomolecule types in cells. srIBI detection of MoC-Abs yielded results akin to those obtained using standard confocal light microscopy but use of iterative srIBI imaging resulted in lateral resolutions of approximately 30 nm. Application of unsupervised dimensional reduction methods enabled the identification of subcellular structures and localization of the small-molecule drug cisplatin. As most drugs can be isotopically labeled with isotopes suc as ^13^C or ^15^N, srIBI will allow for study of localization of a variety of agents. Drug metabolism in and out of cells might also be determined by such imaging. Understanding the subcellular localization of drugs may maximize the success rates of drug candidates in clinical trials, holding promise for a more efficient and cost-effective drug discovery process (*44*).

Although srIBI has theoretical resolutions below the antibody size and potential for at least 50-plex imaging, it is currently hindered by low ion yields at high resolutions, the number of isotopes available for labeling, and the number of available detectors for higher multiplexing. Improvements to the ion yield can be linearly ameliorated through the attachment of more labeling isotopes per probe (which may result in slight loss of resolution). The MoC-Abs used in this work each contain 44 to 66 isotopic labels. An increase in labels per antibody can be readily achieved by increasing the number of substituted nucleotides, increasing the length of labeling oligonucleotide, or through nucleotide-based amplification methods (*28, 45-47*). Better estimations of the placement of the epitope at which the antibody interacts (through use of centroid estimations) might be obtained by directional labeling of the antibody: If one isotope was located at the 5’ end of the mass-oligonucleotide and another isotope at the 3’ end, it would provide a “pointer” to the epitope is located. The atomic elements used for srIBI can be expanded by synthesizing isotope-derivatized nucleotides with previously reported protocols (*48, 49*). Moreover, a recently developed cesium ion beam has attained currents as high as one pA at 25-nm beam size (*50*), which might enable more precise localizations. Physical expansion of biological samples, mediated by polymer swelling as previously described (*51*), would further increase the resolution of srIBI. Other primary beam sources, such as helium-or neon-based beams, which have beam sizes of 0.5 nm and 1.5 nm (*52*), respectively, have been recently adapted for SIMS (*52*). These improvements may enable 3D imaging of single antibodies, hence opening the door for such opportunities as isotope barcoding to increase the number of simultaneously measured channels. For example, with the seven-detector design presented here, a triple-barcoded labeling would theoretically enable deconvolution of 35 targets.

MIBI and IMC are orthogonal approaches to use of fluorescence for imaging the tissue microenvironment (*13-15*). Currently, these methods are focused on large fields of view and fast acquisition times, due to their focus on large-scale tissue cellular contexts. IMC is based on laser-ablation inductively coupled plasma mass spectrometry with a 1000-nm laser beam. In addition, the cutoff at 75 atomic mass units in the Helios system does not allow imaging of naturally occurring organic isotopes. MIBI has a higher lateral resolution than IMC due to a smaller beam size (260 to 500 nm), but the negative charge of the oxygen duoplasmatron limits detection of intrinsic biomolecules. Thus, current protein-targeted MSI technologies cannot be used for the study of biomolecules or small molecules at the nanoscale. Super-resolution light microscopy methods are limited by the number of fluorophores that can be simultaneously recorded. While recent applications overcome this issue by using cycling of oligonucleotide probes (*2, 3, 5, 8*), these methods suffer from drawbacks such as sample drift between cycles, low axial resolution, photobleaching, and long acquisition times. Current methods such as the tetrapod PSF and DNA-PAINT are promising solutions to some of these limitations (*53, 54*).

Since srIBI acquires data on multiple biomolecules simultaneously, it is an alternative to current super-resolution light microscopy techniques. srIBI samples can be stored for extended periods as the isotopes detected are stable; this is not the case for fluorophores used in light imaging. Furthermore, srIBI enables specific protein staining compatible with direct imaging of exogenously added small molecules. Extension of fluorescent-based methods for specific nucleic acid staining, like CRISPR/Cas-FISH (*55*), Oligopaint (*27, 56*), and ATAC-see (*57*), to srIBI will provide a unique opportunity for an integrative understanding of cellular processes at the nanoscale beyond current means.

Multiplexed imaging of an entire cell, at resolutions afforded by srIBI, introduces new challenges in how to interpret the results. To this end, we have designed and implemented an analytical framework that leverages on established methods to infer biological meaning. For example, hierarchical clustering and t-SNE are able to simplify multiplexed srIBI data across multiple cells. Combined with neighborhood interaction maps, we are able to reveal interactions within the subcellular domain not obvious to the naked eye. Future improvements should include better methods to extract data meaningfully. The ability to learn and infer information about other unlabeled but related biomolecules within the cell will provide a way to describe an entire cell with a limited number of parameters (*58, 59*). Further integrative analysis, particularly incorporation of biophysical and genomic information, will also allow more accurate modeling of nuclear structures (*8, 60, 61*). Here, we applied srIBI to study the localization of the cancer drug cisplatin. Cisplatin accumulation was observed in SC35-positive nuclear speckles, which are sites of mRNA processing and splicing. We were able to distinguish between cisplatin at active regions of transcription (H3K27Ac) and nuclear speckles (SC35), placing the drug functionally at sites of RNA splicing in addition to general sites of active transcription. Indeed, cisplatin has been shown to reduce pre-mRNA splicing in a dose-dependent manner which might be additive to its well-known role inducing adducts, single strand gaps post repair and other lesions in DNA (*39, 62*). These results suggest potential vulnerabilities of mRNA processing that can be targeted for combinatorial therapies to increase the effectiveness of cisplatin.

SrIBI is an imaging method that can be applied in single-cell studies of the diverse molecular interactions in subcellular microenvironments. Combination with other unbiased or population-based methods, such as small molecule screens and multiplexed proteomics, promises to reveal new understanding into metabolic pathways or mechanisms of drug resistance. This has the potential to drive new therapeutic discoveries and inform decision-making processes of small molecules.

## Acknowledgments

We thank Maurice Lee and Drs. Michael Angelo and Maria Angulo-Ibanez for critical discussions and reading the manuscript. We thank Angelica Trejo and Gina Jager for technical contributions. We are grateful to the Protein and Nucleic Acid Facility of Stanford University for production of oligonucleotides and technical support;

## Author contributions

X.R.-C. designed and performed most of the experiments, analyzed and interpreted data and wrote the manuscript. S.J. designed and performed experiments, designed the iterative srIBI data analysis, analyzed and interpreted data and wrote the manuscript. Y.B. performed experiments. G.B. and S.B. designed the nuclear neighborhood analysis, analyzed and interpreted data. A.F.C. and G.H. assisted in experimental design. B.Z analyzed data. C.-M.K.H. performed experiments and assisted in optimizing protocols used in MoC-Ab staining. C.H. assisted in data acquisition and experimental design. S.-Y.C. designed and performed experiments, analyzed and interpreted data and wrote the manuscript. F.A.B. assisted in optimizing protocols used in MoC-Ab staining. G.P.N. was responsible for funding acquisition, assisted in experimental design, interpreted data and wrote the manuscript.;

## Data and materials availability

All data is available in the main text or the supplementary materials. Data generated and analyzed during the current study are available from the corresponding author on reasonable request.

## Supplementary Materials

## Materials and Methods

### Cells

TYK-nu cells were grown in Eagle’s minimal essential medium (American Tissue Culture Collection). HeLa cells were grown in DMEM (Gibco, Invitrogen). Media were supplemented with 10% heat-inactivated fetal bovine serum, 100 U/mL penicillin (Gibco, Invitrogen), and 100 mg/mL streptomycin (Gibco, Invitrogen). Cells were cultured in a humidified cell incubator at 37 °C with 5% CO2 conditions and split with TrypLE Express (Gibco, Invitrogen) every 2-3 days.

### MoC-Ab preparation

Oligonucleotides (Table S1) were synthesized at the Stanford Protein and Nucleic Acid Facility with internal isotope-derivatized nucleotides, a fluorophore at the 3’ position, and a maleimide cycloadduct at the 5’ position. The maleimide was deprotected by a retro Diels-Alder reaction. The lyophilized oligonucleotide was suspended in 1 mL anhydrous toluene (MTX07327, Millipore) for 4 hours at 90 °C, washed four times with anhydrous ethanol, and solubilized in buffer C (2 mM Tris, 150 mM NaCl, 1 mM EDTA, pH 7.2). The oligonucleotide concentration was determined using a Nanodrop spectrophotometer. Aliquots of 8.5 nmol were prepared, lyophilized overnight, and stored in a desiccator at −20 °C. Antibodies (Table S2) in carrier-free PBS were conjugated to the deprotected oligonucleotides. Briefly, 50 µg of antibody was loaded into a 50-KDa 0.5-mL centrifugal filter column with 400 µL PBS and reduced with 400 µL reduction buffer (PBS with 2.5 mM TCEP and 2.5 mM EDTA) for 30 minutes at room temperature. Antibodies were then washed with 400 µL of C buffer into a 50-KDa 0.5-mL centrifugal filter column and conjugated to 8.5 nmol of oligonucleotide in 400 µL conjugation buffer (buffer C with 0.5 M NaCl) for 2 hours at room temperature. Antibodies were washed five times with 400 µL of high-salt PBS (PBS with 1 M NaCl), diluted into storage buffer (Candor PBS Antibody Stabilization Solution with 0.5 M NaCl and 5 mM EDTA) and stored at 4 °C. Each MoC-Ab was titrated by immunofluorescence using HeLa cells, as exemplified in Fig. S3, and the staining pattern was compared to the staining pattern of the unconjugated antibody.

### Lanthanide-conjugated and biotin-conjugated antibodies

Antibodies (Table S2) in carrier-free PBS were conjugated to metal-chelated polymers (MaxPAR Antibody Conjugation Kit, Fluidigm) or Sulfo-NHS-SS-Biotin (A39258, ThermoFisher) according to the manufacturer’s protocol. Antibodies were diluted to 0.2 mg/mL in Candor PBS Antibody Stabilization Solution and stored at 4 °C.

### Evaluation of cisplatin accumulation in cells by mass cytometry

TYK-nu cells were cultured in 0.5, 5, or 50 µM of cisplatin (P4394, Sigma-Aldrich) for 24 hours, washed, and treated with 1 µM Rh-intercalator (201103B, Fluidigm) for 15 minutes to discriminate dead from live cells. Cells were analyzed in a CyTOF2 instrument (Fluidigm) as previously described (63, *64*).

### Intracellular staining

Compositions of buffers used for staining are given in Table S3. Cells were fixed and permeabilized during 30 minutes at 4 °C in Fixation/Permeabilization buffer. Cells were then gently washed three times with Wash Buffer and blocked in Block Buffer 1 during 30 minutes at room temperature, washed three times with Wash Buffer, blocked in Block Buffer 2 during 30 minutes at room temperature, and washed three times with Wash Buffer. The cells were stained with a mixture of MoC-Abs in Reaction Buffer for 3 hours at room temperature. Following staining with MoC-Abs, the Reaction Buffer was removed by gently touching the sample with a precision wipe (05511, Kimtech Science). Cells were then washed twice in Wash Buffer and incubated with Reaction Buffer containing FluoroNanogold Streptavidin (1:40 titer, 7016, Nanoprobes) for 30 minutes at room temperature. Cells were washed twice with Wash Buffer before preparing them for confocal microscopy or srIBI analysis. Cell fixation in PBS with 1.6% paraformaldehyde for 10 minutes at room temperature, permeabilization in methanol (10 minutes at -20 °C) or with Triton X-100 (0.5% in PBS, 10 minutes at room temperature) and staining produced comparable results for most of the antibodies tested.

### Confocal microscopy

For confocal microscopy analysis, cells were grown in 12-mm diameter glass coverslips (72226-01, Electron Microscopy Sciences) until 80-90% confluent and stained as described. After staining, cells were washed three times with PBS, rinsed once with water, and mounted using Vectashield with DAPI (H-1200, Vector Laboratories). Anti-mouse-Alexa488 (1:2000 titer, 4408S, Molecular Probes) or anti-mouse-Alexa647 (1:2000 titer, 4410S, Molecular Probes) were the secondary antibodies used to detect unconjugated antibodies. Cells were analyzed in a LSM 880 (Zeiss) with a 63x oil-immersion objective (Zeiss Plan-Apochromat 63x/1.4 Oil).

### Super-resolution Ion Beam Imaging

For srIBI analysis, cells were grown on silicon wafers (7 nm * 7 nm or 18 nm * 18 mm, Silicon Valley Microelectronics). Wafers were rinsed twice in methanol, air-dried with compressed air, washed with ethanol for 10 minutes, and rinsed three times with sterile PBS in a cell culture hood prior to cell seeding. When cells reached 80-90% confluence, they were stained as described above. After staining, cells were washed twice in PBS, fixed for 5 minutes in Post-Fixation buffer, and rinsed five times with water. Cells were dehydrated using a graded ethanol series, air dried in a desiccator chamber, and stored at room temperature in a vacuum desiccator until analysis. srIBI images were acquired with the NanoSIMS 50L mass spectrometer (Cameca) at Stanford University using the CAMECA Microbeam Cesium Source. Before each experiment, tuning of the primary optics, secondary optics, and mass spectrometer was performed. The detectors were tuned by identifying each isotope of interest in an air-dried drop placed on a silicon wafer. The ion detectors were set as follows: detector 1, ^12^C; detector 2, ^19^F; detector 3, ^31^P; detector 4, ^81^Br; detector 5, ^127^I; detector 6, ^194^Pt; and detector 7, ^197^Au. Secondary electrons were detected in parallel with ion information. The secondary ion peaks and E0S were tuned before the acquisition of each new cell to correct for magnetic drift. All images were collected with 1-ms dwell time per pixel. The raster size, number of pixels, repeat scans over the same area, and total scan time for each image are listed in Table S4. For each srIBI image, individual planes described in Table S4 were summed and gaussian blurred with a sigma of one using FIJI.

### Nuclear neighborhood analysis

A mask for each nucleus was created using the phosphorus channel, and 20000 3D voxels, each of dimension (x, y, z) = (6, 6, 20) pixels, were extracted from each of two cells by random sampling from the middle 40 Z-planes (for a total of 40000 voxels). Each channel was z-normalized across all pixels contained within the sampled image regions. Unsupervised hierarchical clustering was performed using the markers SC35, H3K9me3, phosphorus, nucleolin, and H3K27Ac to identify 10 clusters across the 20000 extracted voxels. The average expression of each marker from both cells was then plotted in a heatmap form. Each voxel was colored by its cluster, and replotted on a X-Y plane to recreate the cell. The count distribution of cisplatin within regions assigned to each cluster was computed for each cell.

### Iterative srIBI data analysis

The low pixel signal intensity values of iterative srIBI due to the much lower current during data acquisition than in conventional srIBI is a barrier for robust data analysis. We adapted a filtering strategy described in Keren et al. to filter noise from signal using a k nearest neighbor (KNN) approach (*14*). The assumption for this approach is that in sparse data, the signal density offers more information than signal intensity. Thus, regions with dense signals are more likely to contain true signals, compared with regions with sparse signals. We first summed the image data (1024 x 1024 pixel) across 20 Z-planes, and then measured the average distance between positive pixels for the nearest 25 neighbors within each channel. We next applied a cutoff to distinguish between signal (regions of high signal density) and noise (regions with sparse signal density), to create a mask for each channel. The masks were applied to each individual Z-plane to create a denoised image for subsequent analysis. The 100 pixels around each border were discarded to avoid potential edge effects. A 3D sliding window of dimension (x, y, z) = (10, 10, 5) pixels and step-size of (x, y, z) = (5, 5, 3) was applied to the image to calculate the mean of each channel within the voxel. We randomly sampled 20000 voxels across each image (from 5 different cells) to obtain a final 100000 voxels. Counts were log2 transformed after adding a “0.0001” value to avoid zeros. Unsupervised hierarchical clustering was performed on all voxels using the markers SC35, H3K9me3, phosphorus, nucleolin, H3K27Ac, and cisplatin. A t-Distributed Stochastic Neighbor Embedding (t-SNE) using a Barnes-Hut implementation was performed, and each identified cluster was differentially colored. The means of each cluster was also plotted and subsequently manually merged by similarity into 11 final clusters. The t-SNE plot was also colored by the cell of origin of each voxel to ensure minimal batch effects.

### Nuclear neighborhood interaction frequency

To identify and test for the significance of how frequently clusters interacted (as defined by the number of pixels separating voxels from each other), we implemented a permutation test. First, the Euclidian distance in 3D space between voxels was calculated (Equation 1).

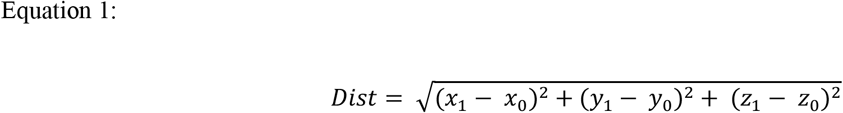

Next, we defined interacting voxels as those less than or equal to 5 pixels away from each other (between the center of each voxel). After 1000 permutations, where the cluster annotation was shuffled, and the p-value calculated on either tail (Equation 2).

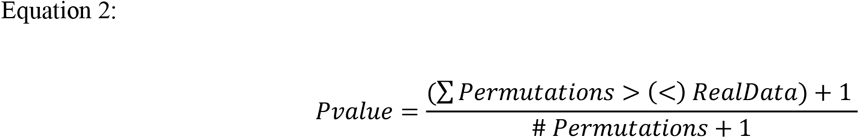

Essentially, 1 plus the number of permutations with distances smaller than (or greater than) the real data was divided by 1 plus the total number of permutations performed. Finally, the fold enrichment of interactions for real data over the mean interactions from the permutation tests was calculated (Equation 3).

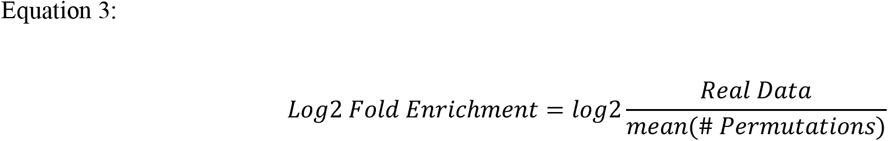

**Figure S1.**
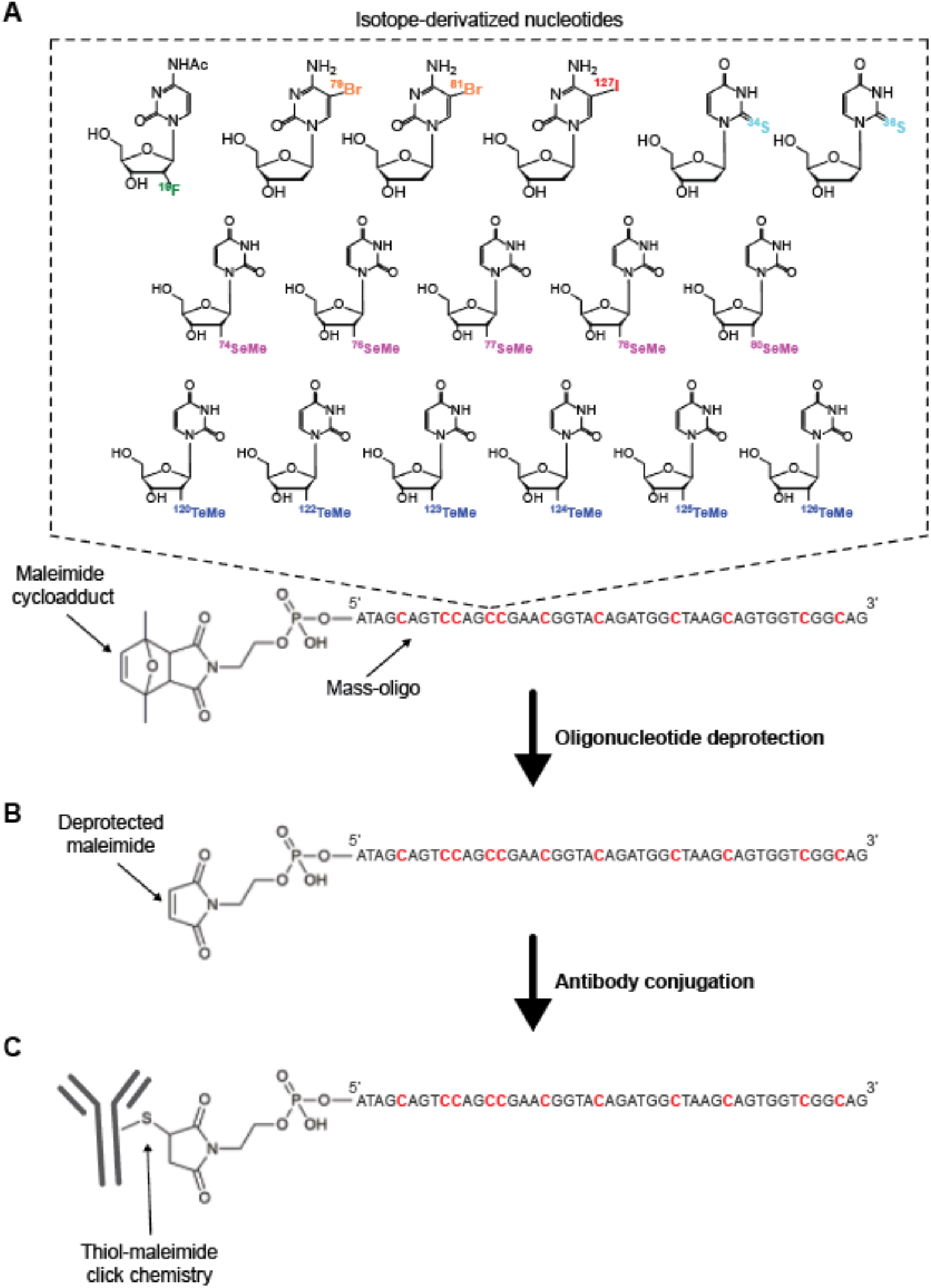
Strategy for the synthesis and conjugation of mass-oligonucleotides to antibodies. (**A**) Mass-oligonucleotides were synthesized with internal isotope-derivatized nucleotides and a maleimide cycloadduct at the 5’ position. Dashed box shows chemical structures of isotope-derivatized cytidines and thymidines that are commercially available or that can be synthesized with previously reported protocols (*48, 49*). In this study, we incorporated 11 fluorine, bromine, or iodine-derivatized cytidines (shown in red) into a 47-nucleotide long oligonucleotide. The DNA sequence was selected to minimize potential for binding to the human genome. (**B**) Oligonucleotides were deprotected in toluene for 4 hours at 90 °C, washed in ethanol, lyophilized, and stored at -20 °C. (**C**) The deprotected oligonucleotides were conjugated to partially reduced antibodies through thiol-maleimide click chemistry. Free oligonucleotides were removed via size-exclusion by centrifugation using a 50-KDa MWCO filter.

**Figure S2.**
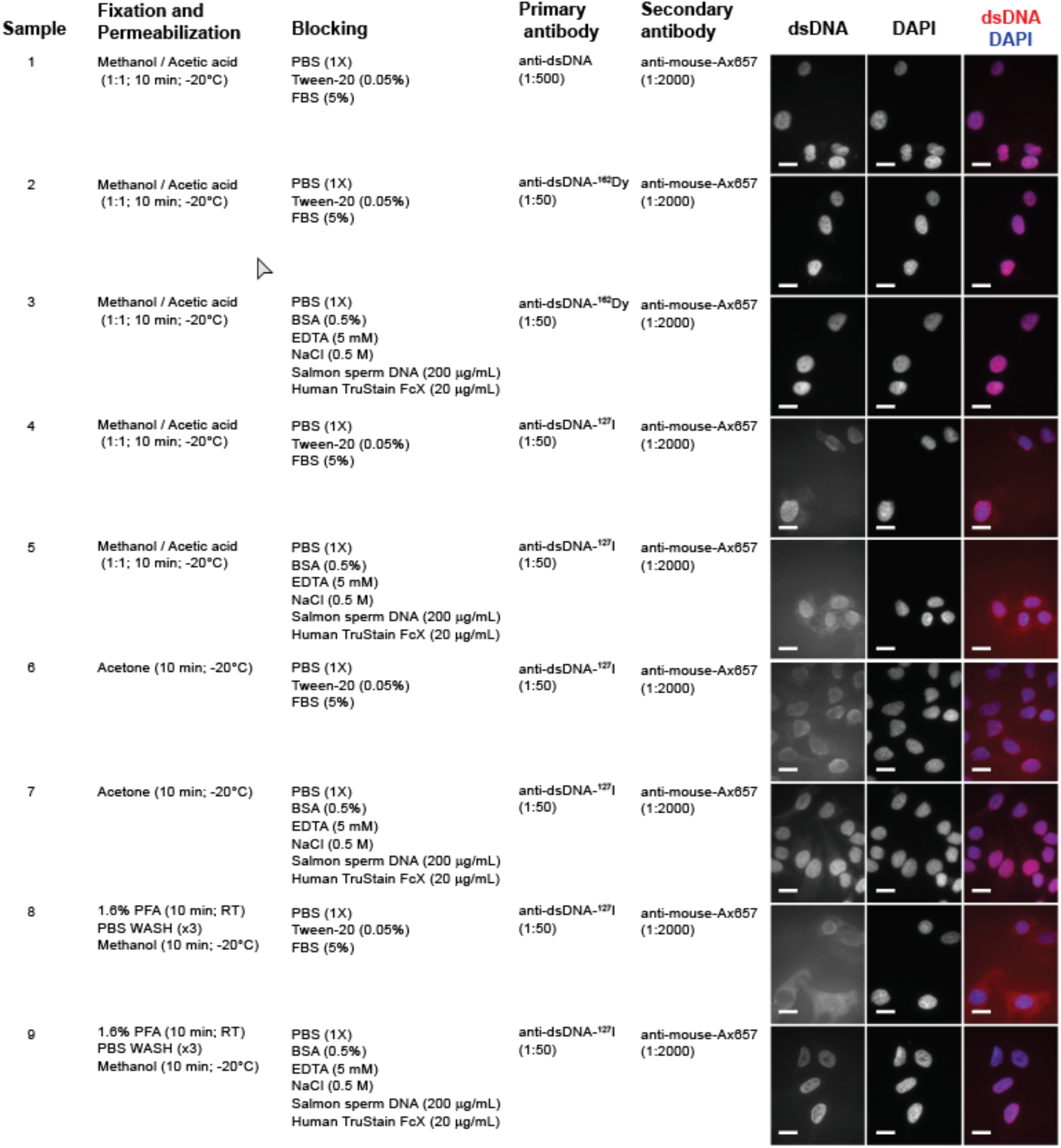
Protocol optimization for intracellular staining with MoC-Abs. HeLa cells were fixed and permeabilized as indicated in the second column. Cells were then blocked for 30 minutes at room temperature as indicated in the third column, stained for 1 hour at room temperature with unconjugated, lanthanide-polymer conjugated (^162^Dy) or mass-oligonucleotide-conjugated (^127^I) primary anti-double-stranded DNA (anti-dsDNA), washed three times in PBS, and stained for 1 hour at room temperature with secondary anti-mouse-Alexa 647. Cells were then washed and mounted with fluoromount containing DAPI for confocal microscopy analysis. In HeLa cells stained with anti-dsDNA (sample 1), specific nuclear staining co-localizing with DAPI was observed as expected. HeLa cells stained with anti-dsDNA-^162^Dy (sample 2) showed the same nuclear staining indicating that that the conjugation process did not alter antibody specificity. The presence of high salt concentration and salmon sperm DNA in cells stained with anti-dsDNA-^162^Dy (sample 3) did not interfere with antibody staining. HeLa cells fixed and permeabilized with methanol (sample 4), acetone (sample 6), or fixed with 1.6% paraformaldehyde and permeabilized with methanol (sample 8) and stained with anti-dsDNA-^127^I showed nonspecific cytoplasmic staining. The addition of high salt concentration and salmon sperm DNA in the blocking buffer of HeLa cells fixed and permeabilized with methanol (sample 5) or acetone (sample 7) and stained with anti-dsDNA-^127^I were stained nonspecifically, although the nonspecific cytoplasmic staining was partially reduced in acetone-treated cells. Fixation with 1.6% paraformaldehyde, permeabilization with methanol, and blocking with a high salt concentration and salmon sperm DNA (sample 9) resulted in a highly specific and background-free visualization of dsDNA in the nucleus of HeLa cells stained with anti-dsDNA-^127^I. Permeabilization with other reagents such as Triton X-100 or saponin was also compatible with specific MoC-Ab staining (data not shown). Scale bars, 20 µm.

**Figure S3.**
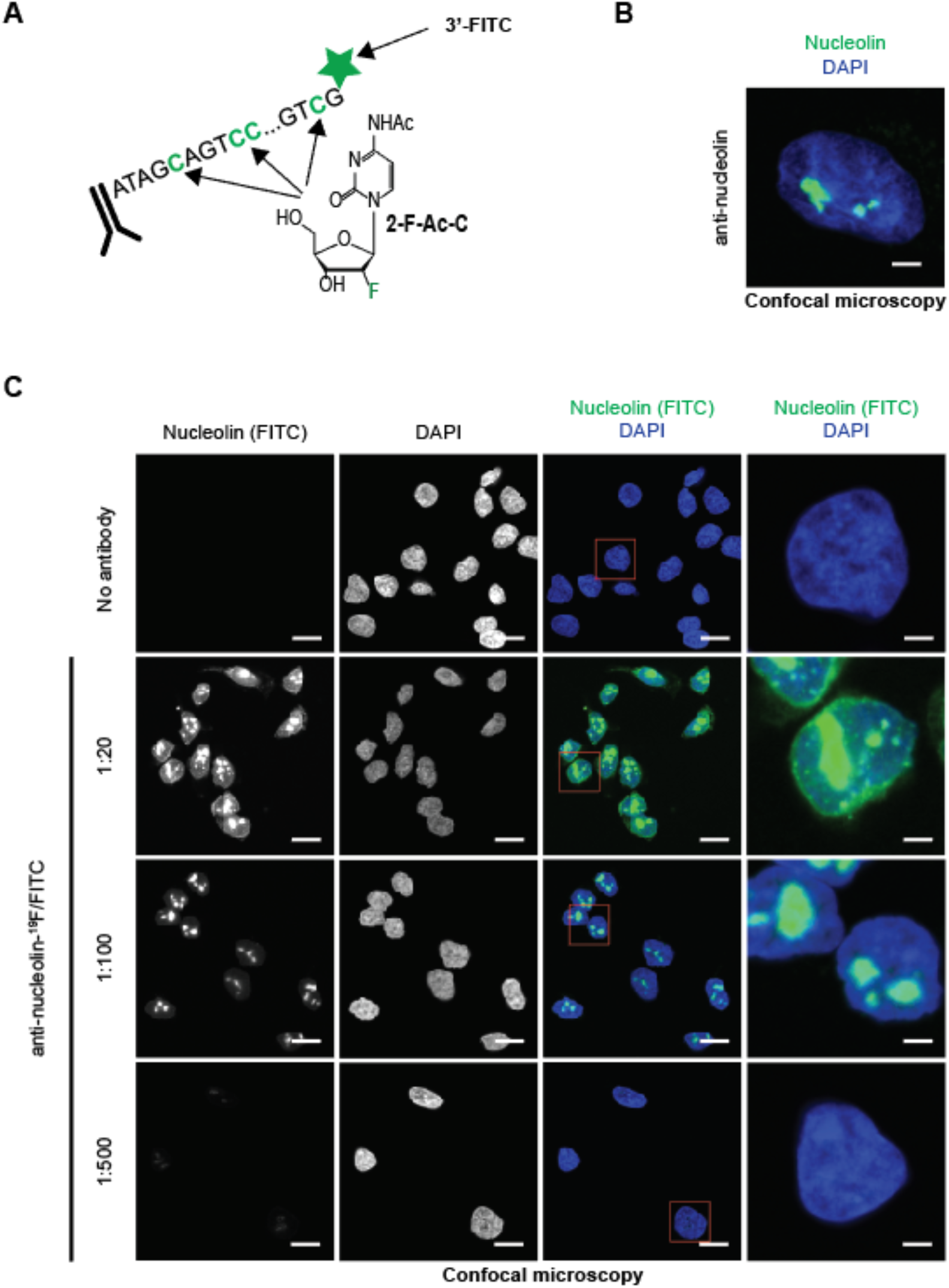
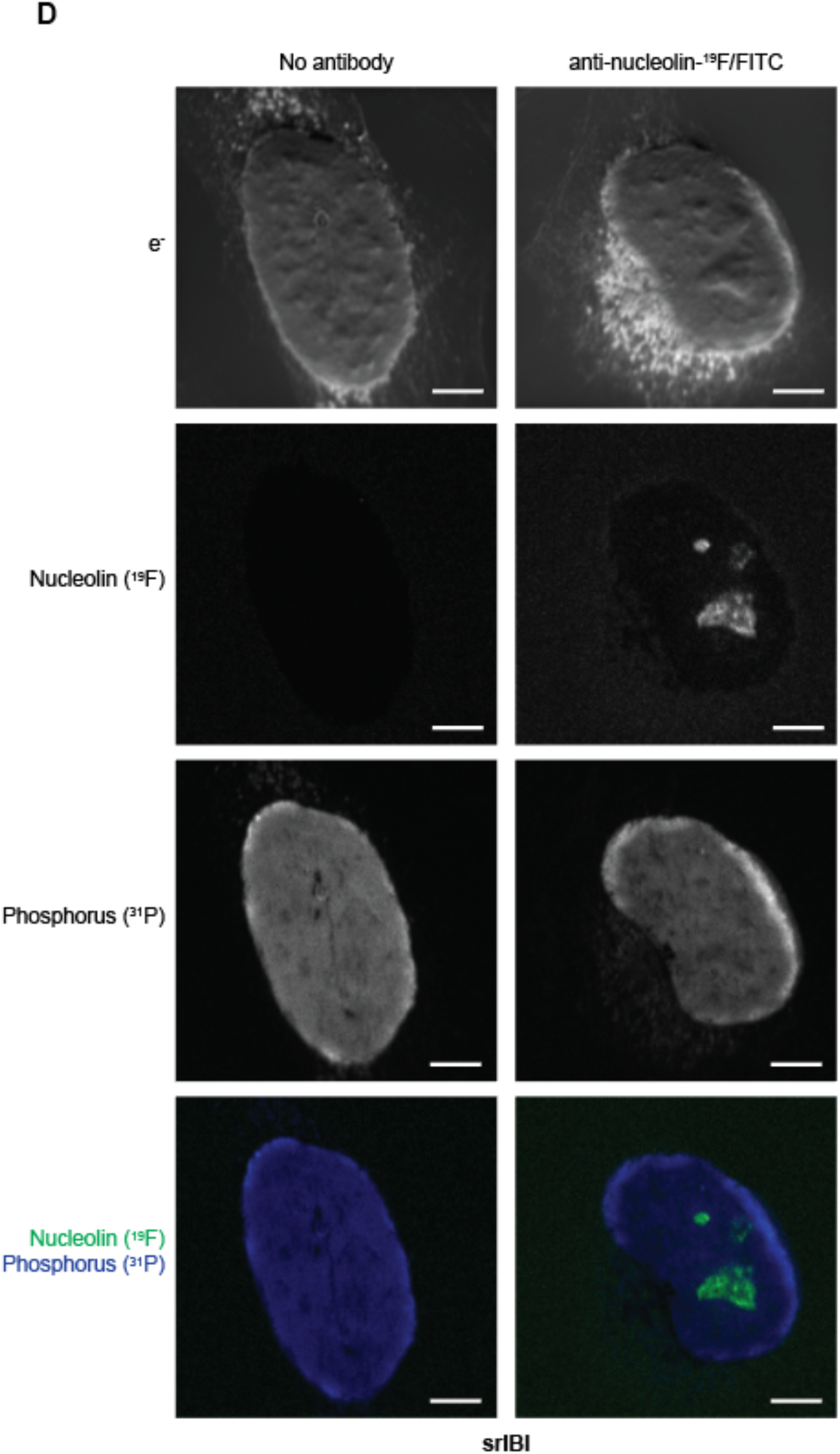
Validation of ^19^F/FITC MoC-Abs for srIBI. (**A**) A schematic of the ^19^F/FITC MoC-Ab. The antibody was conjugated to the 5’ end of an oligonucleotide in which all cytidines were replaced with 2-F-Ac-C. The oligonucleotide also had a 3’ FITC. (**B**) Representative confocal microscopy image of a HeLa cell stained with unconjugated anti-nucleolin-^19^F/FITC and a secondary anti-mouse-Alexa488 (green) and DAPI (blue). Scale bar, 4 µm. (**C**) Representative confocal microscopy images of control HeLa cells (no antibody) or HeLa cells stained with anti-nucleolin-^19^F/FITC (green) at different concentrations. ^31^P is shown in blue. Cells in red boxes in the composite image are shown enlarged in the right column. Scale bars, 20 µm in standard images and 4 µm in enlarged images. A 1:100 dilution of this antibody was used in subsequent experiments. (**D**) Representative srIBI images of a control HeLa cell (no antibody) and HeLa cells stained with anti-nucleolin-^19^F/FITC. Scale bar, 4 µm.

**Figure S4.**
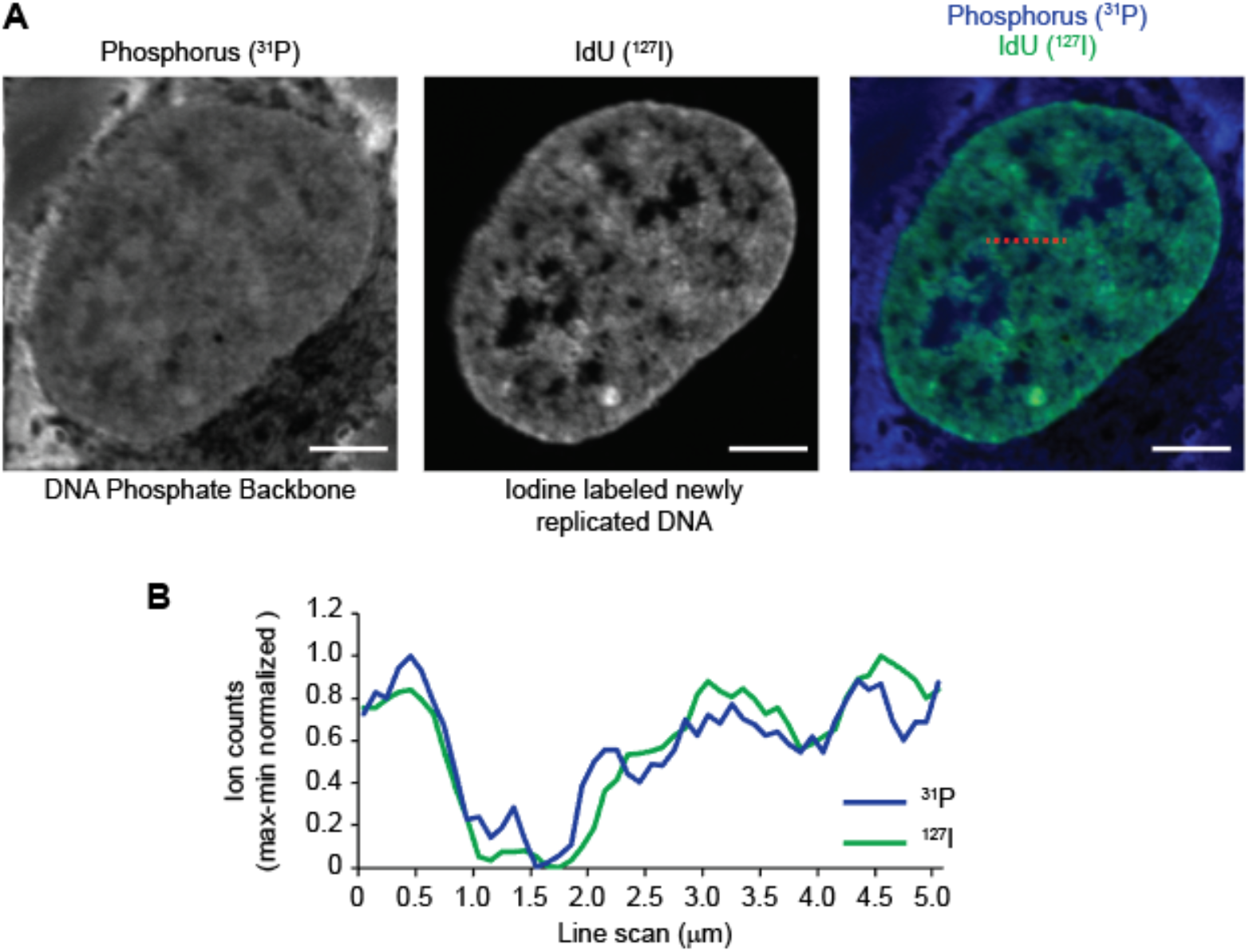
High-phosphorus regions within the nucleus mainly originate from the DNA backbone. (**A**) Representative srIBI image of a HeLa cell treated with 5-iodo-2’-deoxyuridine (IdU) for 24 hours to label DNA. Ion images for phosphorus (^31^P; left panel) and newly synthesized DNA (^127^I; middle panel) are overlaid in the right panel. Scale bars, 5 7m. (**B**) Line scan along the dashed red line in the right image in panel A. Raw ion counts were scaled by max-min normalization. Newly replicated DNA (^127^I; green line) and phosphorus (^31^P; blue line) in the nucleus show a similar pattern, confirming that high-phosphorus regions mainly originate from the DNA backbone rather than from RNA or phosphorylated proteins.

**Figure S5.**
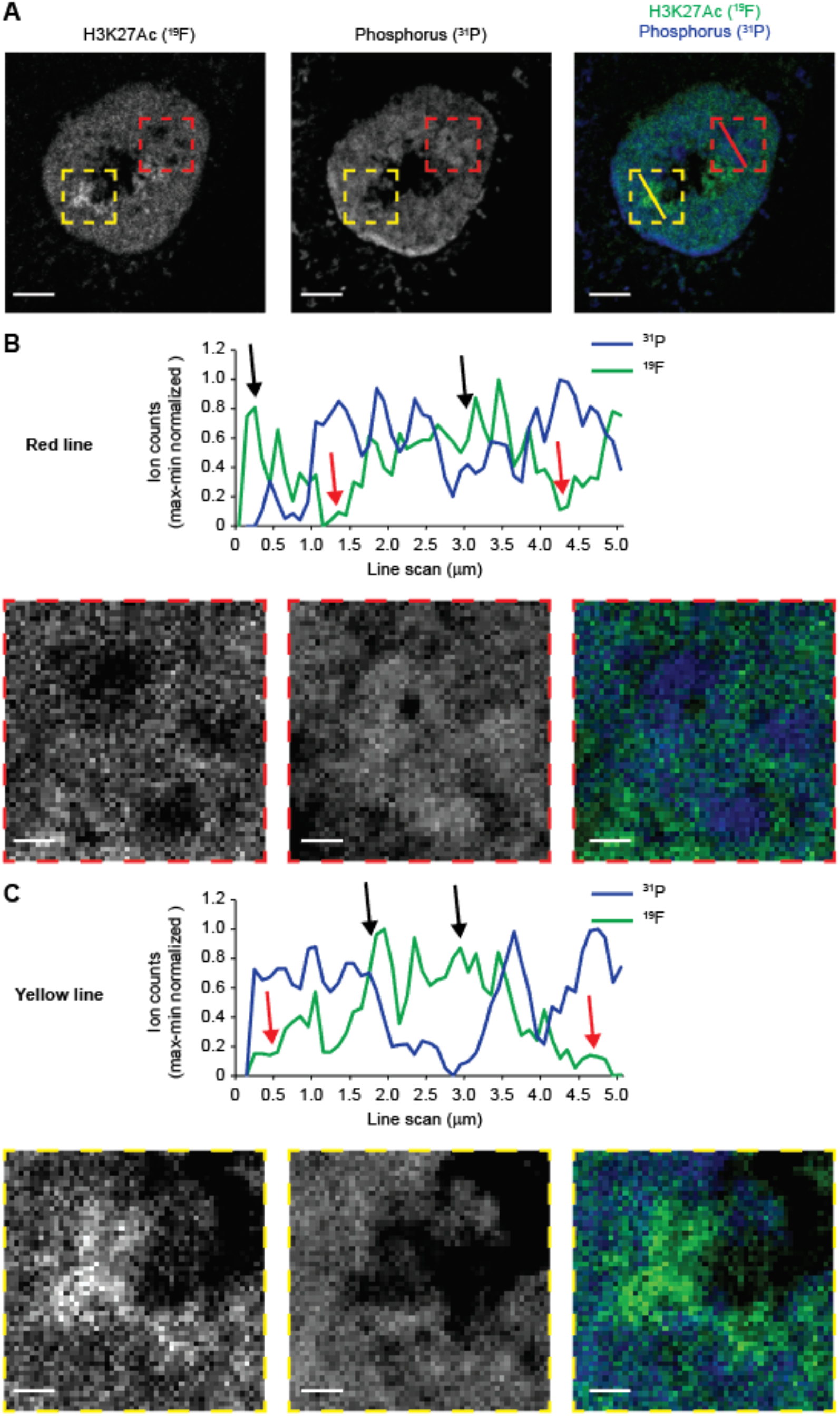
Regions of active transcription within the nucleus mainly originate from low-phosphorus regions. (**A**) Representative srIBI image of a HeLa cell stained with anti-H3K27Ac-^19^F/FITC. Ion images for H3K27Ac (^19^F; left panel) and phosphorus (^31^P; middle panel) are overlaid in the right panel. Scale bars, 4 µm. Cell in the composite image is also shown in Fig. 1D. (**B**-**C**) Line scan along the lines in the boxes outlined in **B**) red and **C**) yellow dashed lines in the image to the right in panel A. Raw ion counts were scaled by mix-man normalization. Black arrows point to regions of high H3K27Ac counts (green line) that are anti-correlated with phosphorus counts (blue line). Red arrows point to low H3K27Ac counts (green line) that are anti-correlated with phosphorus counts (blue line). Images below the line scans are magnified from regions boxed in panel A. Scale bars, 800 nm.

**Figure S6.**
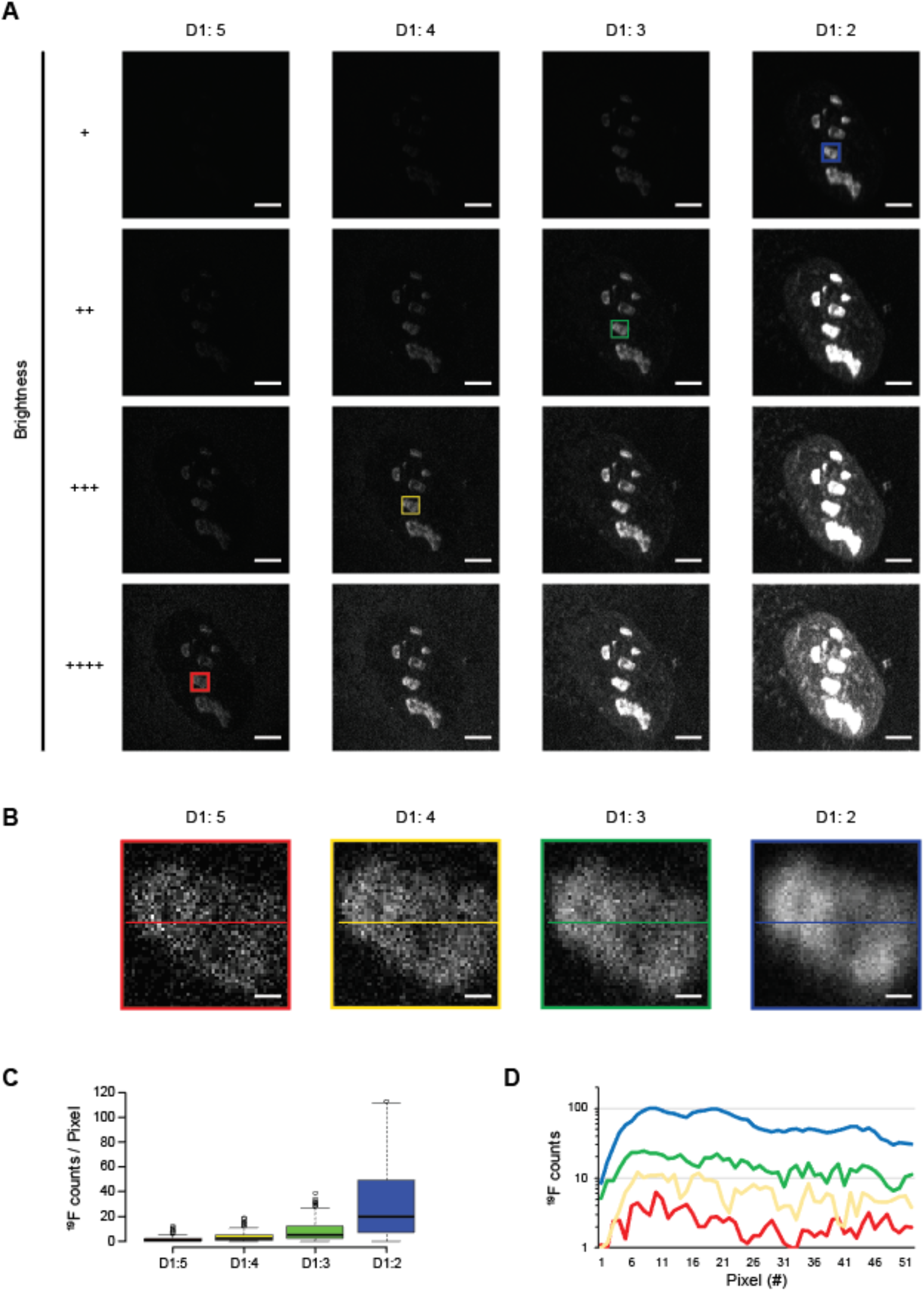
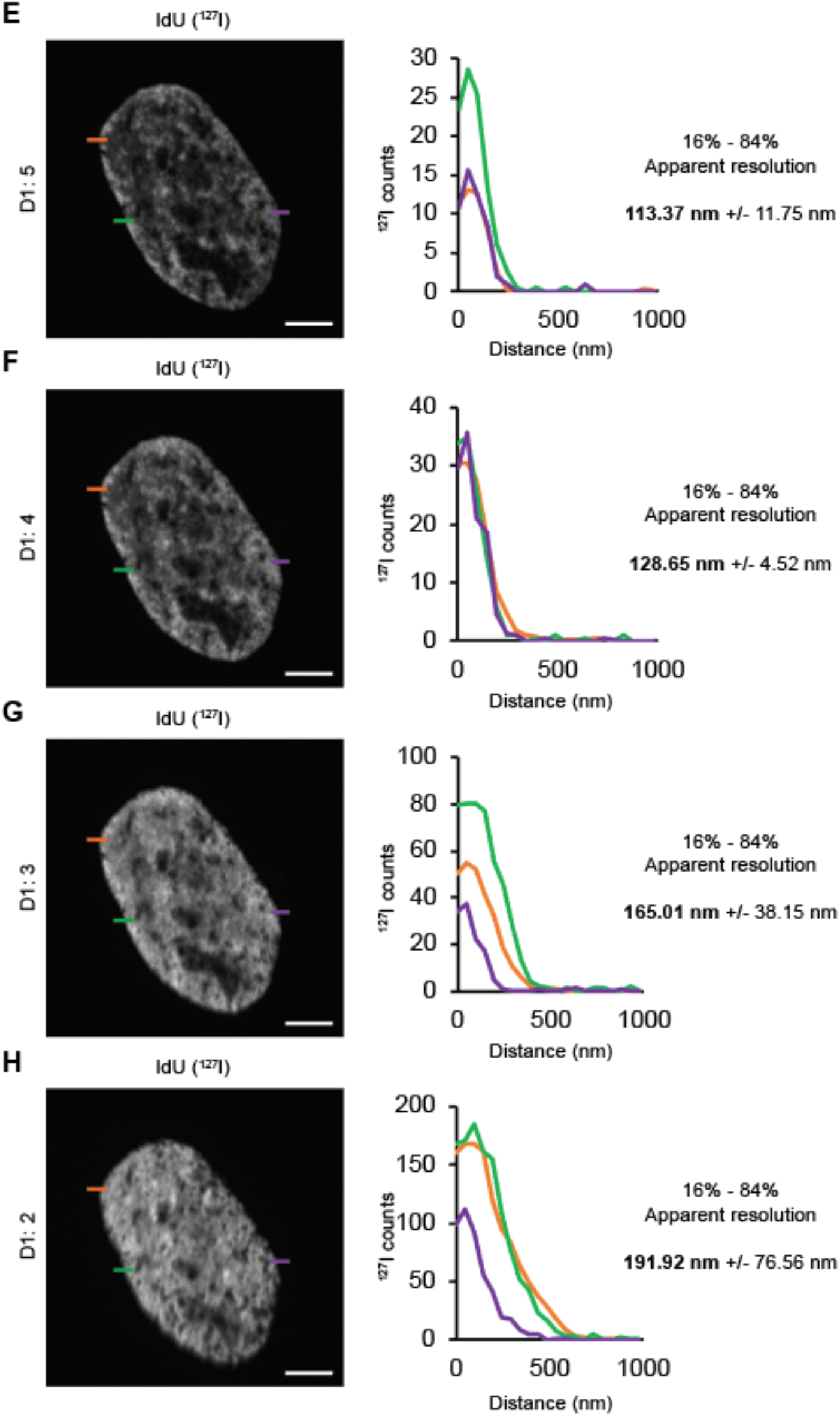
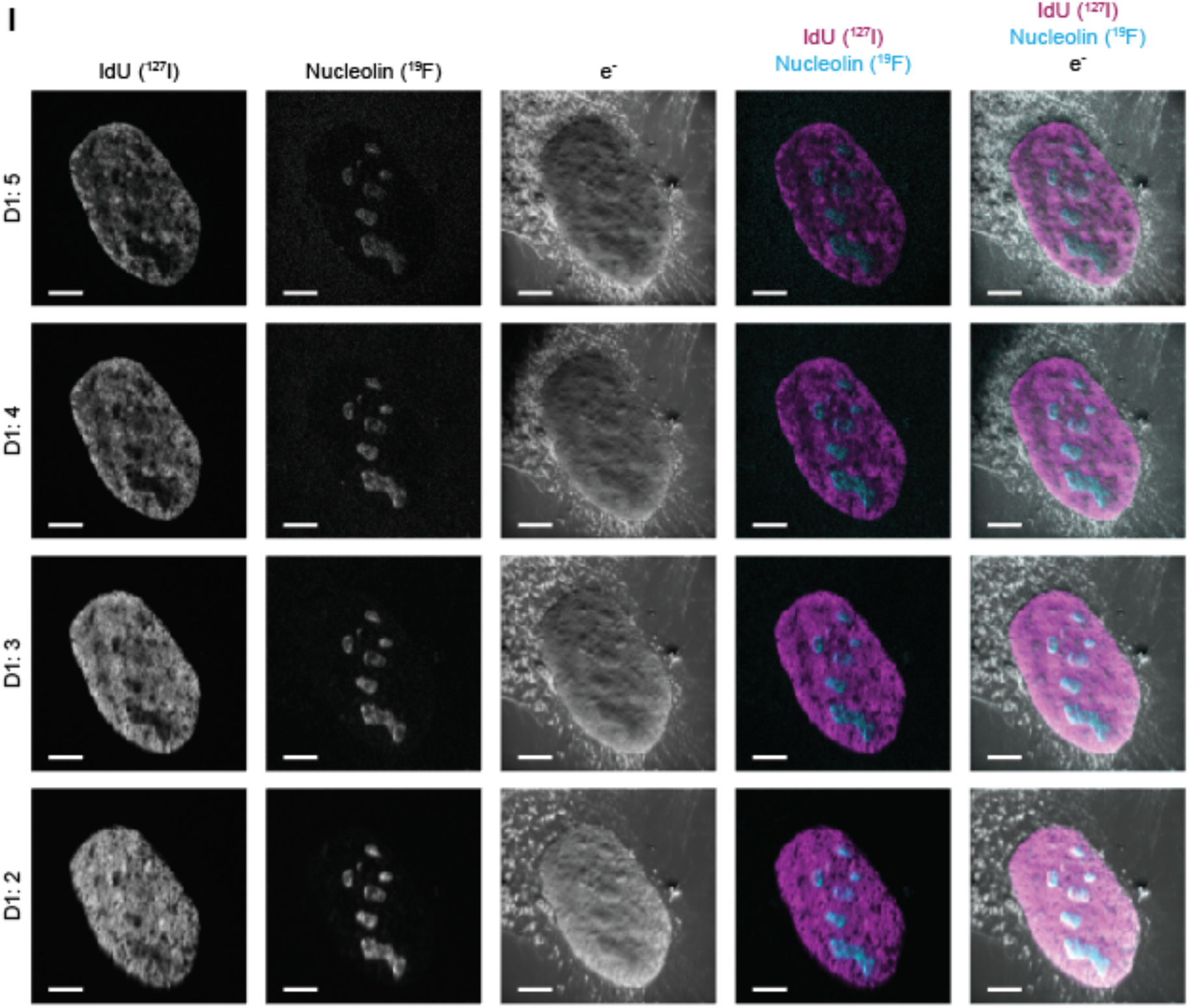
Beam current scales with ion counts per pixel at the expense of resolution. HeLa cells were labeled with IdU for 24 hours and stained with anti-nucleolin^-19^F/FITC. The images were acquired with a constant 25 * 25 µm field of view, 1 ms dwell time per pixel, and 512 * 512 pixels (xy). The beam current was increased every 10 planes by changing the D1 aperture width to obtain a beam diameter of ∼50 nm (D1: 5), ∼75 nm (D1: 4), ∼100 nm (D1: 3) and ∼150 nm (D1: 2). (**A**) Representative srIBI images of nucleolin staining in the same HeLa cell using different aperture widths. Different brightness intensities to normalize for variation in ion counts are shown for clarity. Scale bars, 4 µm. (**B**) Enlarged images of colored boxes within images in panel A show an individual nucleolus. Details are increasingly lost as the beam current is elevated. Scale bars, 400 nm. (**C**) Ion count per pixel of all pixels in images shown in panel B. Each colored boxplot represents color-coded images in panel B. Boxplots at high current (D1: 2 and 3) show higher ion counts per pixel than boxplots at low current (D1: 4 and 5), highlighting loss of ion counts per pixel at lower current. (**D**) Line scans from lines within images shown in panel B. Line scans at high current (D1: 2 and 3) are smoother than line scans at low current (D1: 4 and 5), showing loss of resolution at higher current. (**E-H**) Quantification of lateral resolution using a beam diameter of **E**) ∼50 nm (D1: 5), **F**) ∼75 nm (D1: 4), **G**) ∼100 nm (D1: 3), and **H**) ∼150 nm (D1: 2). (Left) Representative srIBI images of newly replicated DNA (IdU) staining in the same HeLa cell. Scale bars, 4 µm. (Right) Line scan for each colored bar in the images shown on the left. The lateral resolution calculated using the 16%-84% criterion is shown as an average of each analyzed line ± s.d. (**I**) Single channel and composite images of images shown in Fig. 1E of ^-^e, nucleolin, and IdU.

**Figure S7.**
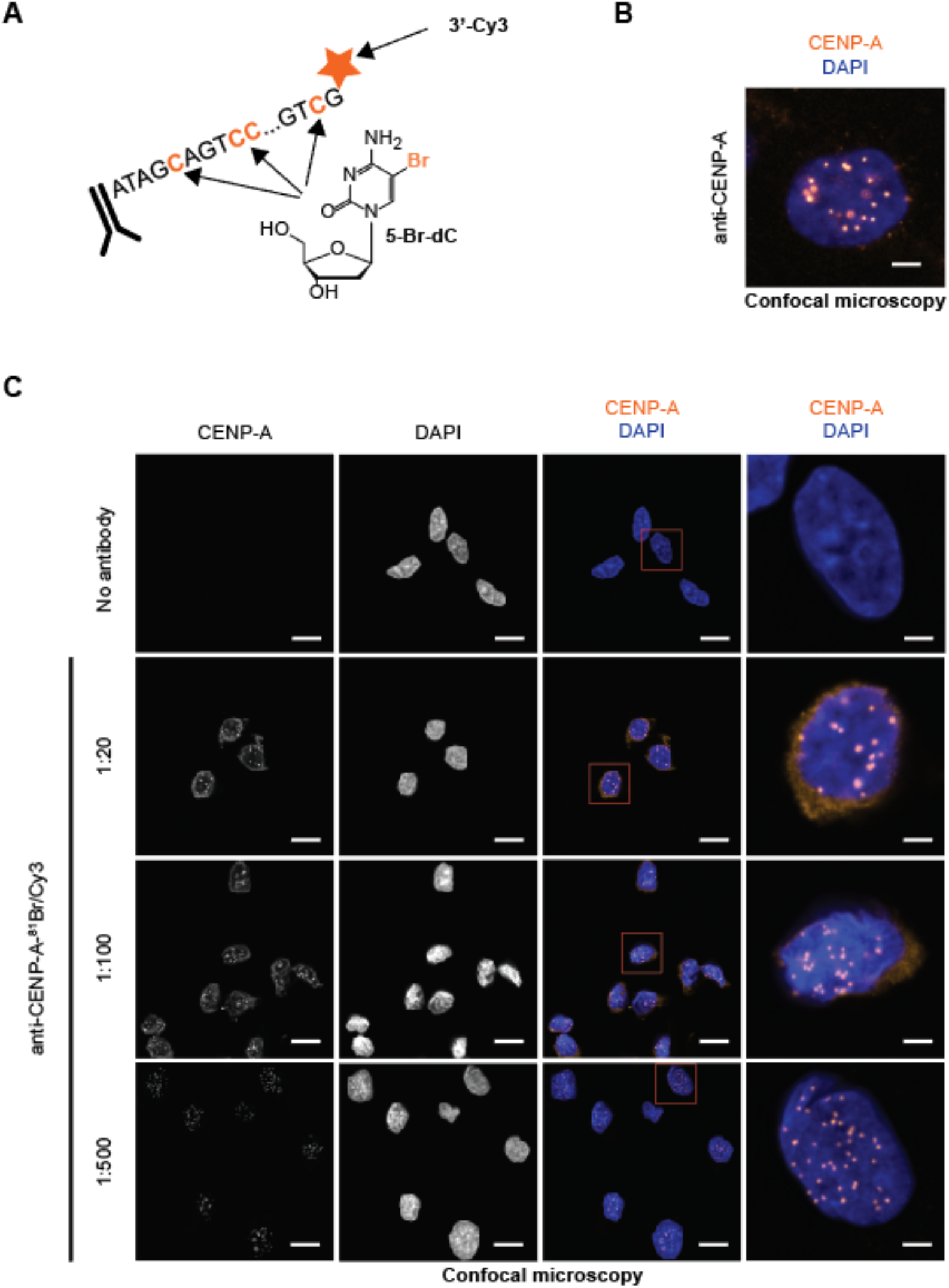
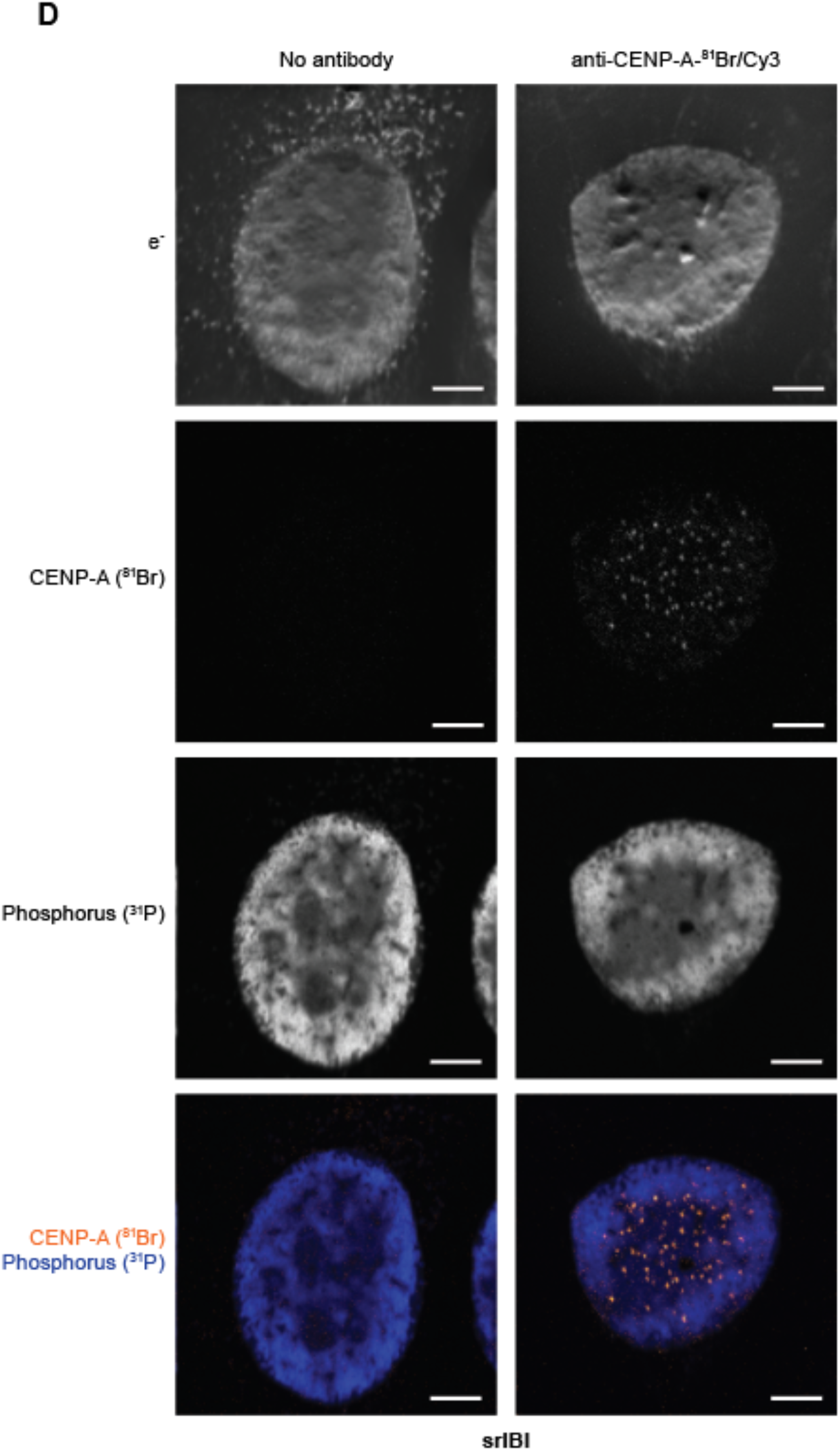
Validation of ^81^Br/Cy3 MoC-Abs for srIBI. (**A**) A schematic of the ^81^Br/Cy3 MoC-Ab. The antibody was conjugated to the 5’ end of an oligonucleotide in which all cytidines were replaced by 5-Br-dC. The oligonucleotide also had a 3’ Cy3. (**B**) Representative confocal microscopy image of a HeLa cell stained with unconjugated anti-CENP-A-^81^Br/Cy3 and a secondary anti-mouse-Alexa488 (green) and DAPI (blue). Scale bar, 4 µm. (**C**) Representative confocal microscopy images of control HeLa cells (no antibody) and HeLa cells stained with anti-CENP-A-^81^Br/Cy3 (orange) at three different concentrations. ^31^P is shown in blue. Cells in red boxes in the composite image are enlarged in the right column. Scale bars, 20 µm in standard images and 4 µm in enlarged images. A 1:500 dilution of this antibody was used in subsequent experiments. (**D**) Representative srIBI images of control HeLa cell (no antibody) and HeLa cell stained withanti-CENP-A-^81^Br/Cy3.Scale bar, 4 µm.

**Figure S8.**
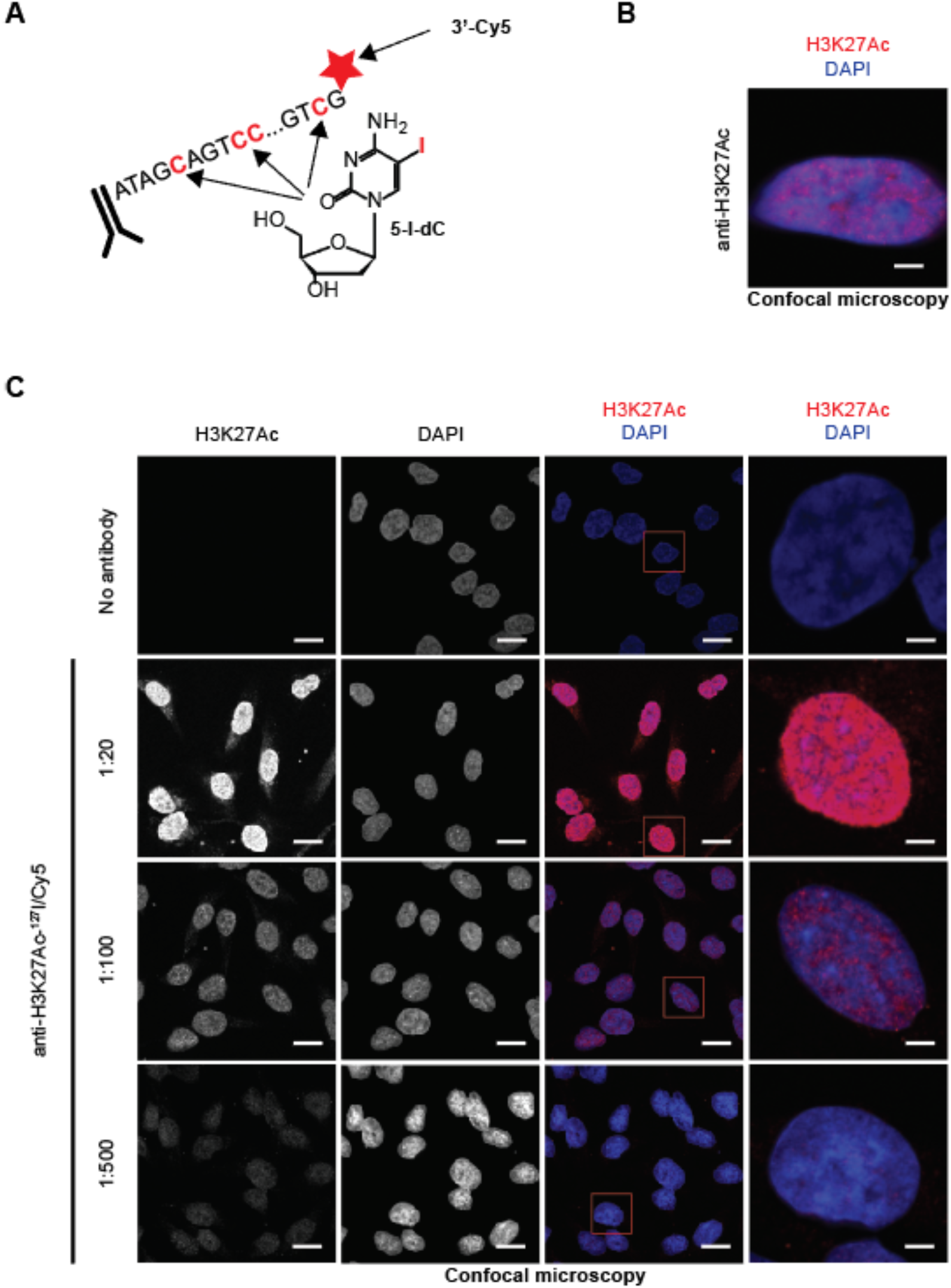
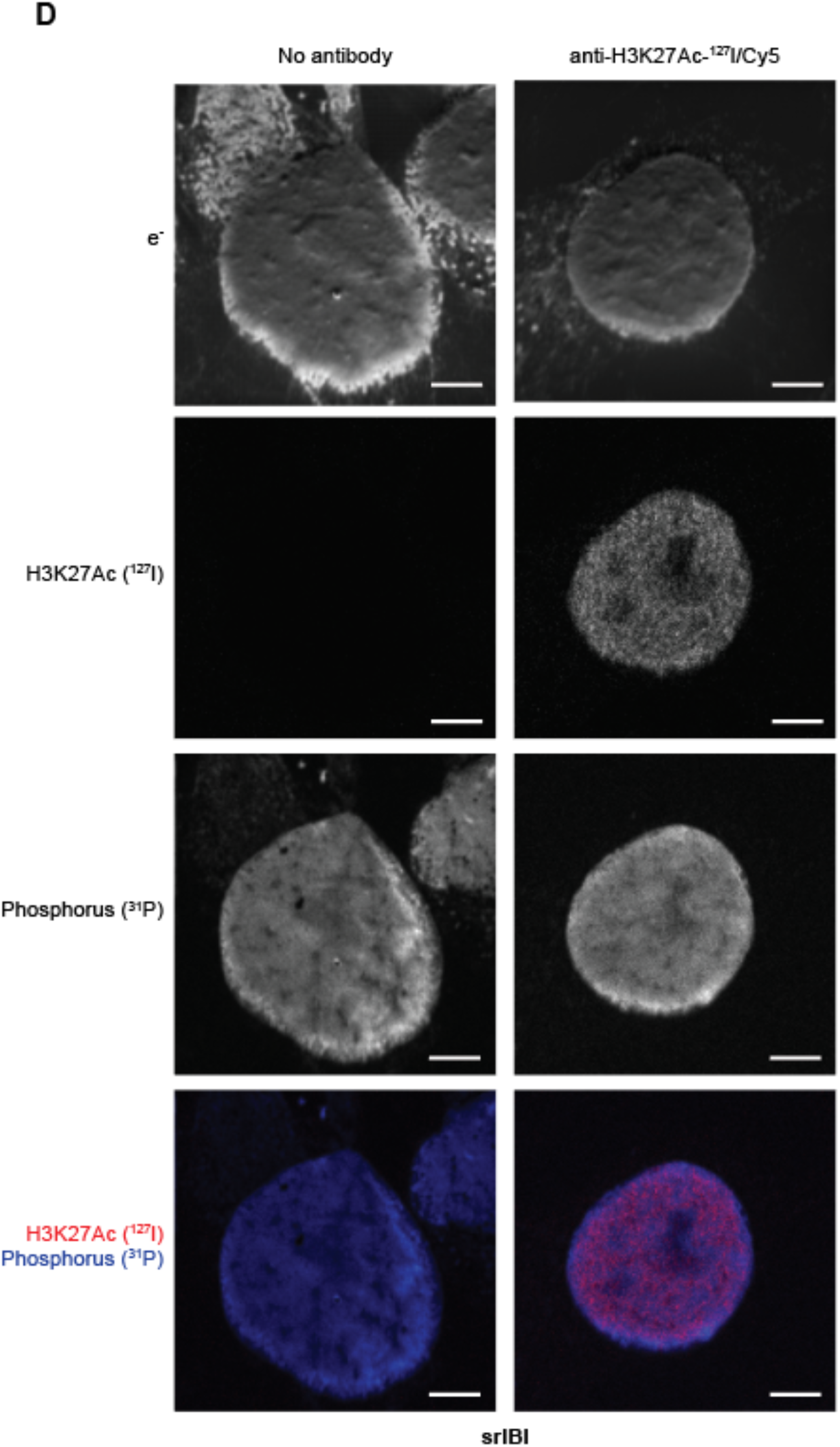
Validation of ^127^I/Cy5 MoC-Abs for srIBI. (**A**) A schematic of the ^127^I/Cy5 MoC-Ab. The antibody was conjugated to the 5’ end of an oligonucleotide in which all cytidines were replaced by 5-I-dC. The oligonucleotide also had a 3’ Cy5. (**B**) Representative confocal microscopy image of a HeLa cell stained with unconjugated anti-H3K27Ac and a secondary anti-mouse-Alexa488 (green) and DAPI (blue). Scale bar, 4 µm. (**C**) Representative confocal microscopy images of control HeLa cells (no antibody) and HeLa cells stained with anti-H3K27Ac-^127^I/Cy5 (red) at three different concentrations. ^31^P is shown in blue. Cells in red boxes in the composite image are enlarged in the right column. Scale bars, 20 µm in standard images and 4 µm in enlarged images. A 1:100 dilution of this antibody was used in subsequent experiments. (**D**) Representative srIBI images of control HeLa cell (no antibody) and HeLa cell stained with anti-H3K27Ac-^127^I/Cy5. Scale bar, 4 µm.

**Figure S9.**
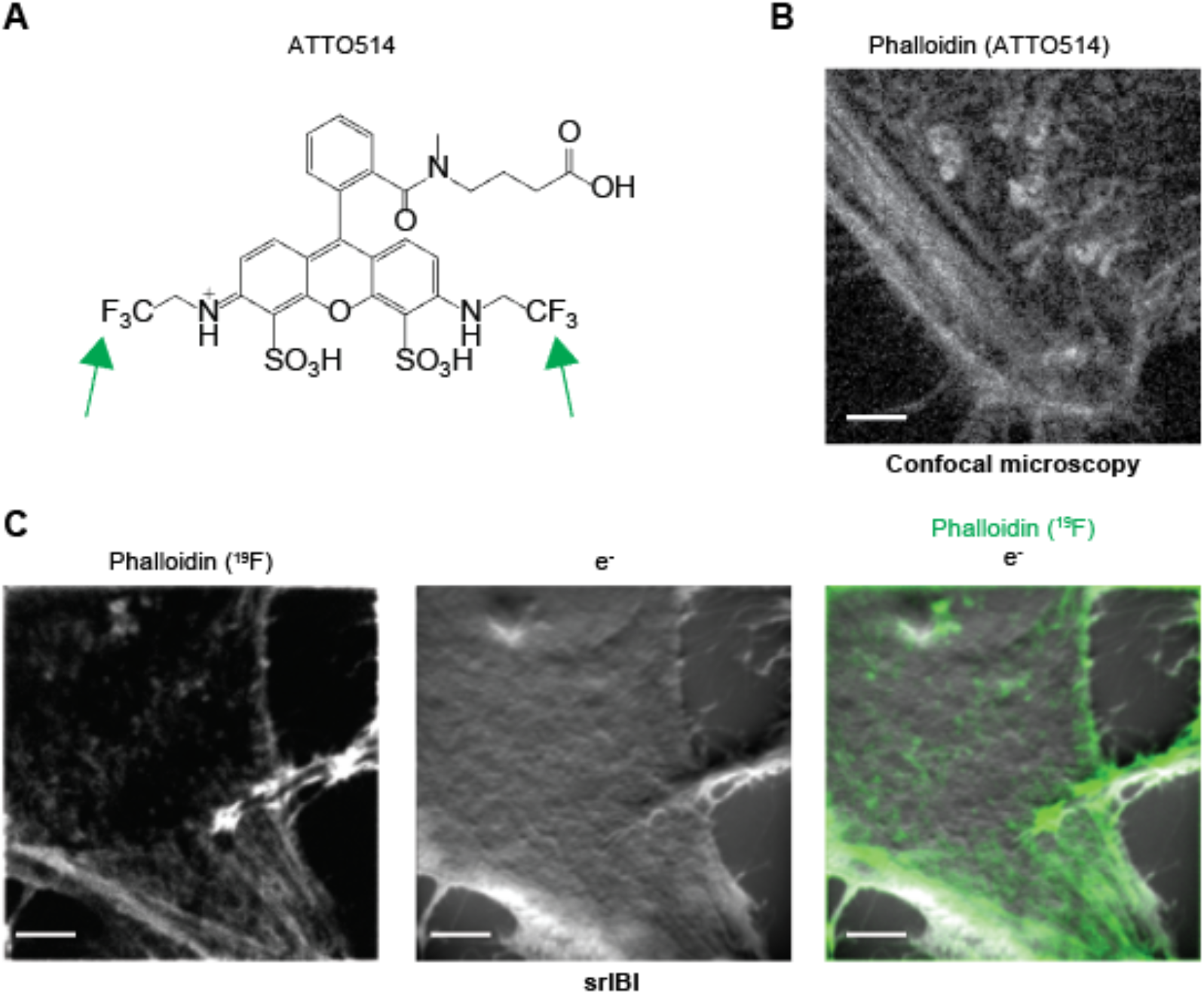
Validation of Phalloidin-ATTO514 for srIBI. (**A**) Chemical structure of ATTO 514. Green arrows indicate the presence of the six ^19^F atoms in ATTO 514. (**B**) Representative confocal microscopy image of a HeLa cell stained with phalloidin-ATTO514. Scale bars, 4 µm. (**C**) Representative srIBI images of a HeLa cell stained with phalloidin-ATTO514. Scale bars, 4 µm.

**Figure S10.**
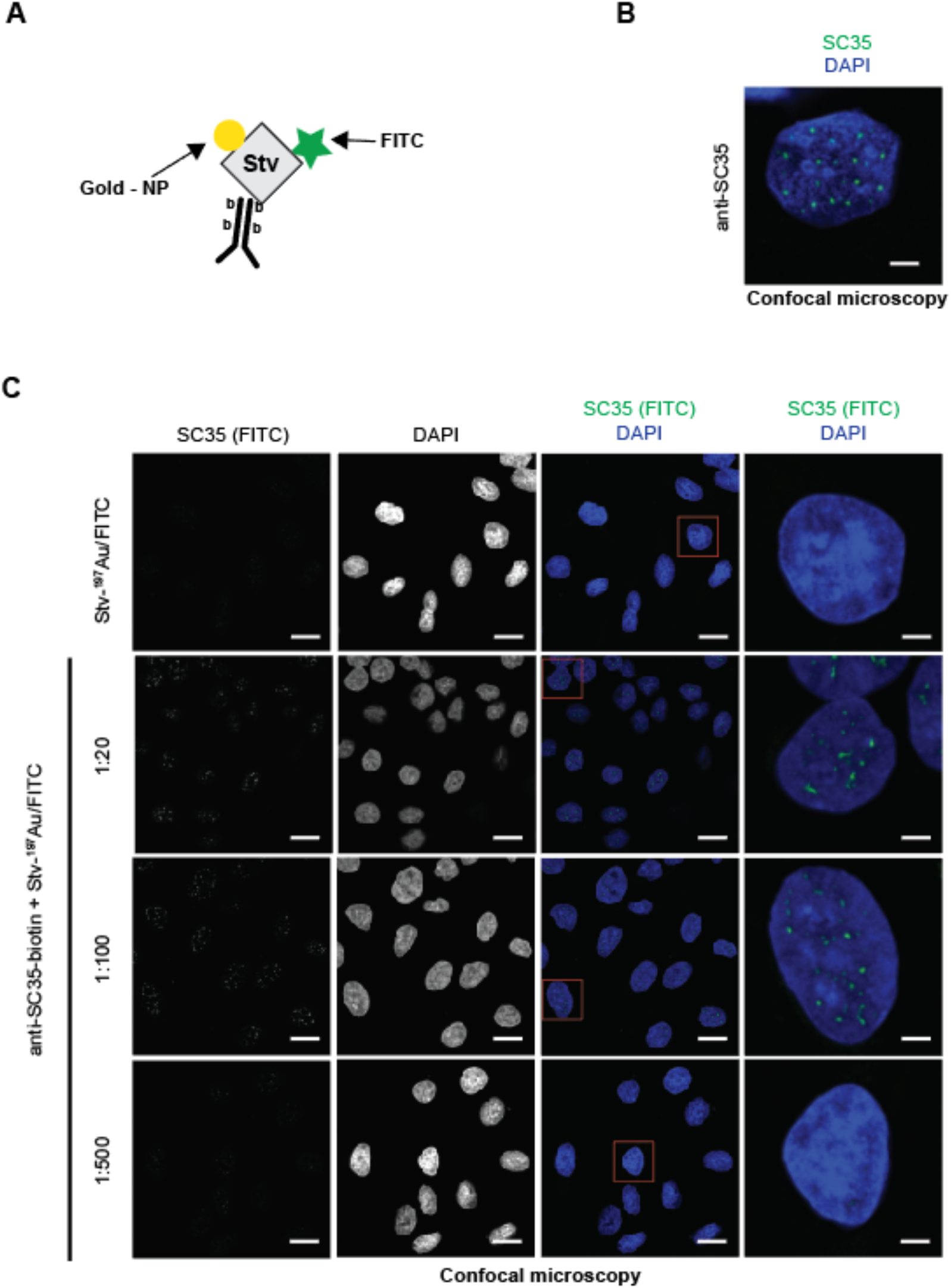
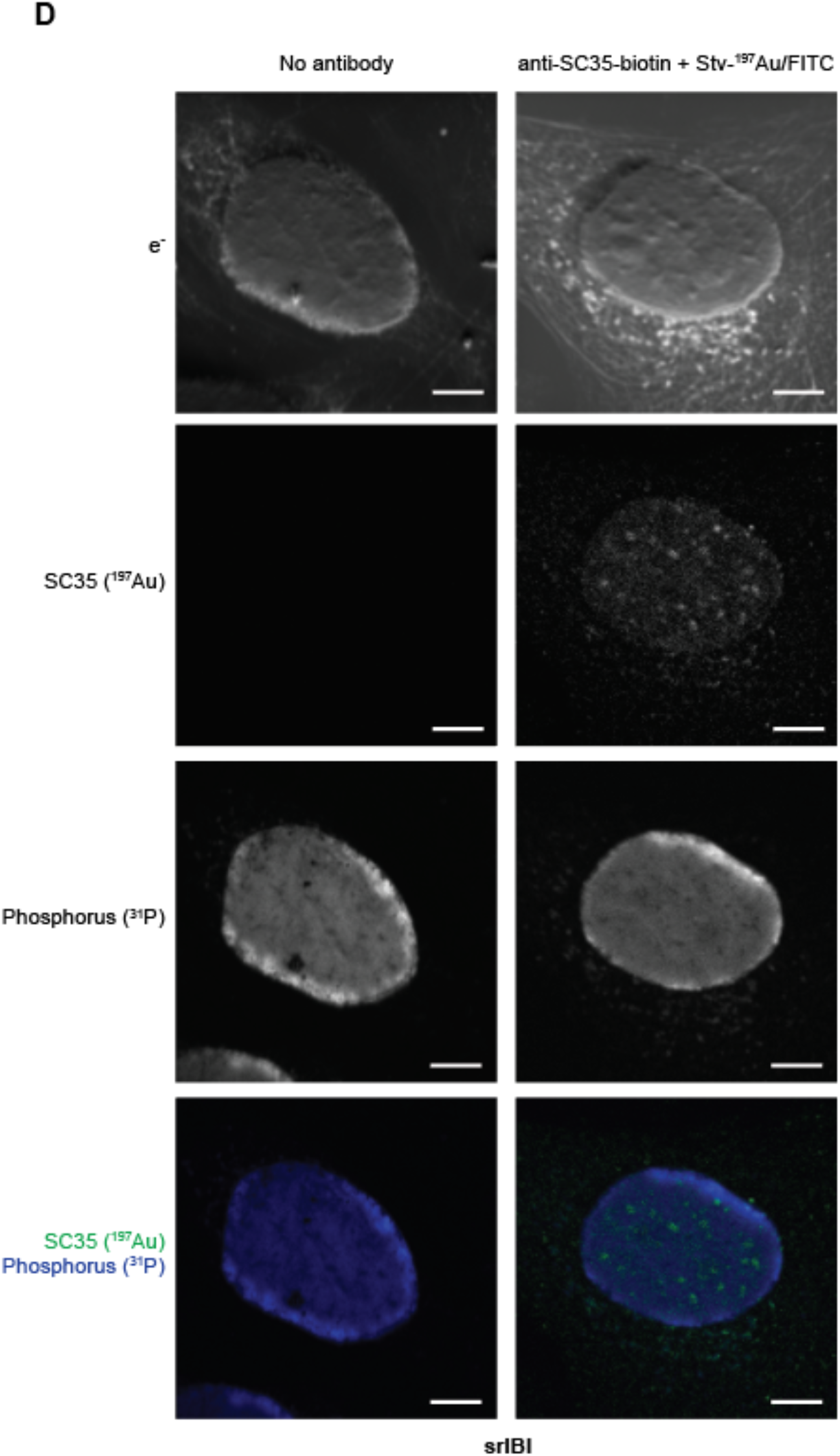
Validation of biotin-conjugated antibodies stained with ^197^Au/FITC-conjugated streptavidin. (**A**) A schematic of the antibody conjugation strategy. Primary antibodies were conjugated to biotin and subsequently stained with ^197^Au/FITC-conjugated streptavidin. The ^197^Au consists of 1.4 nm nanoparticles, a diameter below the theoretical srIBI resolution limit. (**B**) Representative confocal microscopy image of a HeLa cell stained with unconjugated anti-SC35 and a secondary anti-mouse-Alexa488 (green). DAPI (blue) was used to stain the nucleus. Scale bars, 4 µm. (**C**) Representative confocal microscopy images of HeLa cells stained with streptavidin-^197^Au/FITC and HeLa cells stained with anti-SC35-biotin followed by streptavidin-^197^Au/FITC at different concentrations of the primary antibody and 1:40 of streptavidin-^197^Au/FITC (green). DAPI (blue) was used to stain nuclei. Scale bars, 20 µm. Cells in red boxes in the composite image are magnified in the right column. Scale bars, 4 µm. A 1:100 dilution of this antibody was used on following experiments. (**D**) Representative srIBI image of a control HeLa cell (no antibody) and a HeLa cell stained with anti-SC35-biotin followed by streptavidin-^197^Au/FITC. Scale bar, 4 µm.

**Figure S11.**
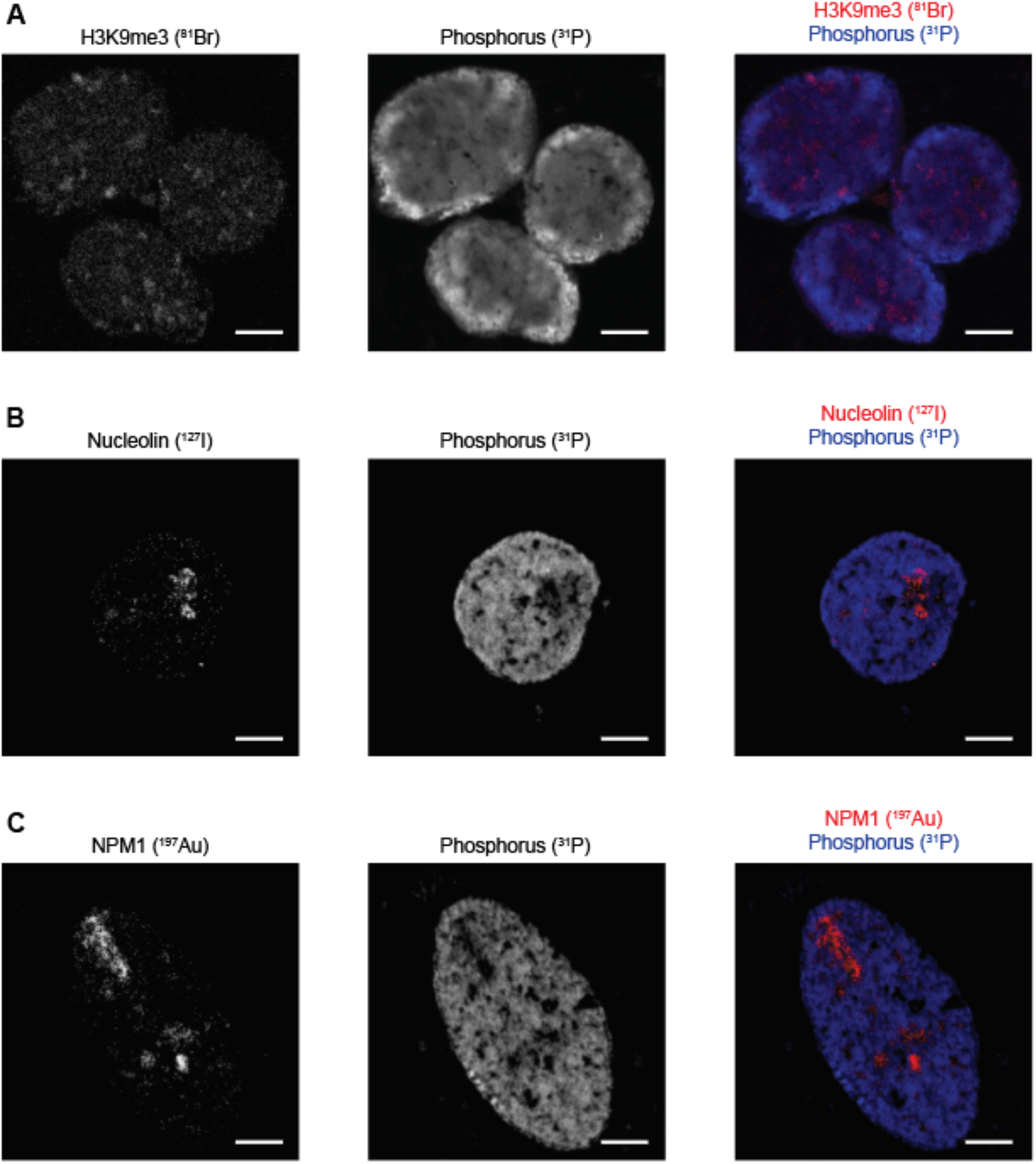
Validation of antibody-based tools for srIBI by targeting other proteins. (**A**) Representative srIBI image of a HeLa cell stained with anti-H3K9me3-^81^Br/Cy3 (red). Phosphorus is shown in blue. Scale bar, 4 µm. (**B**) Representative srIBI image of a HeLa cell stained withanti-nucleolin-^127^I/Cy5 (red). Phosphorus is shown in blue. Scale bar, 4 µm. (**C**) Representative srIBI image of a HeLa cell stained with anti-NPM1-biotin followed by streptavidin-^197^Au/FITC (red). Phosphorus is shown in blue. Scale bars, 4 µm.

**Figure S12.**
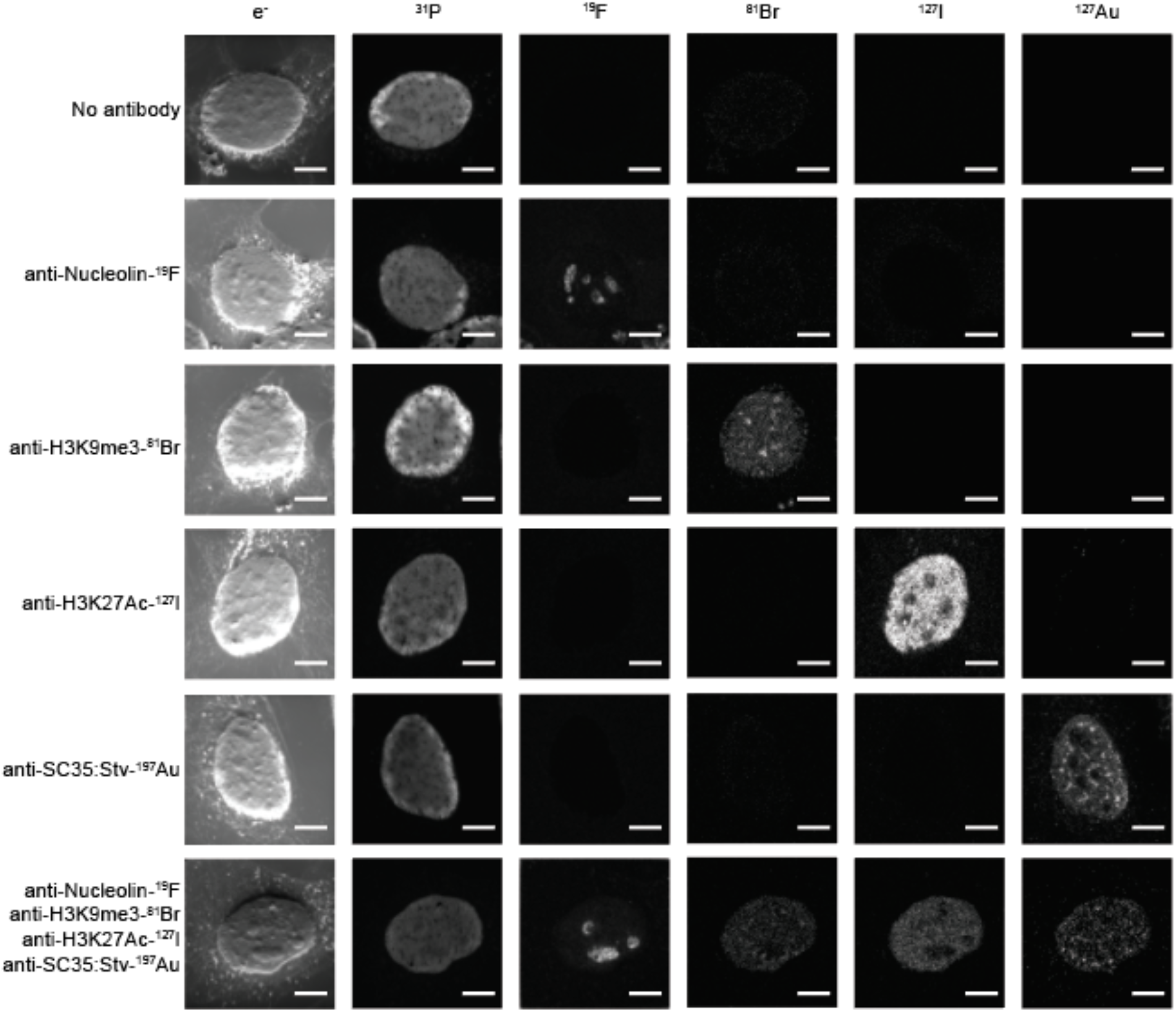
Assessment of srIBI channel crosstalk. Representative srIBI images of a control HeLa cell control (no antibody) and HeLa cells stained with the indicated antibodies. Each row represents a staining condition and each column the ion channel extracted for the indicated isotope. Scale bars, 4 µm.

**Figure S13.**
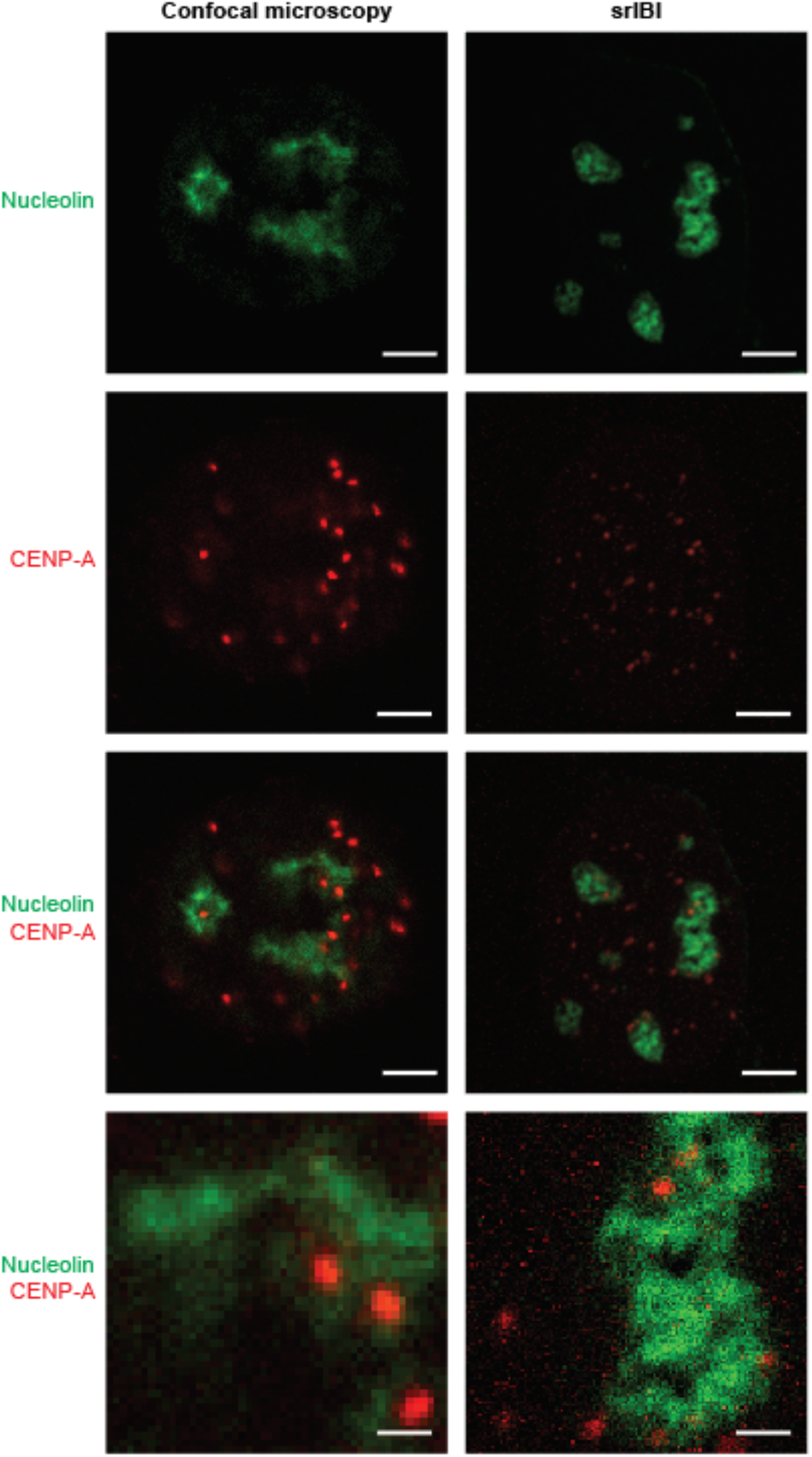
Two-channel srIBI on HeLa cells using a beam diameter of ∼50 nm (D1: 5). Representative confocal microscopy (left) and srIBI (right) images of HeLa cells stained with anti-nucleolin-^19^F/FITC and anti-CENP-A-^81^Br/Cy3 MoC-Ab. Scale bars, 4 µm (top 3 rows) and 1 µm (bottom row).

**Figure S14.**
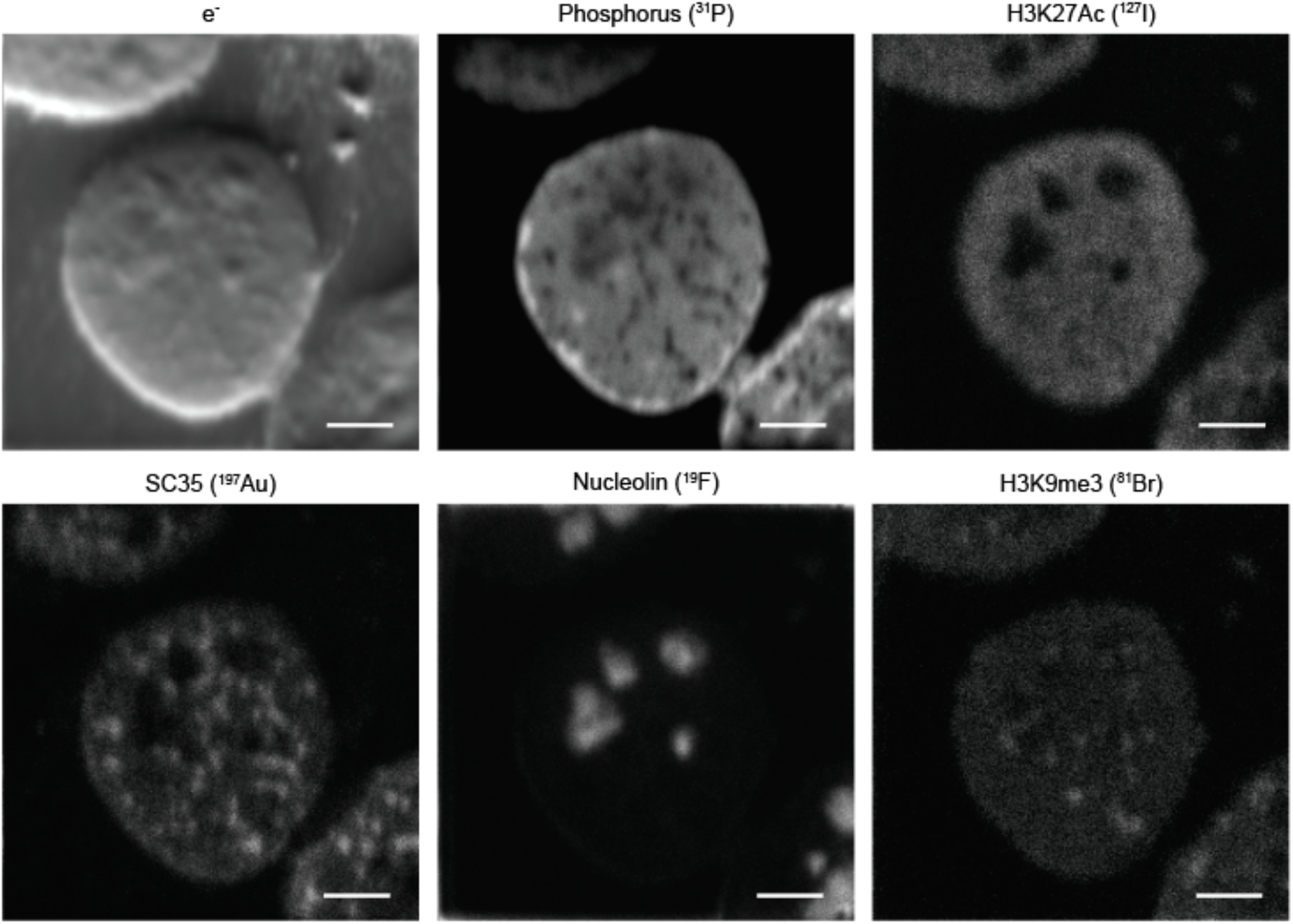
Six-channel srIBI on HeLa cells using a beam diameter of ∼50 nm (D1: 5). Representative srIBI image of a HeLa cell stained with anti-nucleolin-^19^F/FITC, anti-H3K9me3-^81^Br/Cy3, anti-H3K27Ac-^127^I/Cy5, and anti-SC35-biotin (recognized by streptavidin-^197^Au/FITC). Images of the e^-^, nucleolin (^19^F), DNA (^31^P), H3K9me3 (^81^Br), H3K27Ac (^127^I), and SC35 (^197^Au) were simultaneously acquired. A composite image is shown in Fig. 2D. Scale bars, 5 µm.

**Figure S15.**
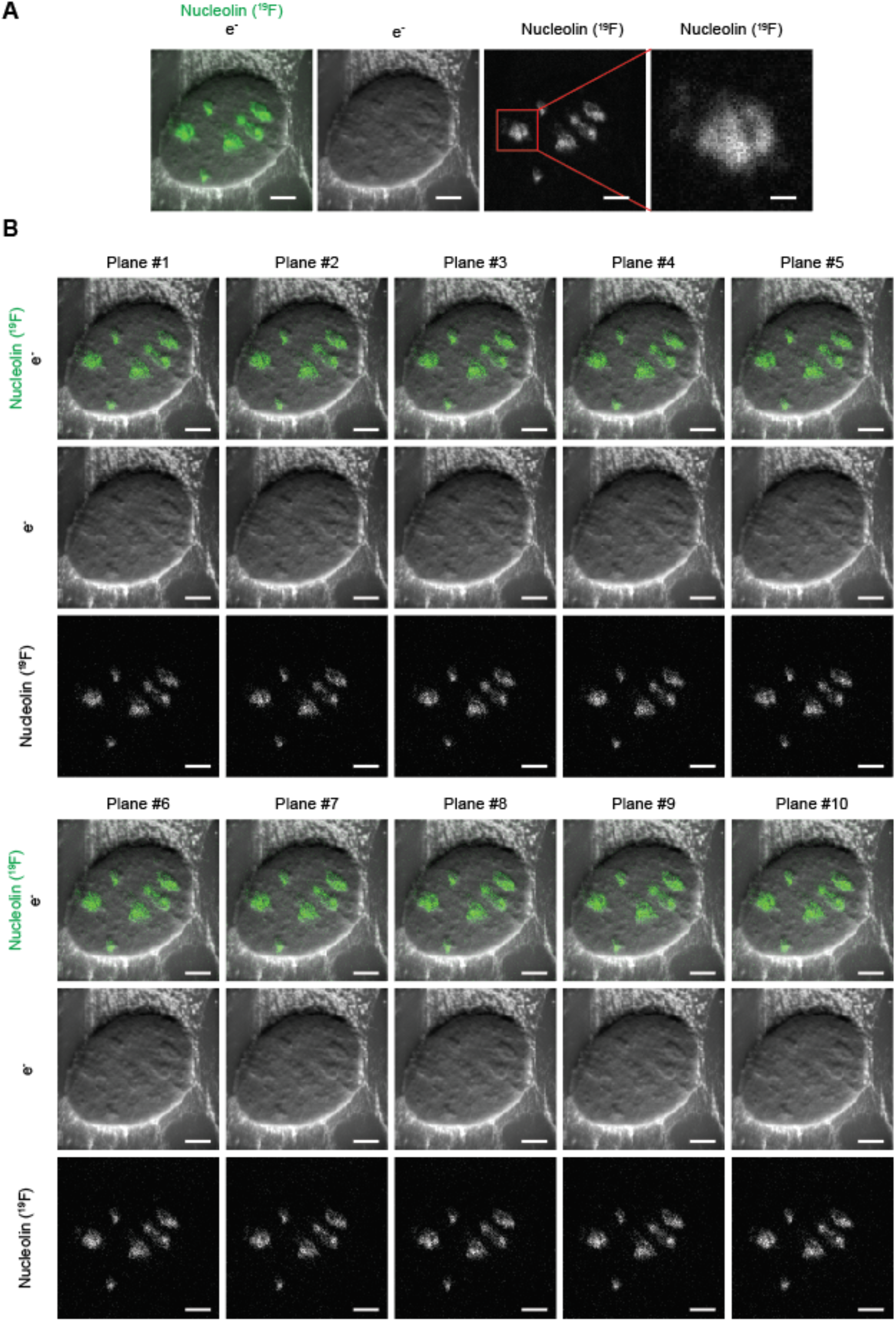
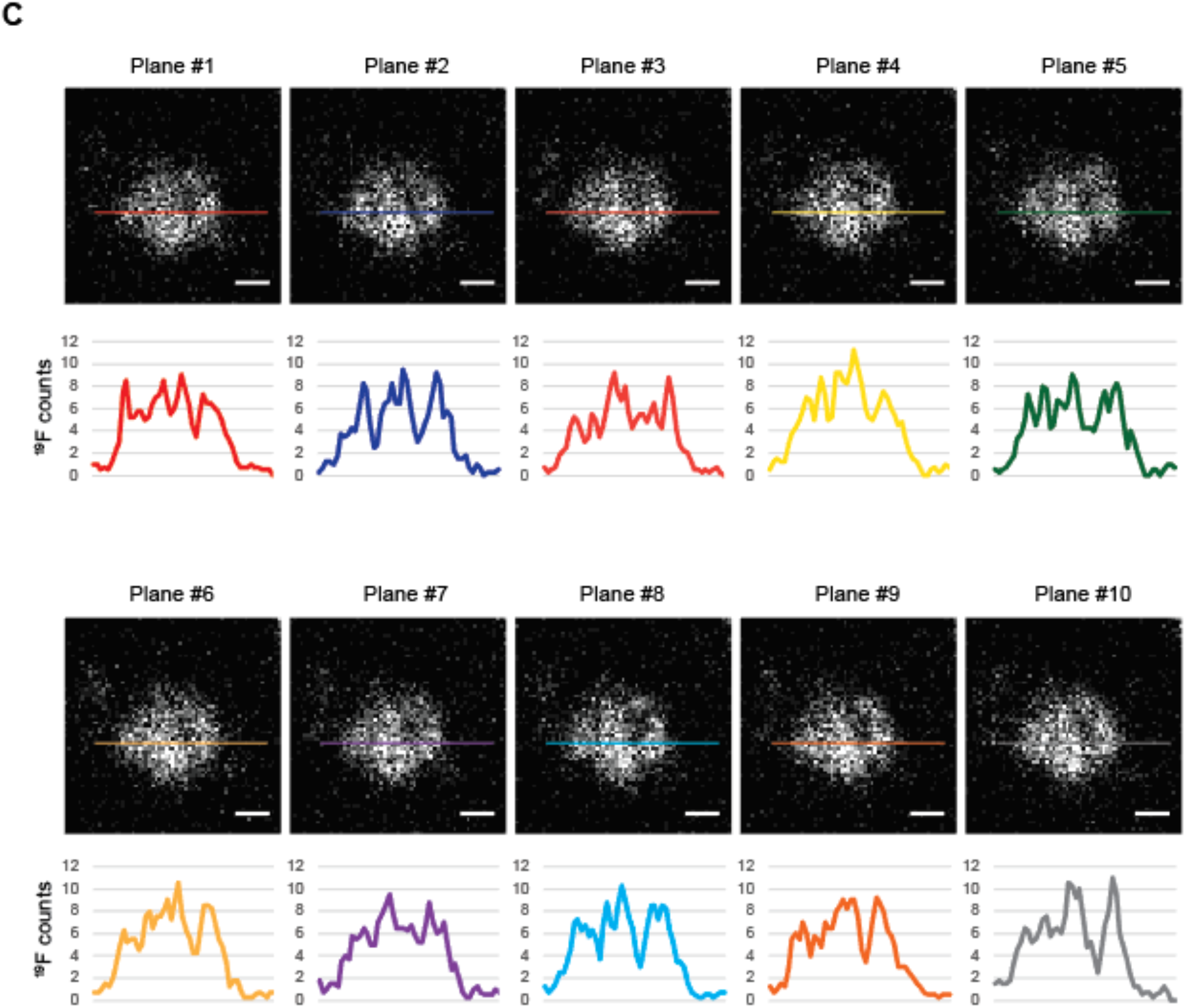
Sequential planes are similar in ion counts. (**A**) Representative srIBI image of a HeLa cell stained with anti-nucleolin-^19^F/FITC. Scale bars, 4 µm; and 1 µm for enlarged images. (**B**) srIBI images of the 10 individual planes summed in panel A in consecutive order. The ^19^F signal is similar in all individual images. Scale bars, 4 µm (**C**) Line scans on the nucleolus enlarged in panel A for each individual plane. Line scans were used to quantify ion counts per pixel in individual planes. Scale bars, 1 µm.

**Figure S16.**
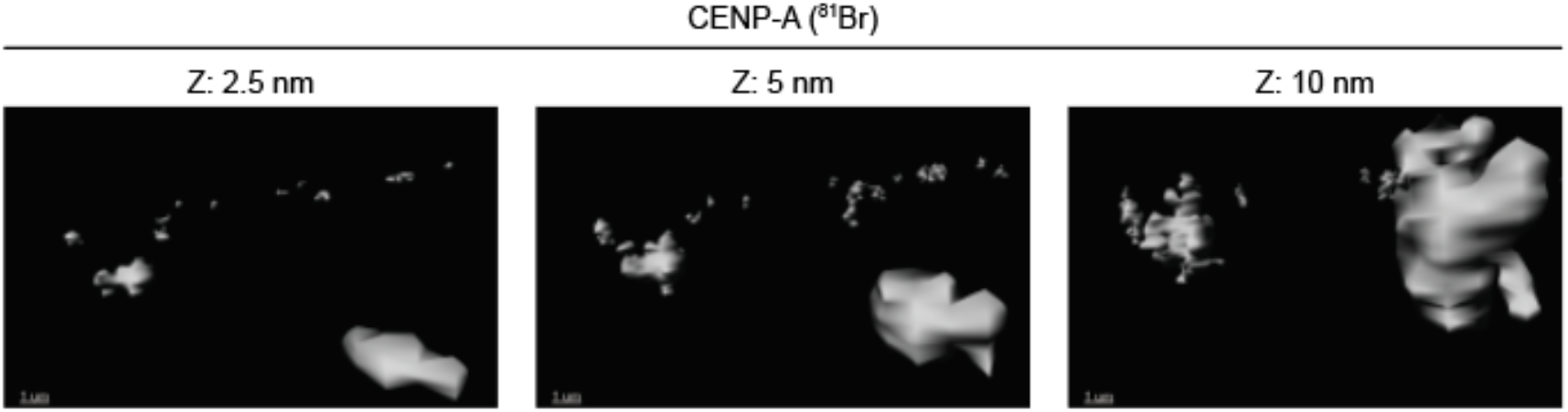
3D surface reconstruction of CENP-A signal at 5 nm Z-depth shows centromeres with a spherical shape. HeLa cells were stained with anti-CENP-A-^81^Br/Cy3, and 40 individual planes were acquired to obtain srIBI images of centromeres from its appearance to its disappearance. 3D surface reconstruction of images of CENP-A at 5 nm reveal centromeres with spherical shape.

**Figure S17.**
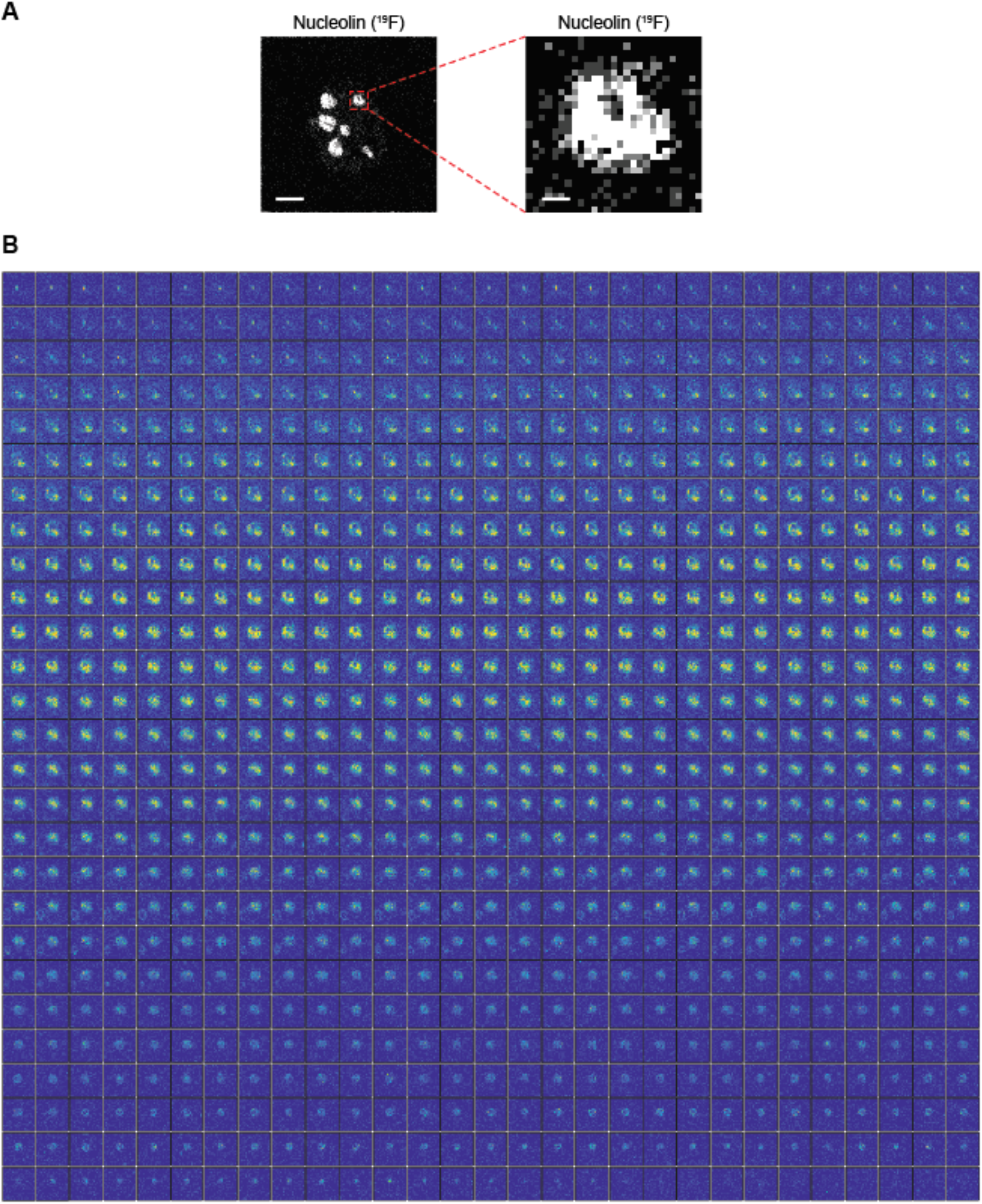
Individual planes of a whole nucleolus. (**A**) (Left) Representative srIBI image of a HeLa cell stained with anti-nucleolin-^19^F/FITC. Scale bar, 4 µm. (Right) Enlarged image of the nucleolus shown in Fig. 3A-B. Scale bar, 400 mm. (**B**) Individual planes of a whole nucleolus, in order from left to right and from top to bottom; these 783 images were used for the volumetric reconstruction of the nucleolus shown in Fig. 3B. Images have the same scale for intensity, and sequential planes have similar ^19^F ion counts.

**Figure S18.**
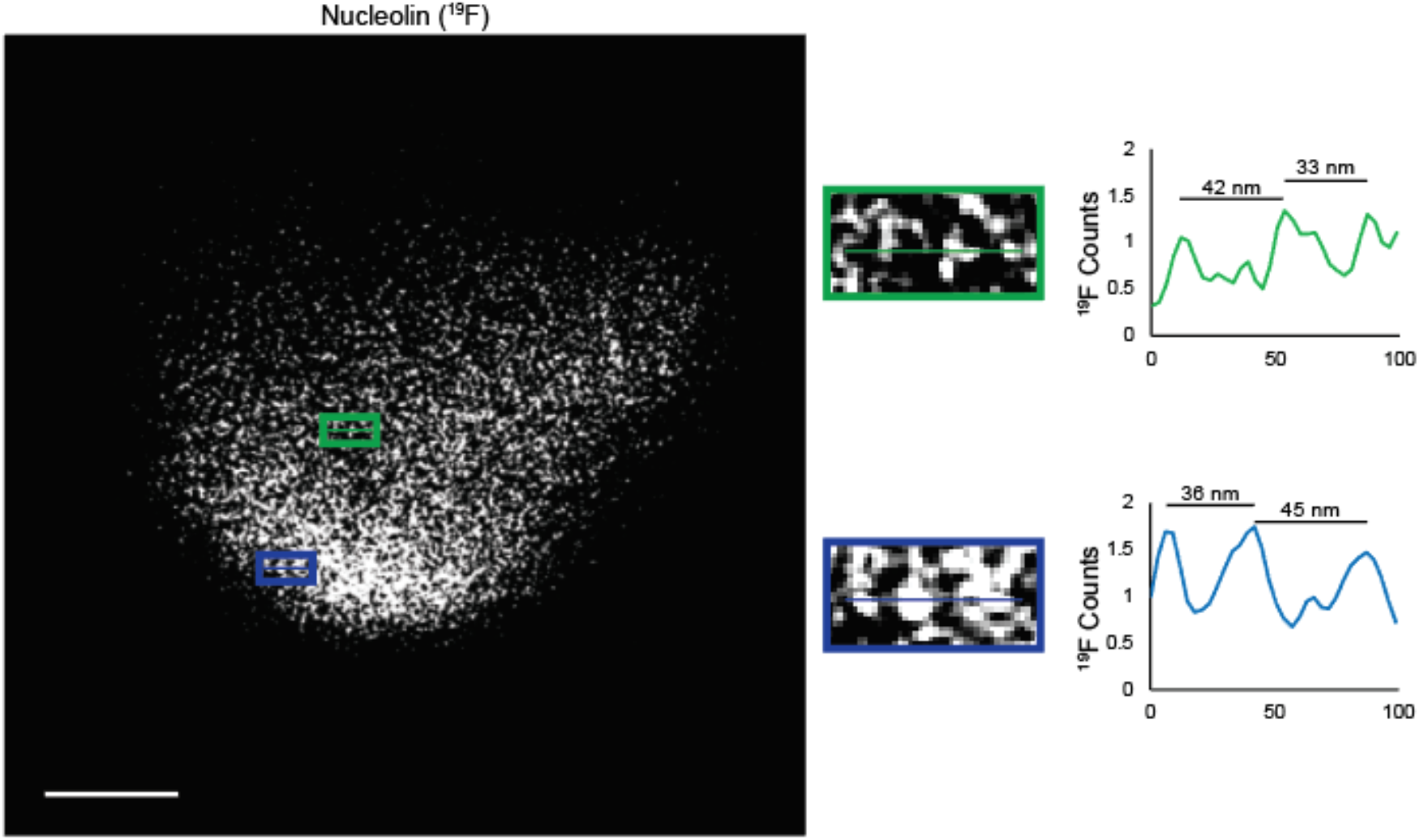
Iterative srIBI is able to resolve MoC-Ab signals at the nanoscale. (Left) Representative enlarged image of a nucleolus of a HeLa cell stained with anti-nucleolin-^19^F/FITC. The same nucleolus is shown in Fig. 3F. Scale bar, 300 nm. (Middle) Higher magnification images of the area boxed in the image on the left. (Right) Line scans of the lines in the middle images demonstrate that srIBI can resolve signals spaced a few tens of nm.

**Figure S19.**
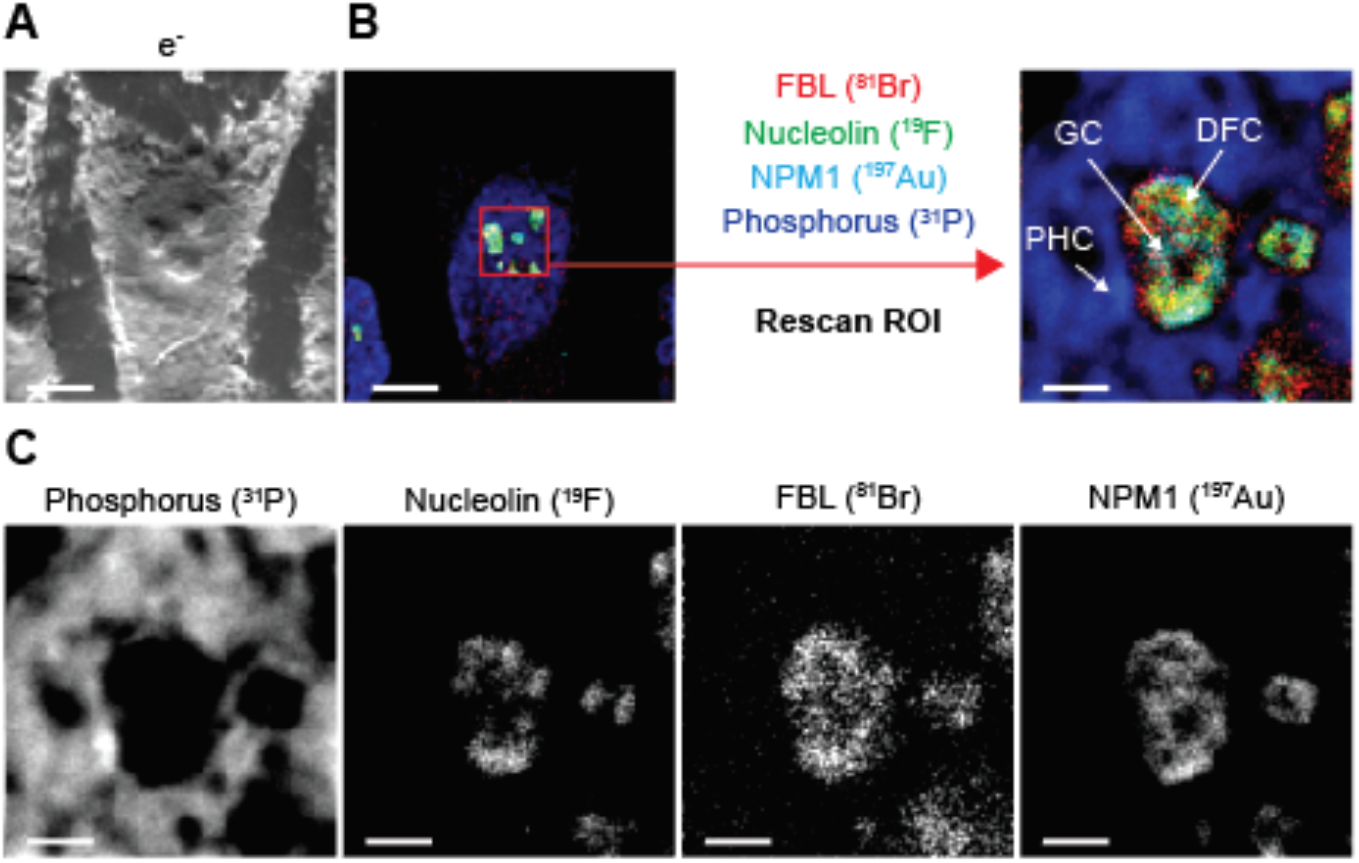
Iterative super-resolution imaging reveals nucleolar substructures. (**A**) Representative secondary electron image of a HeLa cell. Scale bar, 5 µm. (**B**) (Left) High-resolution scan of the HeLa cell used to find a ROI. The HeLa cell shown in panel A was stained with anti-nucleolin-^19^F/FITC, anti-FIB-1-^81^Br/Cy3, and anti-NPM1-biotin (recognized by streptavidin-^197^Au/FITC). Scale bar, 5 µm. (Right) Iterative super-resolution imaging for the visualization of the nucleolar structure. NPM1, a marker of the granular component (GC), is distributed in regions distinct from those stained with FIB-1, a marker of the dense fibrillary component (DFC). Regions with high phosphorus surrounding the nucleolus indicate perinucleolar heterochromatin (PHC). Scale bar, 1 µm. (**C**) Individual images of the super-resolution image shown in panel B. Scale bars, 1 µm.

**Figure S20.**
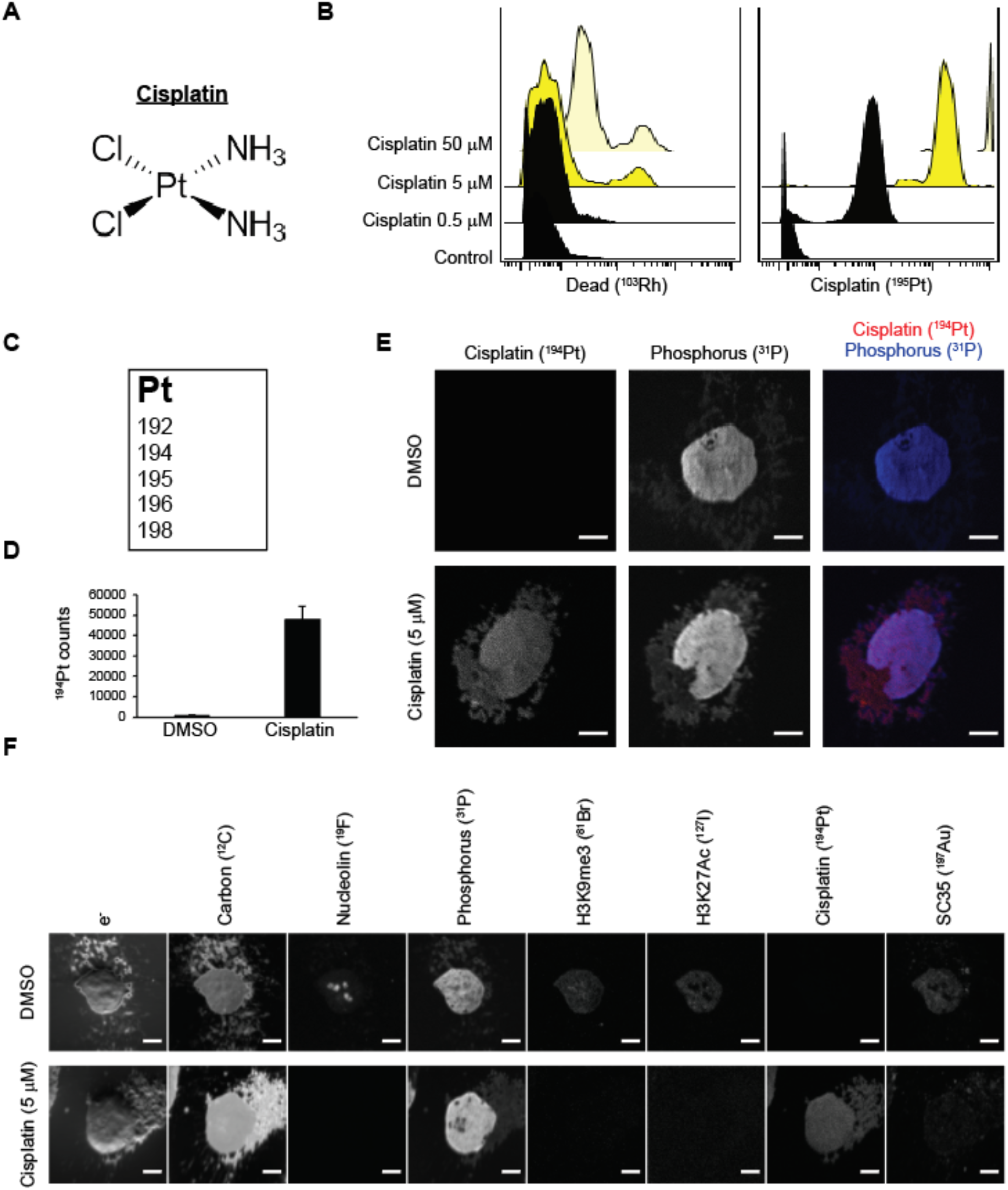
Validation of cisplatin for srIBI. (**A**) Chemical structure of cisplatin. (**B**) CyTOF analysis of cisplatin distribution in TYK-nu ovarian cancer cells. TYK-nu cells were treated with different concentrations of cisplatin for 24 hours, washed, and treated with 1 µM Rh-intercalator for 15 minutes (to discriminate dead from live cells) and analyzed by CyTOF. Maximal drug uptake with minimal cell death at 24 hours was observed with 5 µM. (**C**) The five naturally occurring stable isotopes of cisplatin. (**D**) Mean of ion count per pixel of all pixels in srIBI images from TYK-nu cells treated with cisplatin. TYK-nu cells were treated with DMSO (n=3) or 5 µM cisplatin (n=9) for 24 hours and analyzed by srIBI. Minimal signal was observed in DMSO-treated cells in the ^194^Pt channel. Scale bars, 4 µm. (**E**) Cisplatin is observed in both the cytoplasm and nucleus. Representative srIBI images of a TYK-nu cell treated with DMSO (top) or 5 µM cisplatin for 24 hours without antibody staining (bottom). Scale bars, 4 µm. (**F**) Assessment of ^194^Pt channel crosstalk. (Top) Representative srIBI image of a TYK-nu cell treated with DMSO and stained with anti-nucleolin-^19^F, anti-H3K9me3-^81^Br, anti-H3K27Ac-^127^I, and anti-SC35-biotin (recognized by streptavidin-^197^Au). (Bottom) Representative srIBI image of a TYK-nu cell treated with 5 µM cisplatin for 24 hours without antibody staining. Each row represents a staining condition and each column the ion map for the indicated isotope. Scale bars, 4 µm.

**Figure S21.**
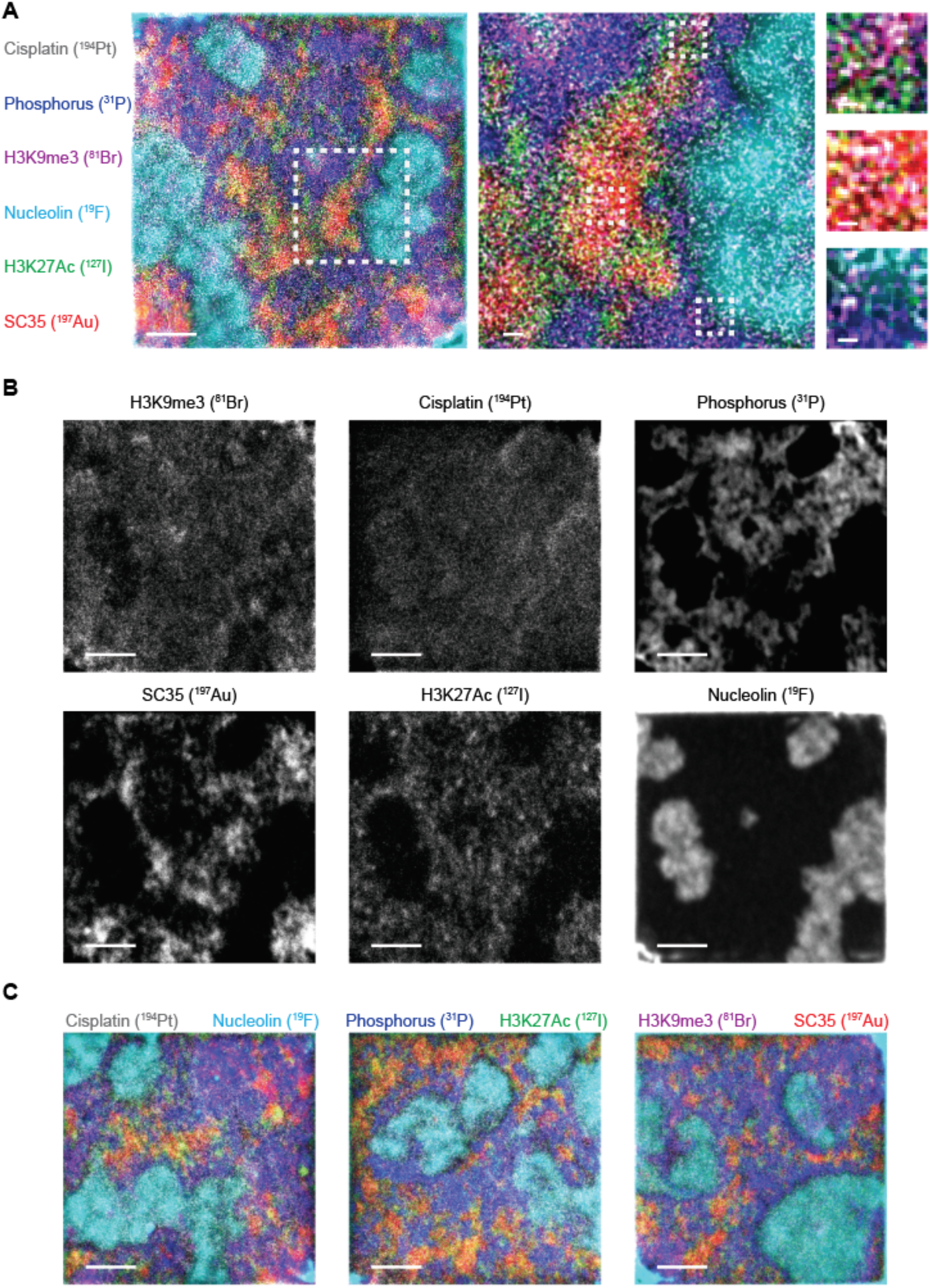
Six-channel srIBI images on TYK-nu cells treated with cisplatin. (**A**-**C**) Representative srIBI images of TYK-nu cells treated with 5 µM cisplatin for 24 hours and stained with anti-nucleolin-^19^F/FITC, anti-H3K9me3-^81^Br/Cy3, anti-H3K27Ac-^127^I/Cy5, and anti-SC35-biotin (recognized by streptavidin-^197^Au). Images of nucleolin (^19^F; cyan), DNA (^31^P; blue), H3K9me3 (^81^Br; magenta), H3K27Ac (^127^I; green), cisplatin (^194^Pt; grey), and SC35 (^197^Au; red) were simultaneously acquired. (**A**) A composite image of a TYK-nu cell nucleus. Scale bars, 2 µm (right image), 200 nm (middle image) and 67 nm (left images). (**B**) Individual images of the cell shown in panel A. Scale bars, 2 µm. (**C**) Composite images from three additional TYK-nu cell nuclei. Scale bars, 2 µm.

**Figure S22.**
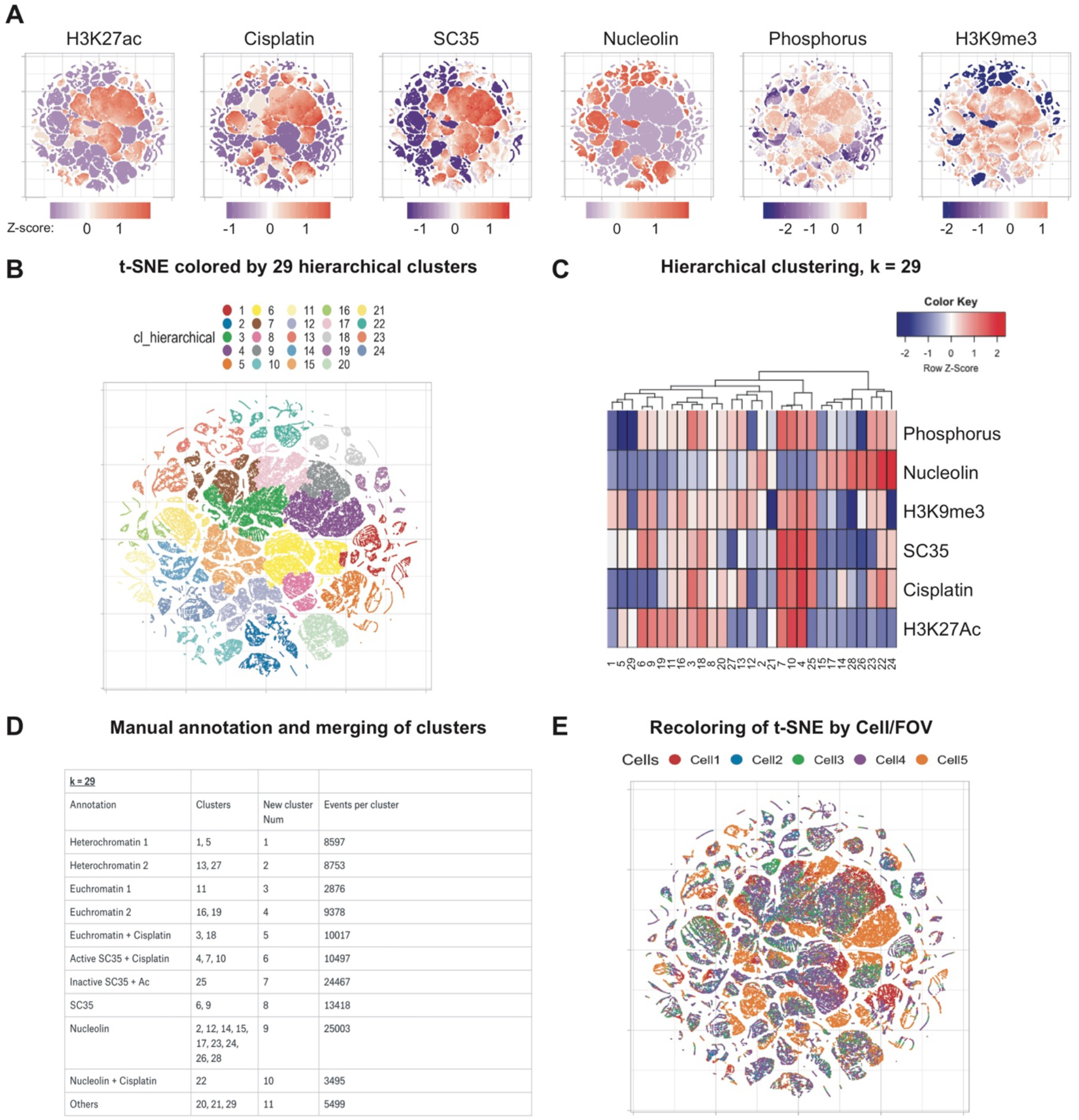
Identification of nuclear neighborhoods by the iterative srIBI analysis framework. (**A**) t-SNE was performed on 100000 voxels of dimensions (x, y, z) = (10, 10, 5) pixels (20000 voxels from each cell were randomly sampled across 5 different cells). Each point represents a voxel, consisting of the expression of six different parameters (phosphorus, nucleolin, H3K9me3, SC35, cisplatin, and H3K27Ac). Voxels grouped into distinct regions based on similar combinatory expression of each marker. The t-SNE map was recolored from left to right to represent the scaled intensity (Z-score) of each indicated marker. (**B**) Unsupervised hierarchical clustering was performed on the voxels to separate them into 29 distinct groups; the t-SNE map from panel A is shown recolored based on these groups. (**C**) A heatmap representation of the 29 distinct clusters identified based on the average expression per cluster of SC35, H3K9me3, phosphorus, nucleolin, H3K27Ac, and cisplatin. The scale intensity of each marker is denoted by the color bar on the top right (Z-score, normalized to each row). (**D**) Clusters were manually merged by tree distance, proximity on the t-SNE map and similar expression profiles. This resulted in a reduction of 29 clusters into 11 clusters, which were then annotated based on their final average expression profile. (**E**) t-SNE map from panel A colored by the cell of origin (from 5 different cells).

**Figure S23.**
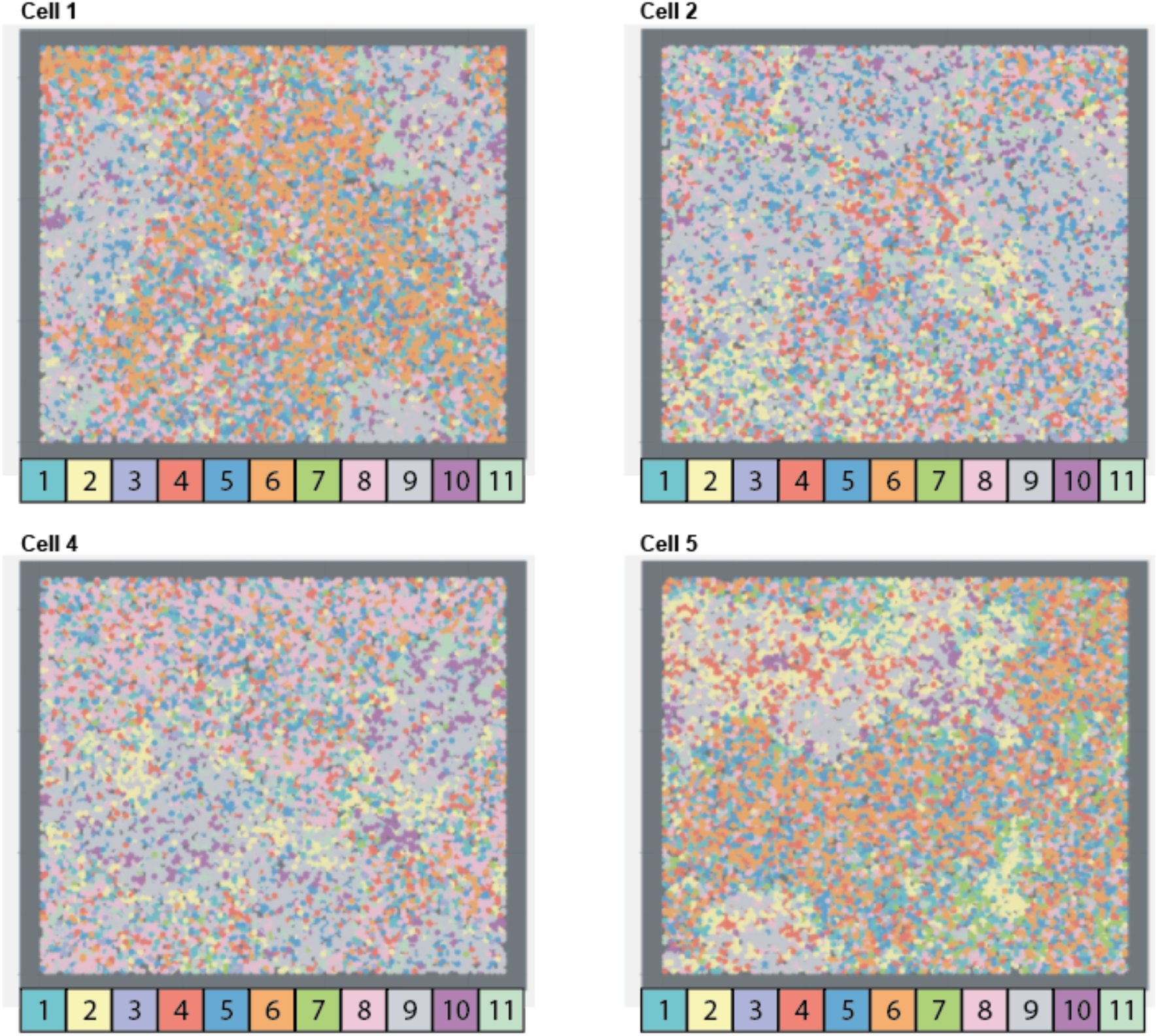
The spatial distribution of nuclear neighborhoods across cells reveals order and diversity in nuclear organization. Nuclear neighborhoods were recolored by each cluster in each of five cells to allow a visual understanding of nuclear neighborhood interactions and conserved and divergent features. Cells 1, 2, 4, and 5 are shown here. Cell 3 is shown in Fig. 5F.

**Figure S24.**
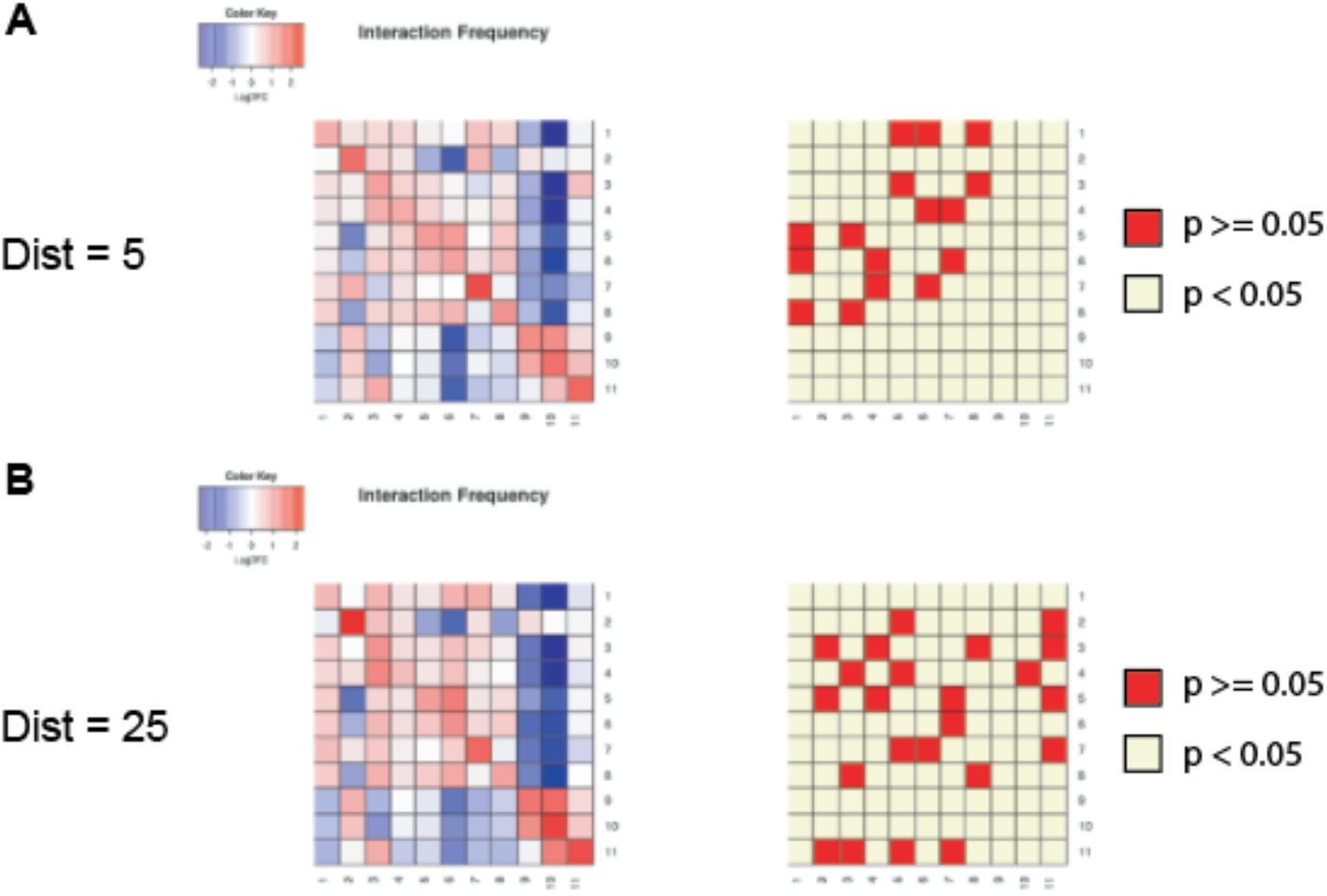
Pairwise neighborhood interaction frequency at different cutoffs. Pairwise voxels were defined as neighbors using a cutoff of *n* pixels in 3D Euclidean space (see Materials and Methods for details). We tested two distances here, using (**A**) 5 pixels and (**B**) 25 pixels as the cutoff. The neighborhood interaction frequency map was calculated using the log2 enrichment of the real over expected number of pairwise interaction frequencies (left). The adjusted p-values are plotted as significant (p< 0.05, cream) or not significant (p >= 0.05, red) (right).

**Figure S25.**
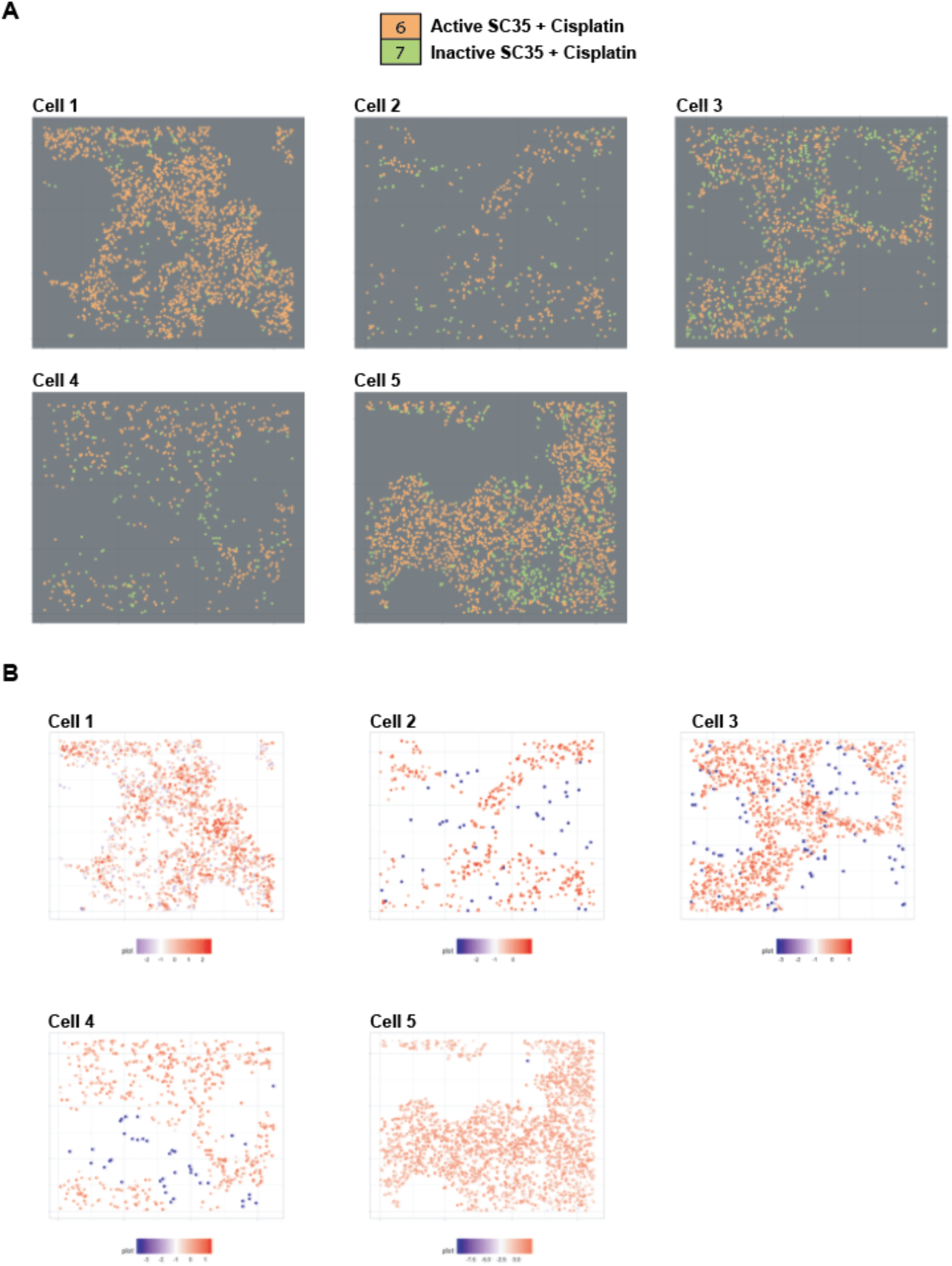
Identification of a cisplatin gradient within SC35 nuclear speckles. (**A**) Neighborhoods 6 (Active SC35 and Cisplatin) and 7 (Inactive SC35 and Cisplatin) were plotted for all 5 cells, to represent the spatial distribution of these two neighborhoods relative to each other. (**B**) Scaled cisplatin levels (Z-score) were plotted on neighborhoods 6 and 7. An increasing gradient of cisplatin concentration is observed across the inactive-active SC35 transition boundary.

**Table S1.** Oligonucleotides.

| Oligo Name | Sequence (5' → 3') | 5' | 3' | Z-Modification |
| --- | --- | --- | --- | --- |
| Det.F.6-FAM | ATAGZAGTZZAGZZGAAZGGTAZAG<br>ATGZZTAAAZAZTTGTZG | Maleimide | 6-FAM | 2'-F-Ac-C |
| Det.Br.Cy3 | ATAGZAGTZZAGZZGAAZGGTAZAG<br>ATGZZTAAAZAZTTGTZG | Maleimide | Cy3 | 5-Br-dC |
| Det.I.Cy5 | ATAGZAGTZZAGZZGAAZGGTAZAG<br>ATGZZTAAAZAZTTGTZG | Maleimide | Cy5 | 5-I-dC |

**Table S2.**
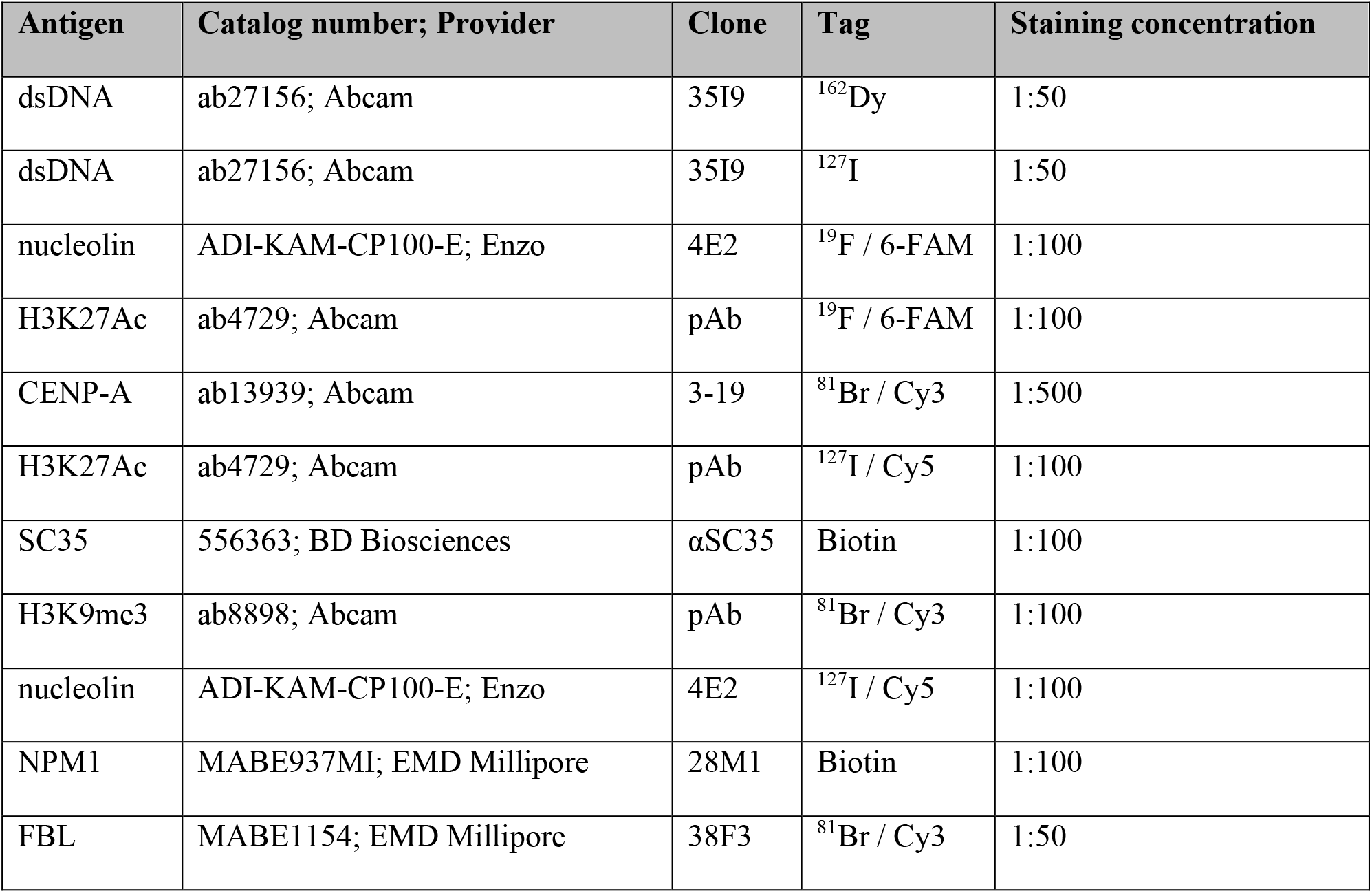
Primary antibodies.

**Table S3.** Buffers for intracellular staining.

| **Buffer** | **Composition** | **Catalog number; Provider** |
| --- | --- | --- |
| Fixation /<br>Permeabilization | 1X Fixation/Permeabilization concentrate<br>in Fixation/Permeabilization diluent | 00-5123; ThermoFisher<br>00-5223; ThermoFisher |
| Wash Buffer | 1X Permeabilization buffer in water | 00-8333; ThermoFisher |
| Block 1 | 1X Permeabilization buffer<br>0.5% BSA<br>0.5 M NaCl<br>200 µg/mL Salmon Sperm DNA<br>10 µg/ml Human FcX Block<br>in avidin buffer | 00-8333; ThermoFisher<br>A3059; Sigma-Aldrich<br>AM9760G; ThermoFisher<br>AM9680; ThermoFisher<br>564765; BD Biosciences<br>927301; BioLegend |
| Block 2 | 1X Permeabilization buffer<br>0.5% BSA<br>0.5 M NaCl<br>200 µg/mL Salmon Sperm DNA<br>10 µg/ml Human FcX Block<br>in biotin buffer | 00-8333; ThermoFisher<br>A3059; Sigma-Aldrich<br>AM9760G; ThermoFisher<br>AM9680; ThermoFisher<br>564765; BD Biosciences<br>927301; BioLegend |
| Reaction | 1X Permeabilization buffer<br>0.5% BSA<br>0.5 M NaCl<br>200 µg/mL Salmon Sperm DNA | 00-8333; ThermoFisher<br>A3059; Sigma-Aldrich<br>AM9760G; ThermoFisher<br>AM9680; ThermoFisher |
|  | in water |  |
| Post-Fixation | 2% Glutaraldehyde in 1X PBS | G7526; Sigma-Aldrich |

**Table S4.** Raster size, pixel number, plane number, and total scan time for each cell.

| **Cell in figure** | **Raster size**<br>**($\mu\text{m}$)** | **Pixel**<br>**number** | **Plane**<br>**number** | **Aperture**<br>**(D1)** | **Total scan**<br>**time (s)** |
| --- | --- | --- | --- | --- | --- |
| 1A | 25 | 512*512 | 160 | 5 | 41966 |
| 1C | 25 | 256*256 | 10 | 3 | 678 |
| 1D | 25 | 256*256 | 10 | 3 | 678 |
| 1E (red dot) | 25 | 512*512 | 10 | 2 | 2644 |
| 1E (orange dot) | 25 | 512*512 | 10 | 3 | 2644 |
| 1E (yellow dot) | 25 | 512*512 | 10 | 4 | 2644 |
| 1E (green dot) | 25 | 512*512 | 10 | 5 | 2644 |
| 2A | 25 | 256*256 | 10 | 3 | 678 |
| 2B | 25 | 256*256 | 10 | 3 | 678 |
| 2C | 25 | 256*256 | 50 | 3 | 3390 |
| 2D-H, J | 25 | 512*512 | 10 | 5 | 2644 |
| 3A, B | 25 | 256*256 | 785 | 3 | 53223 |
| 3C | 25 | 256*256 | 40 | 3 | 2712 |
| 3D | 25 | 256*256 | 400 | 3 | 27120 |
| 3E (top) | 25 | 256*256 | 10 | 3 | 678 |
| 3E (bottom) | 10 | 1024*1024 | 1 | 3 | 1048 |
| 3F | 3 | 1024*1024 | 1 | 5 | 1048 |
| 3G | 10 | 512*512 | 200 | 5 | 52451 |
| 4A, B | 25 | 256*256 | 10 | 3 | 678 |
| 5A | 10 | 1024*1024 | 20 | 5 | 20960 |
| S3D (left) | 25 | 256*256 | 10 | 3 | 678 |
| S3D (right) | 25 | 256*256 | 10 | 3 | 678 |
| S4A | 25 | 256*256 | 100 | 3 | 6780 |
| S5A-C (related to 1D) | 25 | 256*256 | 10 | 3 | 678 |
| S6 (related to 1E) | 25 | 512*512 | 10 | 2 / 3 / 4 / 5 | 2644 |
| S7D (left) | 25 | 256*256 | 10 | 3 | 678 |
| S7D (right) | 25 | 256*256 | 10 | 3 | 678 |
| S8D (left) | 25 | 256*256 | 10 | 3 | 678 |
| S8D (right) | 25 | 256*256 | 10 | 3 | 678 |
| S9C | 25 | 256*256 | 170 | 3 | 11526 |
| S10D (left) | 25 | 256*256 | 10 | 3 | 678 |
| S10D (right) | 25 | 256*256 | 10 | 3 | 678 |
| S11A | 25 | 256*256 | 10 | 3 | 678 |
| S11B | 25 | 256*256 | 20 | 3 | 1356 |
| S11C | 25 | 256*256 | 10 | 3 | 678 |
| S12 (each cell) | 25 | 256*256 | 10 | 3 | 678 |
| S13 (right) | 25 | 512*512 | 40 | 5 | 10576 |
| S14 | 25 | 512*512 | 10 | 5 | 2644 |
| S15 | 25 | 256*256 | 10 | 3 | 678 |
| S16 | 25 | 256*256 | 10 | 3 | 678 |
| S17 (related to 3A-B) | 25 | 256*256 | 785 | 3 | 53223 |
| S18 (related to 3F) | 3 | 1024*1024 | 1 | 5 | 1048 |
| S19 | 5 | 256*256 | 20 | 5 | 1356 |
| S20E (top) | 25 | 256*256 | 10 | 3 | 678 |
| S20E (bottom) | 25 | 256*256 | 10 | 3 | 678 |
| S20F (top) | 25 | 256*256 | 10 | 3 | 678 |
| S20F (bottom) | 25 | 256*256 | 10 | 3 | 678 |
| S21A-B | 10 | 1024*1024 | 21 | 5 | 22043 |
| S21C (each cell; related to 5A) | 10 | 1024*1024 | 20 | 5 | 20960 |
| Movies S1 and S2<br>(related to 3A-B and S17) | 25 | 256*256 | 785 | 3 | 53223 |
| Movie S3 (related to 3D) | 25 | 256*256 | 400 | 3 | 27120 |
| Movie S4 (related to 1A) | 25 | 512*512 | 160 | 5 | 41966 |

**Movie S1. 3D reconstruction of nucleoli from a single cell.**

3D surface reconstruction of the nucleoli from a single HeLa cell. HeLa cells were stained with anti-nucleolin-^19^F/FITC, and 785 individual planes were acquired to obtain srIBI images of a nucleolus from its appearance to its disappearance.

**Movie S2. 3D reconstruction of a single nucleolus.**

3D surface reconstruction of a nucleolus shown in Fig. 3B and Movie S1. HeLa cells were stained with anti-nucleolin-^19^F/FITC, and 785 individual planes were acquired to obtain srIBI images of a nucleolus from its appearance to its disappearance. See Fig. S17 for images of each individual plane.

**Movie S3. 3D reconstruction of a single cell.**

Representative 3D reconstruction of nucleolin (cyan), phosphorus (blue), H3K9me3 (magenta), H3K27Ac (green), and SC35 (red) in a HeLa cell stained with anti-nucleolin-^19^F/FITC, anti-H3K9me3-^81^Br/Cy3, anti-H3K27Ac-^127^I/Cy5, and anti-SC35-biotin (recognized by streptavidin-^197^Au/FITC). The image consists of the 3D reconstruction of a stack of 400 consecutive planes.

**Movie S4. 3D reconstruction of a single cell treated with cisplatin.**

Representative 3D reconstruction of carbon (grey), nucleolin (cyan), phosphorus (blue), H3K9me3 (magenta), H3K27Ac (green), cisplatin (yellow), and SC35 (red) in a TYK-nu cell treated with 5 µM cisplatin for 24 hours and stained with anti-nucleolin-^19^F/FITC, anti-H3K9me3-^81^Br/Cy3, anti-H3K27Ac-^127^I/Cy5, and anti-SC35-biotin (detected with streptavidin-^197^Au/FITC). The image consists of the 3D reconstruction of a stack of 160 consecutive planes.

